# Potential for the Terminal SKP1 Glycosyltransferase to Exert Non-Enzymatic Control of SKP1 in Toxoplasma gondii

**DOI:** 10.64898/2026.06.11.731712

**Authors:** Donovan A. Cantrell, Elizabet Gas-Pascual, Christopher M. West

## Abstract

The SKP1/CUL1/F-Box protein (SCF) family of E3 ubiquitin ligases is responsible for targeting multiple proteins for degradation by the 26S proteosome. Within the family, multiple F-box proteins (FBPs) associate with the SKP1 adaptor which in turn docks to CUL1. In the human parasite *Toxoplasma gondii*, SKP1 is subject to oxygen-dependent regulation. Under normoxic conditions, the prolyl hydroxylase PHYa hydroxylates SKP1 priming it for modification by five SKP1-specific glycosyltransferase activities. Glycosylation weakens competing homodimerization of SKP1, and because SKP1 is limiting in cells, this affects the profile of bound FBPs. A previous study noted that the terminal SKP1 glycosyltransferase, GAT1, is a stable member of the SKP1 interactome regardless of SKP1’s glycosylation status. Furthermore, *gat1*-knockout cells exhibit a unique repertoire of FBPs bound to SKP1 relative to normal and other glycosylation-defective mutants. Utilizing sedimentation velocity analytical ultracentrifugation, we demonstrate that the native GAT1 homodimer complexes with SKP1 monomers with affinity and stoichiometry dictated by its glycostate. Computational modeling validated by mutational probing shows that GAT1 competes with the core hydrophobic interface also utilized by FBPs and the SKP1 homodimer. This interface is complemented by varying, transient fuzzy-like interactions contributed by SKP1’s intrinsically disordered C-terminal region (CTR) that are in turn constrained by the glycan. Furthermore, GAT1 substoichiometrically monomerizes SKP1 in a CTR-dependent manner. Thus, GAT1 is specific for monomeric SKP1, and the mechanism by which this is achieved also modifies the dimer/monomer equilibrium of unmodified SKP1 with downstream effects on the steady-state profile of FBP/SKP1 complexes in cells.

## INTRODUCTION

Toxoplasmosis is a disease of warm-blooded animals caused by exposure to the obligate intracellular protozoan parasite *Toxoplasma gondii*. Infection can occur by ingesting undercooked meat, handling potentially infectious materials such as cat litter, or from receiving an infected person’s blood [1,2]. Toxoplasmosis represents the leading cause of death from foodborne illness within the United States. Acute infection is followed by immune control leading to latency after the parasite transforms into slow growing bradyzoite cysts that are protected from immune clearance or pharmacological treatments [3,4]. However, immunocompromised individuals are at high risk because latent infections can activate with life-threatening or disabling sequelae in the brain and eyes [5]. Due to the inability to clear latent *Toxoplasma* through pharmaceutical means, immunocompromised individuals are forced to undergo treatment for life. In addition, when naïve pregnant women become infected, neonatal toxoplasmosis can lead to debilitating birth defects [6]. Up to a third of the world’s population is thought to be latently infected and this proportion approaches 80% in some western countries [2]. New knowledge about the biochemistry of *Toxoplasma* regulation is expected to expose new opportunities to develop cures.

To effectively infect new hosts, home into preferred host tissues, and differentiate appropriately, the parasite monitors its chemical environment. One varying factor is the concentration of dissolved O_2_. As recently summarized [7], *Toxoplasma* possesses the prolyl hydroxylase PHYa, which is a likely ortholog of human PHD2 that regulates Hypoxia Inducible-1α (HIF1α) in animals and acts as a major oxygen sensing enzyme [8]. In an oral mouse infection model using a *Toxoplasma* mutant deleted for *phyA*, tissue homing and parasite virulence is impaired [9]. PHYa acts by hydroxylating a specific Pro residue conserved within protist SKP1 (P154 for *Toxoplasma*) allowing for the attachment of a pentasaccharide (Fig. 1A). A primary if not sole role of SKP1 is as the adaptor subunit of the SKP1/CUL1/F-box (SCF) complex which is an E3 ubiquitin ligase partially responsible for proteomic control throughout eukaryotes [10,11,12]. SKP1 acts to bridge any of a large variety of substrate binding FBPs, via Subsites A and B, to CUL1 of the SCF complex (Fig. 1B). Upon substrate recognition by the appropriate FBP and its attachment to the SCF complex the bound protein target can be ubiquitinated typically at a target Lys. Repetitive action of the ligase generates linear chains of Ub linked to K48 of the underlying Ub, resulting in recognition of the target and its subsequent degradation by the 26S proteosome.

**Figure 1.**
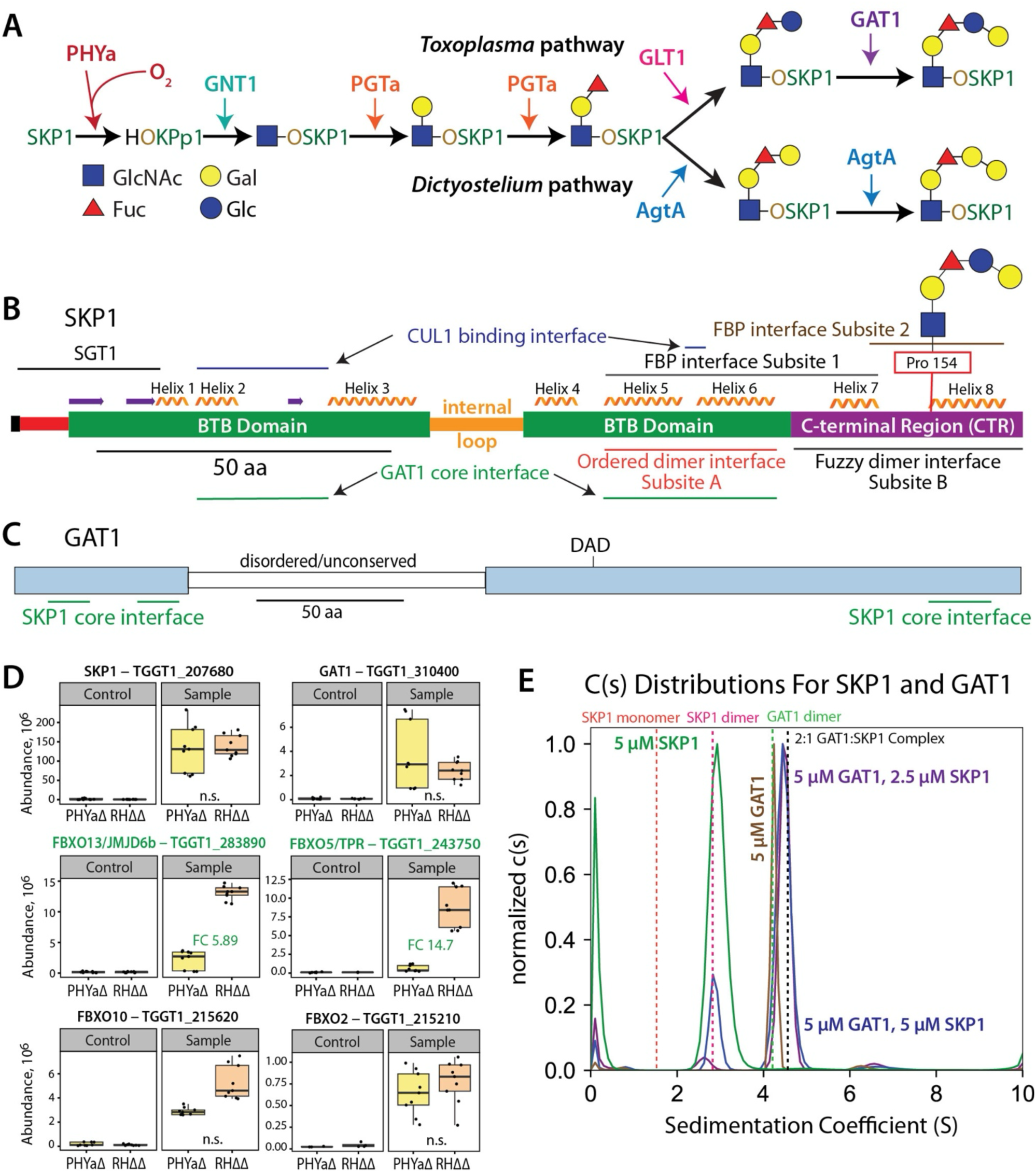
GAT1’s Role in SKP1 Glycosylation and Evidence for GAT1:SKP1 Interactions. (A) Comparison of oxygen dependent glycosylation pathways in *Toxoplasma* and *Dictyostelium*. Modifications and responsible enzyme activities are indicated. (B) A domain diagram of SKP1 indicates the N-terminal disordered region (red), folded BTB domain (green), a disordered internal loop (orange), and a conditionally disordered/ordered C-terminal Region (purple). The positions of secondary structural elements are indicated. α-helixes 5, 6, and 7 constitute SKP1’s F-Box binding Subsite-1 while α-helix 8 represents SKP1’s variable F-box binding Subsite-2. SKP1 hydroxylation and glycosylation occur at the N-terminus of α-helix 8 which favors a disorder to order transition. In free SKP1 α-helices 5 and 6 are similarly folded but the region of 7 and 8 is partially disordered. These interactions occur distally from binding of other SKP1 binding partners such as Cul1 and possibly Sgt1. As shown here, GAT1’s core hydrophobic interface overlaps with the SKP1 homodimer, SKP1/FBP, and SKP1/Cul1 interfaces. (C) A domain diagram of GAT1 indicates sequence components that are part of the catalytic GT-A fold or comprise a disordered internal loop. The portions of GAT1 which bind SKP1 in the core hydrophobic interface are shown. (D) Co-IP experiments indicate stable binding of GAT1 to SKP1 *in vivo* whether or not SKP1 is modified [from 13], in comparison to levels of other FBPs. RHΔΔ refers to the normal parental strain, and gene names are from VEuPathDB. (E) c(s) distributions demonstrate the sedimentation profiles of 5 µM SKP1 (green), 5 µM GAT1 (brown), 5 µM GAT1 with 5 µM SKP1 (blue), and 5 µM GAT1 with 2.5 µM SKP1 (purple). Dashed lines represent the expected sedimentation of the SKP1 monomer (orange), SKP1 dimer (hot pink), GAT1 dimer in the absence of SKP1 (green), and 2:1 GAT1:SKP1 complex (black). Supporting data can be found in Fig S2.

SKP1 hydroxylation and subsequent glycosylation has been observed to alter SKP1’s FBP binding repertoire [13,14]. Studies that originated in another protist that conserves this modification, the social amoeba *Dictyostelium discoideum* (Fig. 1A), suggested that the altered interactome results from changes in SKP1’s conformational ensemble [15–17]. Glycosylation promoted a disorder to order transition in its partially disordered C-terminal region (Fig. 1B) suggesting a direct effect on SKP1’s FBP binding repertoire. A second effect of glycosylation was weakening of the SKP1 homodimer from an apparent *K*_d_ of ∼50 nM to 400 nM [18]. 50 nM is close to the 25 nM *K*_d_ described for a mammalian FBP/SKP1 interaction [19]. Considering their overlapping interfaces, these interactions may compete with one another in the cell. This high affinity homodimer involves a bipartite interface consisting of an ordered Subsite-A, and a charge-based fuzzy/disordered Subsite-B at the C-terminal region (CTR) that includes the glycosylation site [18]. Subsites A and B partially overlap with Subsites-1 and -2 of the interface with FBPs, rendering homodimerization competitive with FBP-binding (Fig. 1B). Weakening of SKP1’s homodimer by glycosylation is believed to occur through effects on the fuzzy component. Elsewhere, the dynamic nature of fuzzy interactions are proposed to affect exchange with other binding partners and be permissive for posttranslational modifications [20–23]. By weakening this tight competitive interaction, glycosylation may promote interactions with FBPs predicted to have lower affinity [12]. Through the effect of glycosylation on the two interactions, the levels of SCF target substrates are expected to be influenced by available O_2_.

The SKP1 glycosyltransferases (GTs) are distinctive in functioning in the cytoplasmic compartment of cells and evidently dedicated solely to modifying SKP1 [12]. Counterintuitively, the terminal GT in the *Toxoplasma* pathway, GAT1, consistently appears when SKP1 is co-immunoprecipitated (co-IP) from *Toxoplasma* regardless of its glycosylation status (Fig. 1D) [13]. It is unusual for a GT to be bound to non-substrate glycoforms and its product, and other GTs were not detected. Furthermore as shown herein, the profile of FBPs found in the SKP1 interactome was uniquely affected compared to wild-type and *phyA*-*gnt1*- and *pgtA*-KO cells. Therefore, we focused our attention on characterizing the *in vitro* interaction between GAT1 and glycosylated or unmodified isoforms of SKP1 for clues as to the significance of the *in vivo* GAT1/SKP1 interaction. First, we utilized sedimentation velocity analytical ultracentrifugation (SV-AUC) to confirm the formation of a direct complex between the two proteins. GAT1 is a stable homodimer and, interestingly, a stoichiometric shift from a 2:1 to a 2:2 GAT1:SKP1 ratio and modestly reduced affinity occurred upon glycosylation of SKP1. AlphaFold3, followed by all atom MD simulations identified a mutationally validated core hydrophobic interface which overlaps with SKP1’s FBP binding and homodimerization interfaces. The interaction was supplemented by dynamic charge-based contributions from SKP1’s intrinsically disordered CTR that also contributes to homodimerization and FBP binding albeit by distinct mechanisms. Despite the high affinity of the SKP1 homodimer, GAT1 thus appears to enzymatically modify only the SKP1 monomer. Furthermore, monomerization of SKP1 also occurs for unmodified and fully glycosylated SKP1 (Gly-Skp1) leading to the accumulation of monomeric SKP1 in solution by a substoichiometric process dependent on SKP1’s CTR. We suggest that GAT1 utilizes its SKP1 recognition mechanism to also moonlight as a SKP1 exchange factor to monomerize all glycoforms of SKP1 for complexing with FBPs, in addition to its role as a terminal GT.

## METHODS

### Strain Constructions

Cultures of human foreskin fibroblast (HFF; ATCC SCRC-1041) or hTERT HFF (BJ-5ta, ATCC CRL-4001) were maintained in Dulbecco’s modified Eagle’s medium supplemented with 10% (v/v) fetal bovine serum, 2 mM L-glutamine, and 100 units/ml penicillin/streptomycin (Corning) at 37 °C in a humidified CO_2_ (5%) incubator. Type 1 RHΔKu80ΔHXGPRT(RHΔΔ) [24] and its derivatives were cultured on these monolayers as described [25]. The previously constructed RHΔΔ/SKP1-SF (SKP1^SF^; Δku80;Δhxgprt;CAT^+^) strain [26] was edited to replace the *gat1* locus using the CRISPR strategy described in [25]. The desired replacement was confirmed by PCR as shown in Fig. S1, and replacement of the pentasaccharide glycoform of SKP1 (GGlFGGn-) with the tetrasaccharide glycoform (GlFGGn-) was confirmed by mass spectrometric analysis of immunoprecipitates as described [25] (data not shown).

### SKP1-SF Interactome Studies

A mixture of intracellular and recently lysed out parasites were subjected to non-ionic detergent solubilization and co-immunoprecipitation with anti-FLAG Ab against the SF-tag essentially as described [13]. The eluted proteins were converted to peptides using trypsin, analyzed by nLC-MS/MS in a Thermo Q-Exactive Plus workstation, and interpreted using Proteome Discoverer 2.5, all as described previously. The MS proteomics data are deposited in the ProteomeXchange Consortium via the PRIDE [27] partner repository with the dataset identifiers PXD079463 and PXD079410.

### Generation of GAT1 Isoform Plasmids

The pET15-HisTEVTgGat1 (also named pET15-His-TEV-GT8a(Gat1)) vector [18] was mutagenized using a site directed approach and primers outlined in Fig. S2. Primers were designed which incorporated the new codon for each point mutation with primer pairs utilizing a 16 bp 5’ homology region. Primer pairs were used to amplify the parental plasmid using GXL PrimeStar Polymerase (TaKaRa), and the product was digested with DpnI (New England Biolabs), ethanol precipitated and transformed into *E. coli* strain ER2566 which effectively used the 16 bp homology regions to join the 5’ ends of the PCR amplicons. Clones were screened using Sanger or Nanopore sequencing.

### Preparation of SKP1 Isoforms and GAT1

Recombinant TgSKP1 and GAT1 were expressed in *E. coli* strain ER2566 (New England Biolabs) and purified essentially to homogeneity as previously described [18]. Recombinant SKP1 lacked detectable posttranslational modifications based on intact protein mass spectrometry but was extended by a SerMet-stub preceding the normally processed Ser N-terminus after TEV-protease mediated removal of the His_6_-tag. Various glycoforms were generated by co-expressing DdPhyA and Dd/DpGnt1 in *E. coli*, and *ex vivo* using DdPgtA, TgGLT1, and TgGAT1, as described [18]. Recombinant FLAG-DdPgtA and His_6_-TgGLT1 were purified from *E. coli* as described previously [28,29]. Recombinant His_6_-TgGAT1 was purified from *E. coli* using an autoinduction protocol as previously described [18], and removal of the His_6_-tag using TEV protease left a Gly-His-Met-Ala-Ser-Met stub before the natural N-terminal Ser after processing.

### Sedimentation Velocity Experiments

Proteins were subjected to gel filtration on a S200 or S75 (16/600) column into AUC Buffer (50 mM K^+^/H^+^ phosphate (pH 7.4), 25 mM KCl). Peak fractions were diluted with the same buffer to desired concentrations as assessed by *A*_280_. For concentration series, two-fold sequential dilutions were mixed 1:1 with the protein being held constant. Samples were loaded into 12 mm double-sector Epon centerpieces with quartz windows and equilibrated for 2 h at 20°C, followed by centrifugation in a Beckman Optima XLA Analytical Ultracentrifuge at 50,000 rpm at 20 °C for 14 h. Absorbance measurements were taken at 215 nm, 230 nm 280 nm, or 295 nm depending on the concentration with a 3 mm step size. Buffer and protein parameters were computed using SedNTerp and c(s) distributions (differential sedimentation coefficient distributions) were modeled using SedFit [30,31]. Data were modeled as a continuous c(s) distribution and fitted using baseline, meniscus, frictional coefficient, buffer, protein, and noise parameters.

Distributions were overlaid in Gussi and regions of interest for each plot were integrated to determine the average S_w_ (weight-average sedimentation coefficient) position for each distribution [32]. Regions corresponding to the SKP1 monomer and dimer species were separately integrated from regions corresponding to the GAT1 homodimer and GAT1/SKP1 complexes (Fig. S6). Plots for identifying the stoichiometry of the GAT1/SKP1 complex were integrated from 3.5-6.0 corresponding to the region where the GAT1 dimer, 2:1 GAT1:SKP1, and 2:2 GAT1:SKP1 species would sediment. Average S_w_ positions were plotted in GraphPad Prism and fitted to a sigmoidal curve to determine the upper asymptote.

Normalized S_w_ values were calculated through the following means. Isotherms aimed at determining apparent affinities were integrated from 1.0-3.5 corresponding to the region where the SKP1 monomer and SKP1 dimer would sediment (Fig. S6). These isotherms were additionally integrated from 3.5 to 6.0 corresponding to the region where the GAT1 complexes would sediment. Extinction coefficients at 280 nm for each protein subunit were predicted using ProtParam tool from ExPASy [33]. Protein quantified using UV-Vis was then serially diluted and approximate extinction coefficients were calculated for relevant wavelengths according to the Beer-Lambert law. Concentrations giving an absorbance close to 0.6 were preferred for the best accuracy. The approximate contribution of each species in the GAT1/SKP1 complex peak were determined by solving for the equation 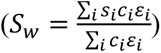 where “s” is the sedimentation value, “c_i_” is the concentration, and “ε_i_” is the extinction coefficient for each species (GAT1 dimer; 2:1 GAT1:SKP1 complex; 2:2 GAT1:SKP1 complex). Known total concentrations for each subunit were used as constraints. Free SKP1 was then calculated by subtracting the amount of complex SKP1 from the total SKP1 concentration. An OD normalized S value was then determined using the equation 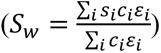 where “ε_i_” values represent the extinction coefficients of each species (SKP1 (1*Monomer + 2*Dimer); GAT1 dimer; 2:1 GAT1:SKP1 complex; 2:2 GAT1:SKP1 complex). Normalized S_w_ positions were plotted in SedPhat where a {(A+A)+(B+B) <-> A+AB+B <-> (AA)B+B <-> A+A(BB) <-> (AA)(BB); self-association A and B; macroscopic K} model was used [34]. Previously determined or estimated SKP1 homodimer affinities, an artificially low GAT1 homodimer affinity (10 nM), and previously determined sedimentation velocities were used for initial fitting. These values were held constant during initial fitting before being incrementally adjusted to yield the best fit to the data. Predicted S_w_ values for each species were calculated using HYDROPRO [35] using models obtained from AlphaFold3 [36] or MD cluster analysis as previously described [18].

### Molecular Modeling of GAT1/SKP1 Complexes

Initial models were generated using AlphaFold3 at the DeepMind web server [36]. Five structures were analyzed for each SKP1 isoform, which were each further refined using all atom molecular dynamics simulations. MD simulations were performed using the pmemd.cuda version of AMBER22.3 [37]. Amino acid residues were parameterized with the ff14SB force field [38] while glycan moieties were parameterized with the GLYCAM_06j-1 force field [39]. The system was built in a truncated octahedral box with a 15 Å distance from the solute to the end of the unit cell, neutralized with Na^+^, and solvated using TIP3P water [40,41]. Electrostatic interactions were simulated using the particle mesh-Ewald algorithm with a nonbonded cutoff of 8 Å [42]. SHAKE was used to constrain hydrogen-containing bonds which allowed for a step size of 2 fs. Solvent minimization was conducted over 5,000 cycles with steepest descent followed by 5,000 cycles of conjugate gradient with solute being restrained with a 100 kcal/mol Å^2^ energy barrier, followed by total system minimization conducted in the same manner. The system was heated to 300° K under NVT (constant particle number, volume, and temperature) conditions over 20 ns before being held steady at 300° K for an additional 10 ns. Conditions were then changed to NPT (constant particle number, pressure, and temperature) and the system was equilibrated for 270 ns. Structures were then subjected to a 1 µs production run for analysis.

AlphaFold3 generated complexes utilizing SKP1’s C-Terminal Region (CTR) were split into two groups with five structures each. The first group (Folded CTR) utilized the AlphaFold3 generated structures which contained α-helixes 7 and 8 typically observed in complexes with FBPs. Due to the high probability of disorder in this region [17], the second group (Disordered CTR) replaced the structure of SKP1’s CTR with an extended conformation using the Modeler homology modeling software [43].

Models of Gly-SKP1 were generated using AlphaFold3 structures which utilized SKP1’s CTR. First, the lowest energy state of the previously described [25] pentasaccharide from *Toxoplasma* was modeled using GLYCAM-Web [44]. Since NMR data indicated that glycosylation promotes a disorder to order transition in *Dictyostelium* SKP1’s CTR [17], AlphaFold3 was utilized to generate structures of *Toxoplasma* SKP1 with TgFBXO1, TgFBXO13, and TgFBXO14. The structure with the highest average predicted Local Distance Difference Test (pLDDT) score within SKP1’s CTR (TgFBXO13) was selected as the template for an ordered CTR. The glycan was linked to this ordered CTR according to previous modelling [16,25] utilizing UCSF Chimera [45]. SKP1’s CTR was linked to its BTB domain from the FBP bound model of SKP1 using manual PDB editing and Chimera. The positioning of SKP1’s CTR was further refined in Chimera to avoid clashes. A no-glycan control was then generated by deleting the SKP1 glycan in these models and replacing the OLP (O-linked Hyp) with a PRO designation.

Every 0.5 ns frame of the production phase was utilized for MMGBSA (Molecular Mechanics/Generalized Born Surface Area) which utilized the mmpbsa.py script [46]. The SKP1 protein (with the glycan if applicable) was treated as the ligand with the rest of the complex being treated as the receptor. An igb of 2 was used with an idecomp of 3 for pairwise interactions and 1 for per residue interaction energies. Total per residue interaction energies were averaged across all five simulations for each group. Pairwise interaction energies of electrostatic residues contributing more than -1 kCal/mol were identified and their distances were computed over the length of the production runs. Atoms representing each charge group were CZ for Arg, NZ for Lys, CD for Glu, CG for Asp, N for the N-terminal residue, and C for the C-terminal residue. Ensemble clustering has been shown to improve the reliability of protein structure analysis of dynamic proteins and complexes in several examples [18,47]. Cluster analysis was carried out in a similar manner as previously described [18]. The MD Ensemble Cluster analysis tool was utilized in Chimera. This tool groups similar structures into ensemble clusters and selects a representative structure from each cluster as previously described [48]. The top 5 representative clusters were obtained and their s-values were predicted using HYDROPRO [35]. Buried surface area was calculated using FreeSASA [49], which utilizes the Lee & Richards [50] and Shrake & Rupley [51] algorithms to calculate solvent accessible surface area (SASA). First, SASA was calculated for the SKP1 subunit, GAT1 dimer, and the total complex obtained from each representative structure from ensemble clustering. Buried surface area of each representative structure was then calculated according to the following equation: (((GAT1 dimer SASA)+(SKP1 SASA))-(Complex SASA)=(Complex buried SASA)). This was repeated for the total buried surface area, buried hydrophobic surface area (hydrophobic residues only), and buried polar surface area (electrostatic and polar residues only). Final values from each group of simulations were obtained from the weighted average of each representative structure weighted by the number of frames in each cluster.

A variability analysis was carried out using a matrix of Cα-Cα distances. Distances for each Cα pair within the structure were obtained using cpptraj. The average distances and standard deviation for each Cα pair were computed and organized into a matrix. Standard deviations organized into a Distance Variability Matrix (DVM) represent the variability of distances between residues in a protein chain and assesses the flexibility of the predicted structure. Values for each position within these DVMs were averaged across all five simulations to give the final DVM representing the specific group of simulations. Due to potential alignment artifacts for large proteins or proteins with large intrinsically disordered regions [52], RMSF was not used to score flexibility for each residue. Instead, residues were scored utilizing the DVM for each individual simulation. The Trimmed Average/Mean of the Standard Deviation utilizing 50% of the data (TASD50) score was calculated by removing the highest and lowest 25% of values to remove biases from very close and particularly flexible residues. The mean of the remaining data was obtained to yield the TASD50 score for a particular residue. TASD50 scores from each simulation were then averaged to give the final TASD50 score for each residue within a simulation group.

### Computational Scanning Mutagenesis

Computational scanning mutagenesis (CSM) was performed as previously described for the *Dictyostelium* SKP1 homodimer [53]. The top five ensemble clusters from each group of simulations representing the 2:1 GAT1:SKP1 complex with a pre-folded and disordered CTR were used as starting structures. Non-electrostatic residues on GAT1 which contributed more than -1 kcal/mol of energy in both groups of simulations were selected for mutagenesis. Starting structures were relaxed using the “fast relax” protocol in the Rosetta software suite [54,55]. Utilizing relaxed structures from each ensemble cluster, the designated residues were then subjected to saturation mutagenesis via the flex ddG protocol [56,57]. This method takes advantage of the Rosetta backrub protocol to allow for more effective optimization of mutated structures [58]. The original script was adapted to mutate both subunits of the GAT1 homodimer simultaneously. The “fa_talaris2014” scoring function was utilized with the “talaris2014” weights, and default values were used for other settings. The weighted average of the three most favorable clusters for each group (Folded or Disordered) was calculated weighted by the population each cluster.

The influence of each mutation on protein stability was calculated using the ddG monomer tool in Rosetta [59]. Five simulations representing the GAT1 dimer were clustered and representative structures from the top 5 clusters were relaxed using the “fast relax” protocol as for CSM. Relaxed structures were iteratively mutated on each subunit simultaneously at each targeted residue. Soft-repulsive weights were utilized for initial sidechain optimization. A full-atom maximum interaction distance of 9.0 Å was utilized. The local optimization restriction was set to “false” to allow wider optimization of the mutated protein structure. The three lowest scores from 50 replicates were averaged to give the initial stability score for each residue. These scores were then normalized against the wild-type control mutation for each residue to give the final residue score for each cluster. The weighted average of the three lowest scoring clusters for each mutation was then divided by two, to account for each subunit, to give the final stability score for each mutation.

### SKP1 Hydroxylation Assays

SKP1 hydroxylation assays were utilized to determine if GAT1 can inhibit SKP1 hydroxylation. Common Master Mix solutions were prepared for each reaction condition. Master Mix 1 (4x concentration of SKP1) contained 50 mM HEPES-NaOH (pH 7.4), 400 nM SKP1, 10 µg/ml leupeptin, 10 µg/ml aprotinin, 0.25 mg/ml BSA, 5 µM FeSO_4_, 1.5 mM α-ketoglutarate, 2 mM ascorbate, 0.8 mg/ml catalase, and 1 mM TCEP. Master Mix 2 (4X concentration of PHYa) contained 50 mM HEPES-NaOH (pH 7.4), 10 µg/ml leupeptin, 10 µg/ml aprotinin, 0.25 mg/ml BSA, 5 µM FeSO_4_, 1.5 mM α-ketoglutarate, 2 mM ascorbate, 1 mM TCEP, and 20 nM PHYa. GAT1 and control inhibitors were mixed with Master Mix 1 to yield a pre-initiated mix at 75% the total volume, and the reaction initiated by rapidly mixing with Master Mix 2. Final inhibitor concentrations were 100 nM GAT1 (1:1), 200 nM GAT1 (2:1), 600 nM GAT1 (6:1), 2 µM GAT1 (20:1), 500 µM protocatechuic acid, or 500 µM folate. Reactions were incubated for 23 min and 1.5 h and quenched with SDS-PAGE loading buffer. Background was measured at time zero by adding SDS-PAGE Sample buffer to the pre-initiation mix. Samples were analyzed by SDS-PAGE and Western blotting, and the presence of hydroxylated SKP1 (HO-SKP1) was measured by probing with pAb UOK85, which is specific for this isoform [12], followed by alexa-680 labeled goat anti-rabbit IgG. A calibration curve was established by mixing fully hydroxylated or unmodified SKP1 in varying ratios and quantitated densitometrically in a Li-Cor Odyssey infrared scanner with processing using NIH Image J software.

### Fbs1-GAT1 Competition Assays

Gel filtration chromatography was performed essentially as previously described [15]. Briefly, the column buffer consisted of 50 mM HEPES-NaOH (pH 7.4), 60 mM NaCl, and 1 mM DTT. Solutions containing 5 µM Fbs1, 10 µM GAT1, or appropriate mixtures were prepared followed by addition of 5 µM SKP1. Solutions were preincubated for 30 min before injecting 50 µl onto a Superdex 200 PC 3.2/30 column on a Pharmacia SMART-System at 80 µl/min with *A*_280_ monitoring. 50 µl fractions were analyzed by SDS-PAGE and Coomassie blue staining, or by Western Blotting using pAb UOK75 for SKP1 and murine anti-His Ab for Fbs1, followed by alexa-680 goat anti-rabbit or goat anti-mouse IgG. Densitometry was employed to analyze performed as above.

### UDP-Glo Enzyme Assays

Assays were based on detection of the UDP reaction product from the donor substrate UDP-Gal, as modified from [60].

#### Activity Against Small Molecule Substrates

Recombinant TgGAT1 variants were incubated at 100 nM in 50 mM HEPES-NaOH (pH 7.0), 500 µM potassium phosphate (pH 7.4), 1.5 mM KCl, 100 mM NaCl, 2 mM DTT, 2 mM MnCl_2_, 0.2% (v/v) Tween-20, 5% (w/v) Ficoll-400, and 1 mg/ml BSA. For pNP-maltoside curves, UDP-Gal was held constant at 100 µM with variable pNP-maltoside concentrations. For UDP-Gal curves, pNP-maltoside was held constant at 50 mM with variable UDP-Gal concentrations. Reactions were initiated by adding GAT1 in a total volume of 10 µl and run at 37°C for 10 min (pNP-maltoside trials) or 5 min (UDP-Gal trials). Generated UDP was quantified using the UDP-Glo Assay (Promega). 5 µl of each reaction was mixed with 5 µl of final Nucleotide Detection Reagent in every other well of a white 384 well plate. Plates were incubated for one hour before being read using a Biotek Synergy 2 luminometer. A UDP standard curve was used to quantify generated UDP, while no GAT1 and hydrolysis controls confirmed minimal background signal. Data points were averaged over four replicates and fit to the Michaels Minton equation.

#### Activity Against SKP1

Recombinant TgGAT1 variants were incubated at 100 nM in 50 mM HEPES-NaOH (pH 7.0), 19 mM potassium phosphate (pH 7.4), 22 mM KCl, 100 mM NaCl, 2 mM DTT, 2 mM MnCl_2_, 0.2% (v/v) Tween-20, 5% (w/v) Ficoll-400, 1 mg/ml BSA, and 100 µM UDP-Gal. 7 µM GlFGGn-SKP1 was the substrate and 50 mM pNP-maltoside and water served as positive and negative controls. The UDP product was quantified as above, averaged over three replicates.

## RESULTS

### Loss of GAT1 Uniquely Alters the SKP1-SF Interactome

A previous study probed the interactome of SKP1 by anti-FLAG based immunoprecipitation of SKP1 from SKP1-SF strains [13]. Here the approach was extended to the analysis of *gat1*-KO cells after re-deriving the KO in a previously described SKP1-SF strain (Fig. S1). *gnt1*-KO and normal (RHΔΔ) cells were analyzed for comparison. As before, multiple FBPs were detected in their SKP1-SF interactomes at levels that varied over 2 orders of magnitude based on peptide spectrum matches. They could be classified as enriched in normal vs. *gnt1*-KO backgrounds (designated Group 2, Fig. 2B) or unaffected by the genetic background (Group 1, Fig. 2C). The representation of Group 2 FBPs in *gat1*-KO cells, which accumulate the tetrasaccharide glycoform of SKP1, was intermediate between normal and *gnt1*-KO cells, similar to what was previously observed in *pgtA*-KO cells (which accumulate the monosaccharide glycoform). As expected, Group 1 FBPs were similarly represented in all strains including *pgtA*-KOs [13], and SKP1-SF was detected at similar levels in each (Fig. 2D). In contrast, 4 FBPs exhibited a novel pattern in that they were more highly represented in the *gat1*-KO compared to both normal and *gnt1*-KO backgrounds (Fig. 2A). Now classified as Group 3 rather than Group 1 FBPs, they might differentially interact with the tetrasaccharide glycoform of SKP1-SF. However, the presence of GAT1 itself in the interactome of all genetic backgrounds analyzed [13] suggested a stable interaction with SKP1, which might indirectly influence interactions with select FBPs, whose F-box sequences are highly varied. Therefore, the interactions of recombinant versions of GAT1 and SKP1 were characterized *in vitro*.

**Figure 2.**
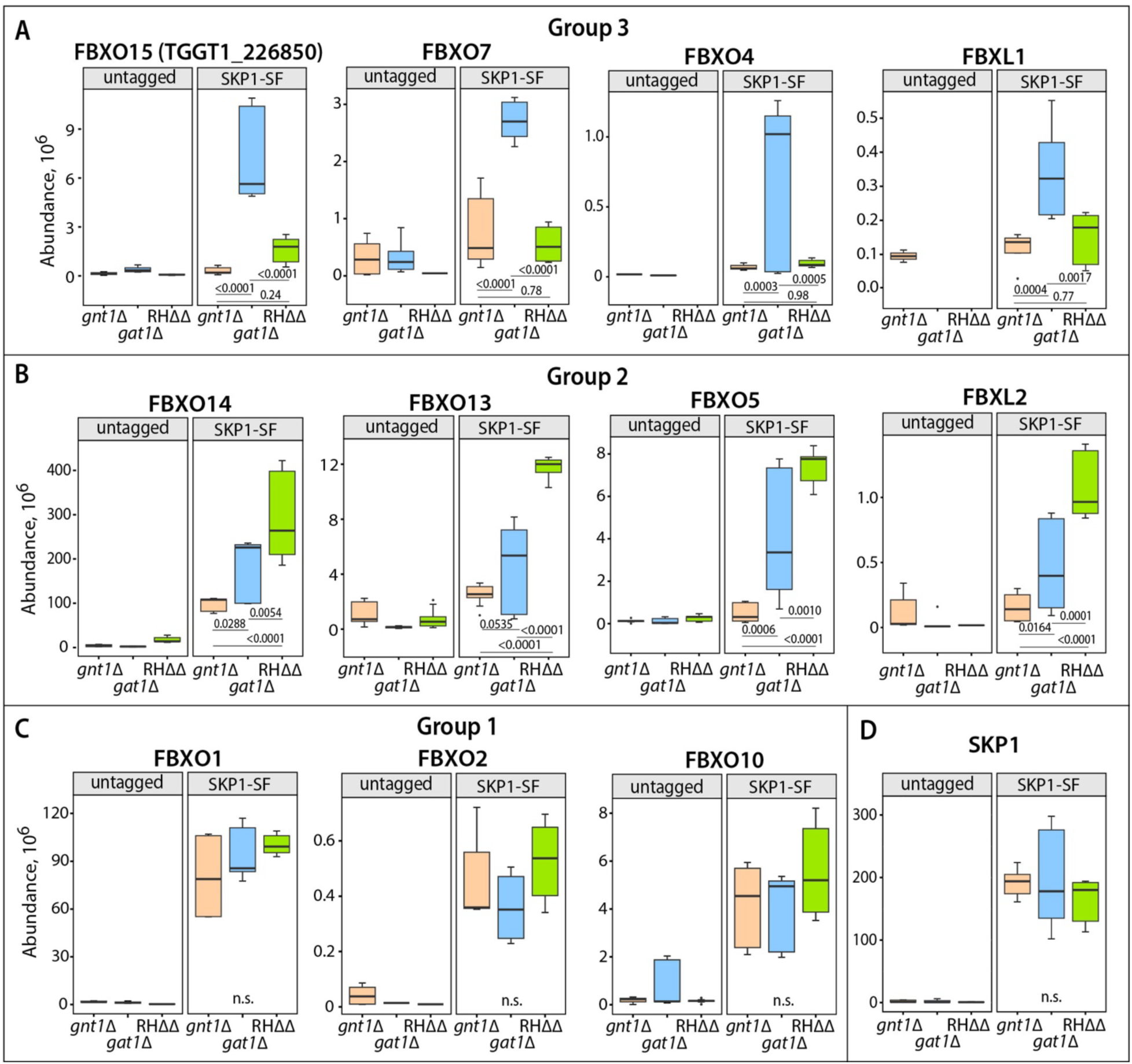
SKP1-SF interactome inf *gat1*-KO parasites. Comparison of FBPs detected in SKP1-SF interactomes from *gat1*-KO (*gat1*Δ), *gnt1*-KO (*gnt1*Δ) and normal (RHΔΔ) strains. Quantification was based on counts from Peptide-Spectrum Matches (PSMs) from nLC-MS-MS analysis of co-IPs. Non-specific levels of interaction are given by parallel analysis of untagged strains in the left panels. Boxplots represent expression values for 3 independent biological replicates, each with 3 technical replicates. Boxes represent the two middle quartiles, the line within the box represents the median, and the whiskers represent the lowest the highest values of the dataset. (A) Group 3 FBPs, most highly represented in the SKP1-SF interactomes of *gat1*-KO parasites. (B) Group 2 FBPs, most highly represented in RHΔΔ interactomes. (C) Group 1 FBPs showing no significant differences among strains. (D) Similar levels of SKP1-SF were captured in each strain. p-values report significance tested using one-way ANOVA with Tukey HSD post-test. n.s., difference not significant (p>0.05).

### GAT1 Stably Binds SKP1 in a Complex whose *K*_d_ and Stoichiometry are Dependent on SKP1’s Glycosylation State

To characterize the nature of GAT1’s association with SKP1 indicated by the co-IP pulldowns, SKP1 and GAT1 were expressed in *E. coli* and purified to homogeneity. The recombinant proteins were identical to the native proteins, except for an N-terminal hexapeptide extension relative to the normally processed form that has Met removed and potential N-terminal acetylation, owing to TEV-protease mediated removal of a His_6_-tag used for purification. Using sedimentation velocity, clear peaks were observed for the individual homodimers of both SKP1 and GAT1 at 5 µM concentration (Fig. 1E). The lack of visible monomer species for either protein is consistent with the previously reported high affinity homodimer of each protein [18,25]. Upon mixing equimolar concentrations of SKP1 and GAT1, a reduction in the magnitude of the SKP1 homodimer peak was observed. This accompanied a shift in the GAT1 dimer peak to a position consistent with a 2:1 GAT1:SKP1 complex based on modeling (Fig. 1E). This indicates dissociation of the SKP1 homodimer in favor of interacting with GAT1. With an approximate 2-fold excess of GAT1, mostly complete depopulation of the SKP1 homodimer species was observed with a minimal shift in the position of the GAT1:SKP1 complex (Fig. 1E). The lack of discrete species for either the GAT1 homodimer or a 2:2 GAT1:SKP1 complex suggests a relatively rapid equilibrium to the formation of a 2:1 complex on the timescale of the experiment (Fig. 1E).

Protein concentrations were then lowered to more physiological levels. SKP1 was held constant at 500 nM, which still allowed detection of the relatively weak SKP1 absorbance signal, while the concentration of GAT1 was varied. The region of each distribution corresponding to the GAT1 homodimer and the GAT1/SKP1 complex was then integrated to determine the saturating S_w_ value (Fig. S6). As shown in Fig. 3A, the S_w_ position for the GAT1/SKP1 complex effectively saturated at a position slightly below the 4.7 value predicted for the 2:1 GAT1:SKP1 complex. The significance of SKP1’s pentasaccharide was evaluated with the same titration using Gly-SKP1. Interestingly, the saturating S_w_ position for the GAT1:SKP1 species increased to a value consistent with a 2:2 GAT1:SKP1 complex (Fig. 3C). This indicates that stoichiometry of the GAT1/SKP1 complex is dictated by SKP1’s glycosylation status. Glycosylation of SKP1 was previously reported to weaken SKP1 homodimerization by disrupting a charge-based fuzzy interaction of SKP1’s CTR [18]. Given this, we next examined a potential contribution of SKP1’s CTR to the stoichiometry of the GAT1/SKP1 complex. As shown in Fig. 3E, SKP1ΔCTR formed a 2:2 GAT1:SKP1 complex analogous to the behavior of Gly-SKP1. Thus, the data indicate that SKP1’s CTR drives asymmetry of the complex. This suggests glycosylation controls stoichiometry through effects on the SKP1 protein itself rather than a lectin-like binding interaction between GAT1 and the glycan.

**Figure 3.**
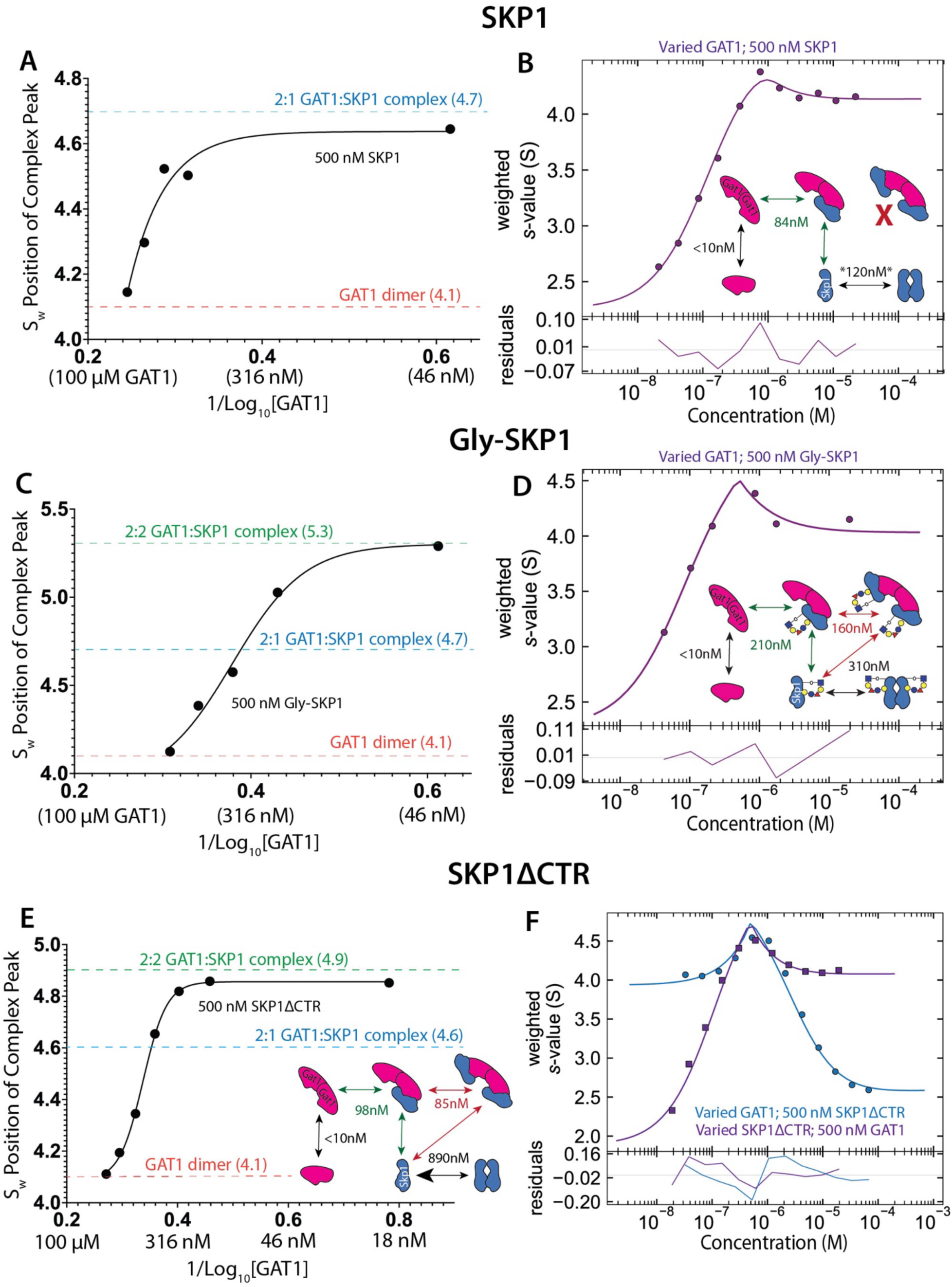
Sedimentation Velocity Analysis of the GAT1:SKP1 Complex. Plots showing the average S_w_ position for the GAT1-homodimer and GAT1:SKP1 complex peaks when 500 nM SKP1 (A), Gly-SKP1 (C) or SKP1ΔCTR (E) are titrated with GAT1. The decreasing concentrations of GAT1 left to right represent increasing ratios of SKP1 to GAT1. S_w_ positions were fit to a sigmoidal curve and the upper asymptote was analyzed. The positions of the GAT1 homodimer (orange), 2:1 GAT1:SKP1 complex (blue), and 2:2 GAT1:SKP1 complex (green) are shown by dashed lines. The weighted sums of S_w_ positions for the SKP1 monomer, SKP1 homodimer, GAT1 monomer, GAT1 homodimer, and GAT1:SKP1 complexes were plotted in SedPhat and the apparent affinity of each interaction was modeled with SKP1 (B), Gly-SKP1 (D), and SKP1ΔCTR (F). Isotherms where the SKP1 (purple) or GAT1 (blue) concentration was varied are shown. Apparent affinities for each relevant interaction were modeled and are depicted in interaction diagrams. Supporting data are in Figs. S3-S6.

We next modeled the apparent affinities of each interaction using SedPhat. The range of S_w_ values associated with the SKP1 monomer and SKP1 dimer were integrated and used alongside the range associated with the GAT1 dimer and GAT1/SKP1 complex (Fig. S6). These integrations were used to generate a normalized S_w_ value which was plotted as a function of GAT1 concentration into an isotherm. As shown in Fig. 3B, these plots exhibit three regions of interest. The component observable at negligible GAT1 concentrations represents the SKP1 monomer-dimer equilibrium. The isotherm at this region represents the contribution of the SKP1 homodimer at this concentration (500 nM). As more GAT1 is titrated into the solution, the average S_w_ of the system increases consistent with both the increased representation of the GAT1 species and greater incorporation of SKP1 into the larger GAT1:SKP1 complex. The average S_w_ of the system peaks when the maximum amount of both SKP1 and GAT1 have been bound up by the GAT1/SKP1 complex. The position, shape, and height of this peak are determined by both the stoichiometry and affinities of this complex. Once this peak has been attained, titration of increasing GAT1 drives the average S_w_ toward a value representing the GAT1 homodimer. Overrepresentation of the GAT1 homodimer in this region brings down the average S_w_ value of the system.

The SKP1 homodimer [18] and GAT1 homodimer [25] were previously characterized to have affinities with *K*_d_ values ≤50 nM. Based on these values, the S_w_ values for each species and complex, and the stoichiometries of the GAT1/SKP1 complexes, apparent affinities for each GAT1/SKP1 interaction were calculated. The apparent affinity of SKP1 with GAT1 was estimated at 84 nM (*K*_d_) (Fig. 3B) and attempts to model the interaction as a 2:2 GAT1:SKP1 complex were unsuccessful. The apparent affinity for the GAT1 homodimer modeled to <10 nM consistent with previous observations. Interestingly, the homodimer affinity of SKP1 modeled to ∼120 nM, which is more than twofold higher than previously estimated [18]. This difference may result from a dissociative effect of the GAT1 dimer on the SKP1 homodimer (see Fig. 8A below). Upon modeling a similar isotherm with Gly-SKP1, two binding events were effectively modeled with each having an apparent affinity close to 200 nM, or about 2.5-fold higher than that of unmodified SKP1 (Fig. 3D). This is also weaker than the affinity seen with SKP1ΔCTR suggesting that rather than contribute to binding, Gly-SKP1’s CTR slightly disfavors GAT1 binding (Fig. 3F). This may represent the loss of favorable contacts upon glycosylation, a shift toward a less favorable SKP1 conformational ensemble, entropic excluded volume effects, or a combination of effects.

### SKP1’s CTR is Likely Relatively Disordered in the GAT1/SKP1 Complex

To explore the mechanism by which SKP1’s CTR may control complex asymmetry, we turned to computational modeling starting with AlphaFold3. We first modeled the 2:1 GAT1:SKP1 complex observed for unmodified SKP1 (Fig. 4A). In this model GAT1 forms a homodimer which matches closely (Cα RMSD = 0.947 Å) with the crystal structure solved for the highly similar *Pythium* GAT1 (Fig. S7). A relatively large and poorly conserved intervening loop is present in GAT1 relative to *Pythium* GAT1 [25] as a region of low confidence in AlphaFold3’s pLDDT which interprets confidence in local structural features (Fig. 4C). It is also clearly visible using AlphaFold3’s Predicted Alignment Error (PAE) matrix (Fig. 4B), which interprets confidence in the global structural model. This can indicate flexible or disordered regions as well as reliability of the predicted interface. A low confidence helix is predicted within this intervening loop (Fig. 4A,C) which may be the result of patches of electrostatic residues. Large and especially alternating patches of electrostatic residues are known to promote free helix formation, such as E-R/K helixes [61–63]. SKP1 is positioned with high confidence on a single subunit of the GAT1 homodimer (Fig. 4A-C). No interactions of an intact SKP1 homodimer were observed when a second SKP1 subunit was modeled. The positioning of SKP1 onto the single GAT1 subunit is likely directed by a core hydrophobic interface surrounded by both polar and electrostatic interactions (Figs. 5,6). This interface is also utilized by the SKP1 homodimer and offers a mechanism for incorporating a single SKP1 subunit for a first binding event. This is a necessary feature of a modeled GAT1/SKP1 complex due to a 2:1 GAT1:SKP1 complex being observed at concentrations even orders of magnitude above the SKP1 homodimer *K*_d_ [18]. Additionally, this prediction is consistent with gel filtration studies that show that elution of SKP1 with the soluble mammalian FBP Fbs1 is not affected by addition of GAT1 (Fig. S9). SKP1 bound into a complex with GAT1 becomes recalcitrant to hydroxylation by PHYa (Fig. S10), an effect also observed with FBPs [63] and *Dictyostelium* AgtA [64]. SKP1’s N-terminal extension is modeled with low confidence extending into the solvent consistent with a disordered protein region (Fig. 4A,C). SKP1’s internal loop is modeled with relatively low confidence but higher confidence than other typically disordered regions of the protein (Fig. 4A-C). This likely stems from the presence of flanking anchoring sites for this disordered protein segment as opposed to the single anchoring site for the N- and C-termini. SKP1’s CTR is modeled by AlphaFold3 with low confidence though folding as helixes 7 and 8 is similar to complexes with FBPs [10,11], likely owing to an abundance of FBP containing structures of SKP1 in the PDB. Given the low confidence in this region, there is a high likelihood that this portion of the protein lacks stable structure. Positioning of these helixes is also relatively inconsistent between independently generated models (Fig. S8). Whether or not SKP1’s CTR is assumed to be disordered, the positioning of SKP1 orients the CTR to face toward GAT1’s active site (Figs. 4A, S10). This is significant, as orientation of SKP1’s CTR toward GAT1’s active site would likely be necessary for efficient enzymatic activity of GAT1 for SKP1.

**Figure 4.**
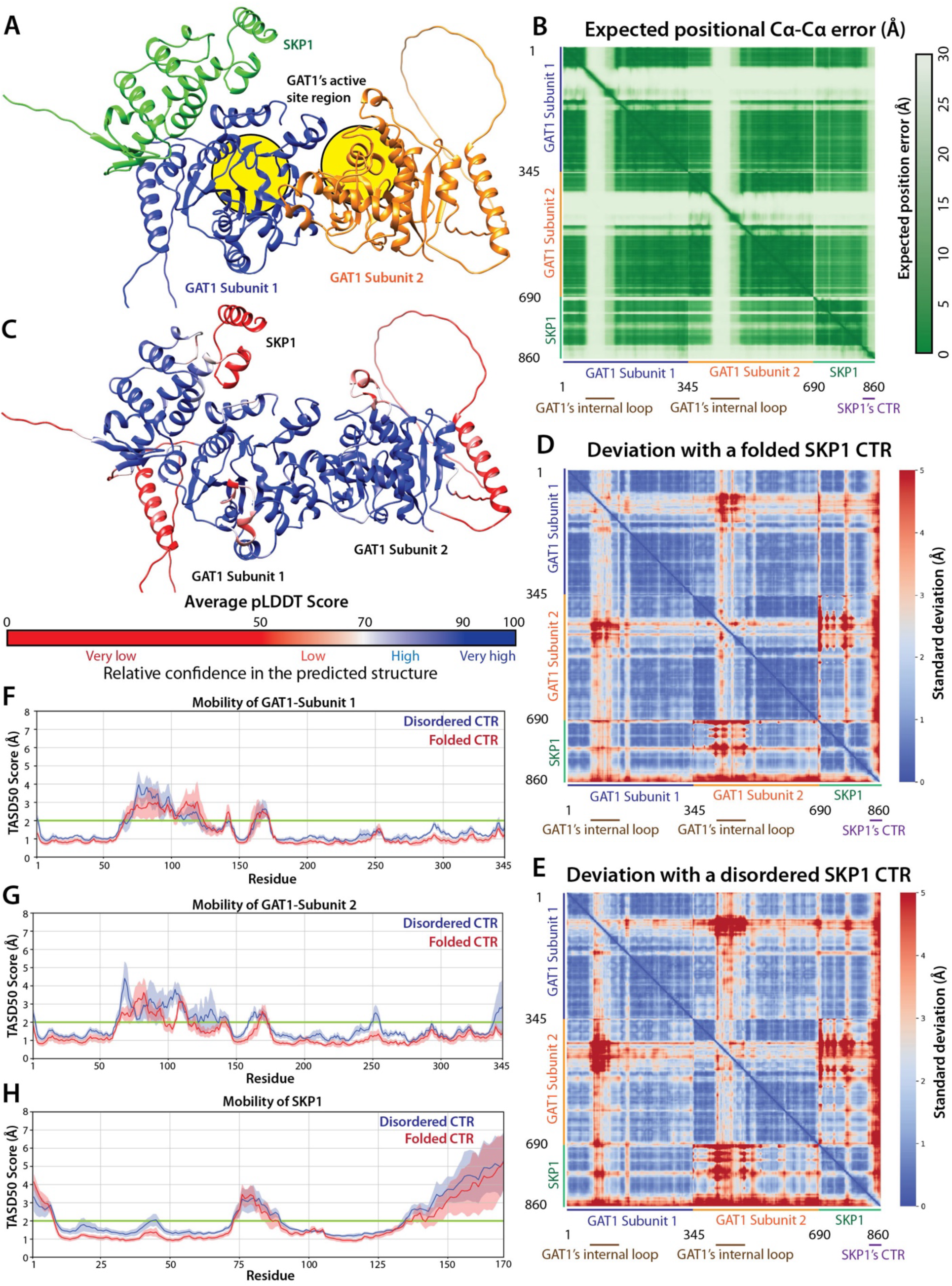
Computational Modeling of the GAT1:SKP1 Complex. (A) General structure of the GAT1/SKP1 complex as predicted by AlphaFold3. SKP1 (green), GAT1 Subunit 1 (blue), and GAT1 Subunit 2 (orange) are shown with the location of each active site circled in yellow. (B) Predicted Alignment Error (PAE) matrix for the AlphaFold3 generated GAT1/SKP1 complex. The position of each complex subunit and intrinsically disordered regions (high error scores) are shown. (C) Average predicted Local Distance Difference Test (pLDDT) scores for the GAT1/SKP1 complex are shown and colored by confidence. (D) Distance variability matrix for the group of simulations where a folded SKP1 CTR was used to start MD simulations. Cα distances are colored by the average of their standard deviations across all five simulations. (E) Same as (E) except that a disordered SKP1 CTR initiated the MD simulations. (F-H) Calculated TASD50 scores (Trimmed Average/Mean of the Standard Deviation utilizing 50% of the data) for each residue in GAT1 Subunit 1 (F), GAT1 Subunit 2 (G), and SKP1 (H). The 2 Å variability cutoff for “disordered” segments is highlighted green. Values for simulations initiated with a folded (red) or a disordered (blue) CTR are shown with shading representing the standard error of the mean.

**Figure 5.**
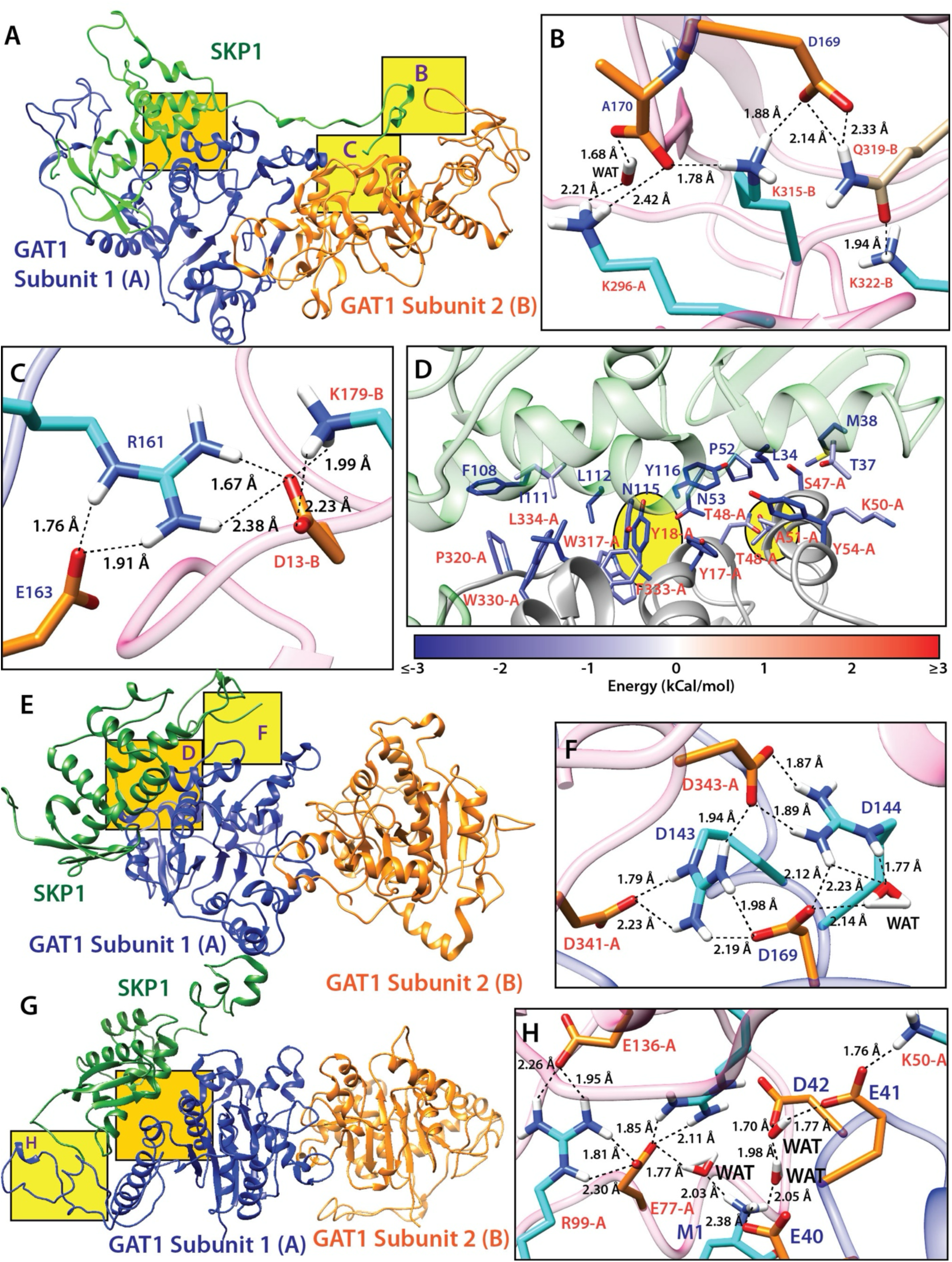
Examples of Interactions in the GAT1/SKP1 Complex. Frequently occurring structures from the molecular dynamics simulations are shown in panels A, E and G, with zoom-ins keyed to the square boxes as noted. (A) A representative structure of the GAT1/SKP1 complex in Simulation 5. Subunits are colored as in Fig 4A, with SKP1 in green and the GAT1 subunits in blue and orange. Yellow boxes highlight regions shown in panels B and C, and the orange box, which is shared with panels E and F, denotes a region illustrated in panel D but not from this simulation. (B, C) Electrostatic interaction networks involving the SKP1 CTR. Protein backbones are shown as ribbons with GAT1 colored hot pink and SKP1 colored blue, and SKP1 residue labels are blue while GAT1 residue labels are orange. Residues from GAT1 Subunit 1 are denoted by “-A” and “-B” for Subunit 2. (D) A structure of the core hydrophobic interface from Simulation 3. The protein backbones are shown as ribbons with SKP1 being green and GAT1 being grey. Only consistent interactions (<–1 kCal/mol contribution in both simulation groups) are shown. Residues are colored by interaction energies predicted by MMGBSA in the Disordered CTR group of simulations. (E, G) Frequently occurring structures of the GAT1/SKP1 complex from Simulations 2 and 1 respectively. (F, H) Zoom-in views of electrostatic interaction networks from Simulations 2 and 1, respectively, with labels and colors as in (B). Examples in Fig. S16 document that these examples occur at high frequency in most of the simulations.

**Figure 6.**
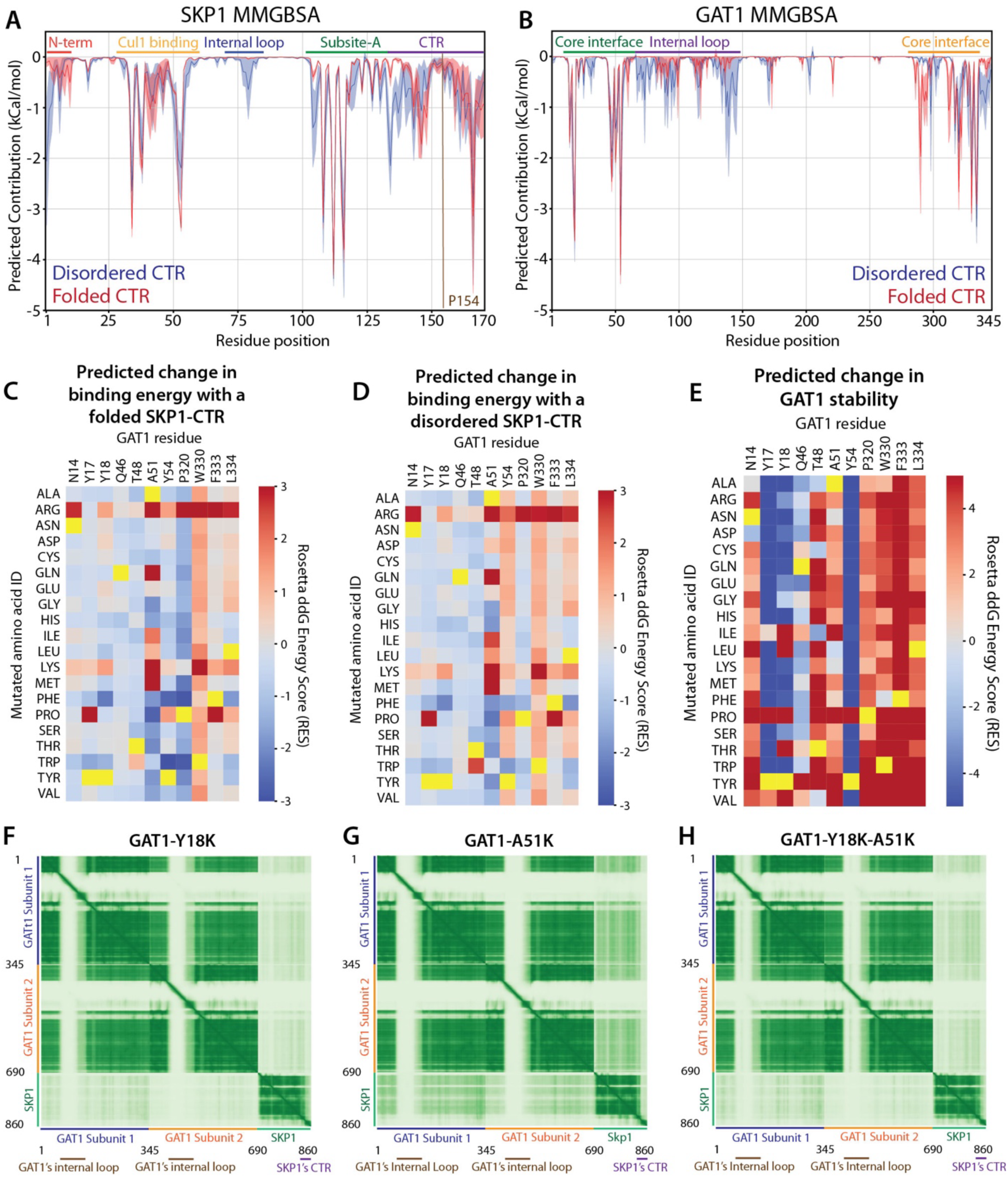
Energetic and Mutational Analysis of the GAT1/SKP1 Complex. (A) MMGBSA predicted binding energies toward GAT1 for residues in SKP1. Values for simulations starting with a folded (red) or a disordered (blue) CTR are shown with shading representing the standard error of the mean. (B) Same as in (A) except for predicted binding energies toward SKP1 for residues in GAT1. (C) Predicted change in binding energy for amino acid substitutions made for significant GAT1 residues in simulations where a folded SKP1 CTR was used for starting structures. (D) Same as (C) except using simulations for which a disordered CTR was used. (E) Predicted changes in protein stability for mutations made against residues in GAT1. (F-H) PAE matrices for a 2:1 GAT1:SKP1 complex where the GAT1(Y18K) (F), GAT1(A51K) (G), and GAT1(Y18K/A51K) (H) variants are examined. See Fig. S13 for supporting data.

AlphaFold3 can often correctly model the general structure of a protein complex but is less able to model specific interactions [66–68] or intrinsically disordered regions [67–69]. Thus, we turned to all atom molecular dynamics simulations to investigate specific interactions within the GAT1/SKP1 complex and their dynamics. Five structures generated by AlphaFold3 were directly used to initiate independent molecular dynamics simulations termed “Folded CTR Simulations.” A second group of five structures was generated where SKP1’s CTR was assumed to be initially disordered termed “Disordered CTR Simulations.” This was done by deleting the residues associated with SKP1’s CTR and modeling them back in an extended conformation using Modeler as done previously with the SKP1 homodimer [18]. After an initial heating step, structures were subjected to refinement through a 270 ns equilibration step which allowed for structural adjustment based on Amber’s force fields. Following this equilibration step, a 1 µs simulation was run and the structural and dynamic properties of the complex were analyzed. Given the relatively large size of the complexes (up to 100 kDa) and large proportion of disordered regions, a high impact from alignment artifacts can be expected from techniques such as RMSF [52]. Given this, an approach utilizing distance matrixes was selected as more appropriate for analyzing protein flexibility. To do this, we calculated and plotted the standard deviation of Cα-Cα distances into a Distance Variability Matrix (Fig. 4D-E), which showed similar dynamic behavior for both groups of simulations. The regions with high degrees of flexibility correspond well to the regions of low confidence in AlphaFold3’s PAE matrix. A high degree of flexibility is observed within the internal loop of each GAT1 subunit. Additionally, high flexibility is observed within the N-terminal region, internal loop, and CTR of SKP1. Relatively low variability is observed between the well-ordered regions of each complex subunit indicating a substantially immobile complex with stable protein-protein interfaces including with SKP1. To further analyze the mobility of individual residues, the trimmed mean of average standard deviations was calculated for all Cα-Cα distances per each atom; 50% of the total values (25% of the highest and lowest) were excluded when calculating the mean standard deviation of each residue. This reduced artifacts from both highly mobile and nearby residues to give a Trimmed Average Standard Deviation (TASD50) score for each residue. As shown in Figs. 4F-H, the protein subunits from each group of simulations behaved similarly. Regions of high flexibility (TASD50 > 2Å) correspond well to regions of the structure with relatively low pLDDT confidence scores. Specifically, a relatively high TASD50 score for SKP1’s CTR was determined for both groups of simulations supporting the model where SKP1’s CTR remains highly mobile within a complex between SKP1 and GAT1.

### SKP1 Binds GAT1 Utilizing a Core Hydrophobic Interface

The GAT1/SKP1 interface buries an expansive surface area of 3380 Å^2^ when a disordered CTR is modeled and 3180 Å^2^ when a folded CTR is modeled. The buried hydrophobic surface area represents a large component of this with 2070 Å^2^ and 1840 Å^2^ for the disordered and folded CTRs, respectively. To determine which residues are likely most important for maintaining the GAT1/SKP1 complex, we turned to analysis through MMGBSA. As graphed in Fig. 6A and illustrated in Fig. 5D, significant contributions were observed to occur from SKP1 F108, I111, L112, N115, and Y116, which also contribute significantly to SKP1 homodimerization via Subsite-A (Fig. 1B) [18]. Significant, but less consistent contributions were observed for other residues in Subsite-A including Q104, K105, A109, M133, and I134. Hydrophobic contacts from SKP1’s N-terminal lobe, involved in CUL1 binding, included L34, T37, M38, P52, and N53 and less consistent contributions from E41, L51, and V54. Significant, but highly conformationally dependent interactions were also observed within SKP1’s CTR that is reminiscent of previously observed homo-interactions within SKP1’s fuzzy Subsite-B (Fig. 1B,C,F). Importantly, many SKP1 residues identified to be most important for binding to GAT1 are also important for SKP1 homodimerization. This and our previous AUC data support two interpretations about the GAT1/SKP1 interaction: it is competitive with SKP1 homodimerization, and mutagenesis studies targeting the SKP1 side of the interface would be confounded by effects on SKP1 homodimerization.

From the GAT1 side, large (<–1 kcal/mol) energetic contributions to the interaction were observed for both simulation groups from GAT1 residues N14, Y17, Y18, S47, T48, K50, A51, Y54, W317, P320, W330, F333, and L334 (Figs. 6B, 5D). Less consistent but still significant interactions were made by F16, S45, Q46, Q55, R137, R139, H289, M290, Y298, S312, Q319, A321, K331, and T335. The charged residues are suspected to contribute transient protein contacts (Fig. 5). To test the role of the largely hydrophobic interface, we turned to Computational Scanning Mutagenesis (CSM) with the Rosetta software suite. Contact residues from the GAT1 portion of the interface were targeted because the predicted SKP1 interface is also involved with FBP binding and SKP1 homodimerization. The role of substitutions by every other aa in influencing the GAT1/SKP1 complex were tested using the flex ddG algorithm. Substitutions in both simulation groups (folded and disordered SKP1 CTR) were predicted to have similar effects on the stability of the GAT1/SKP1 interface (Fig. 6C,D). The ddG Monomer algorithm was then utilized to test the influence of each mutation on the stability of GAT1 (Fig. 6E). From these datasets, the Y18K and A51K point mutations were predicted to weaken the SKP1/GAT1 interface while simultaneously preventing destabilization of GAT1 and were chosen for further analysis. Since AlphaFold3 has proven useful in predicting the effects of mutagenesis on protein-protein complex formation [68], the PAE matrixes from each of the mutant GAT1/SKP1 structures were analyzed and showed a large decrease in the confidence of placing SKP1 in the GAT1/SKP1 complex (Fig. 6F-H). A relatively small decrease in the pLDDT score was observed for interface residues in each instance (Fig. S15), but predicted SKP1/GAT1 docking maintained a higher degree of confidence with this metric. Additionally, the Y18K and Y18K/A51K mutants resulted in inconsistent placement of SKP1 in the complex. Taken together, despite AlphaFold3 confidence metrics showing high confidence in each protein’s fold, the ability to place SKP1 properly in the GAT1/SKP1 complex was substantially hindered by these point mutations.

The Y18K and A51K point mutations were generated in recombinant GAT1 as a test of the predicted SKP1/GAT1 interface. As a check for preservation of GAT1’s galactosyltransferase activity, the recombinant enzyme was assayed in the presence of pNP-maltoside as the acceptor substrate and UDP-galactose as the donor substrate. Activity was assessed based on the generation of the product UDP, which was assayed based on a coupled reaction that generates ATP that drives a luciferase reaction. As shown in Fig. 7, the data indicate that the individual point mutants, and the Y18K/A51K double mutant, maintained similar apparent *K*_m_ and *V*_max_ values relative to the normal enzyme for UDP-Gal (panel A) and pNP-maltoside (panel B), suggesting that overall protein structure is preserved. Similar differences were observed by assaying the low level of spontaneous hydrolysis of UDP-Gal in the absence of acceptor (data not shown). A previous study [25] showed that GAT1 is substantially more active toward GlFGGn-SKP1 than pNP-maltoside and the native glycan linked to the pNP aglycon (GlFGGn-pNP). This indicated a requirement for recognizing SKP1 to obtain optimal activity. Consistent with this interpretation, minimal activity was detected for the Y18K and A51K mutants and none was detected for the double mutant using a GlFGGn-SKP1 substrate (Fig. 7C). Interestingly, the slightly higher activity of the A51K relative to the Y18K mutant is more consistent with the magnitude of the AlphaFold3 prediction (Fig. 6F,G) compared to that of the Rosetta prediction (Fig. 6C,D). The finding that the point mutations selectively affected activity toward GlFGGn-SKP1 relative to pNP-maltoside provides strong support for the predicted interface with GAT1 as depicted in Fig. 7D (orange oval).

**Figure 7.**
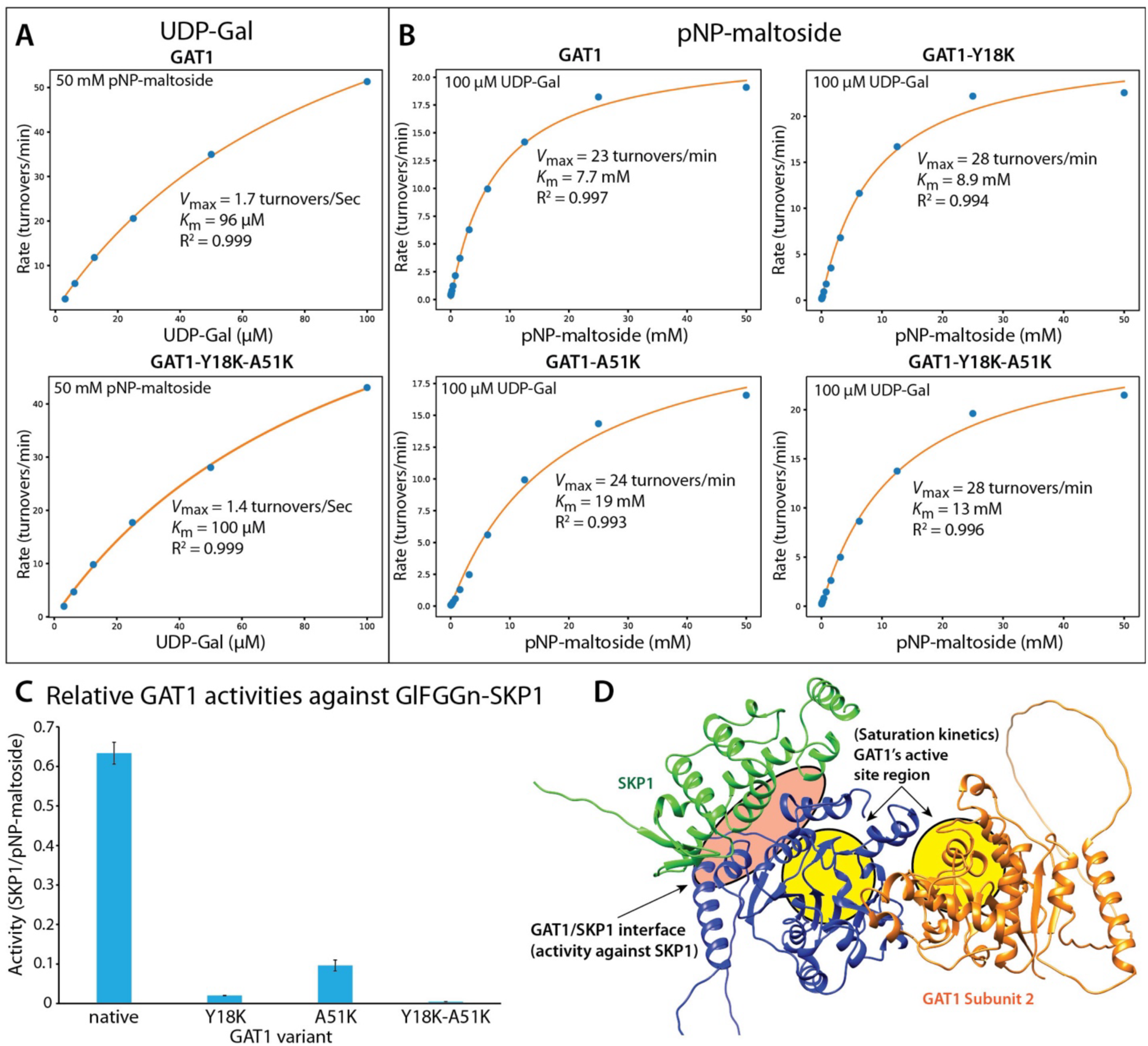
Enzyme Assays for Tested GAT1 Variants. (A) UDP-Gal “saturation curves” with 50 mM pNP-maltoside used as the receptor. (B) pNP-maltoside “saturation curves” with 100 µM UDP-Gal used as the donor sugar. (C) Ratio of catalytic turnover rates measured between 7 µM GlFGGn-SKP1 and 50 mM pNP-maltoside using 100 µM UDP-Gal for both reactions. (D) Map of where each group of enzymatic assays measure effects of point mutations of the GAT1 structure. Small molecule “saturation curves” measure the impact of point mutations on GAT1’s catalytic core (yellow circles). SKP1 reactions measure the impact on GAT1’s ability to recognize its SKP1 substrate (orange oval).

### GAT1 and SKP1 Contribute a Hybrid Ordered and Fuzzy Interface

SKP1’s CTR and GAT1’s internal loop showed high mobility (Fig. 4) and exhibited a high diversity of contacts with one another and other surfaces on each protein (Fig. 5A,E,G). In all simulations a core hydrophobic and ordered interface similar to that depicted in Fig. 5D is present. As visualized through MMGBSA, significant electrostatic based interactions are present within the GAT1/SKP1 complex (Fig. S12). Much of this energy is centered around clusters of positively or negatively charged residues, an effect observed in simulations of SKP1’s fuzzy interface [18]. To observe the kinetics of these interactions, pairwise MMGBSA energies were calculated for each pair of oppositely charged electrostatic residues. Distances between electrostatic pairs with contributions greater than -1 kCal/mol in an individual simulation were plotted across the length of the simulation (see Fig. S16 for representative examples). A total of 146 examples of these electrostatic pairs were observed across all 10 simulations to give an average of 14.6 electrostatic pairs per simulation. A strong bias was observed in favor of a starting disordered CTR with 97 electrostatic pairs versus the 49 observed with a folded CTR. This indicates that conformations accessible within “disordered CTR” simulations provided 19.4 productive contacts per simulation versus the 9.8 observed when SKP1’s CTR was pre-folded. The transient formation of electrostatic interaction networks was observed in every simulation. For example, SKP1’s CTR frequently formed interactions with charge clusters both on Subunit-1 (Fig. 5F) and Subunit-2 (Fig. 5B,C) of GAT1. Interestingly, GAT1’s internal loop, including several positive charge clusters, also readily formed electrostatic interaction networks with SKP1’s CTR and its BTB domain (Figs. 5H, S11, S15).

### GAT1 Monomerizes SKP1 in a CTR Dependent Manner

Previous studies have suggested that the high kinetic exchange of specific interactions in fuzzy protein complexes can facilitate exchange between interacting macromolecules [20–23]. Therefore, we investigated a possible role of GAT1 in facilitating the exchange of the tight SKP1 homodimer. As shown in Fig. 8A, a substoichiometric concentration of GAT1 was found to dramatically increase the pool of SKP1 monomers at the expense of the dimer pool during the 2-h preincubation period that persisted during separation during centrifugation. The affinity of the full-length, unmodified construct was previously estimated at ∼50 nM [18]. Even at a 20 nM concentration of GAT1 (10 nM in terms of the dimer), a significant shift in the equilibrium of 500 nM SKP1 was observed. This is significant considering that 500 nM is an order of magnitude above SKP1’s *K*_d_, and more significant considering GAT1 binds SKP1 in a strict 2:1 GAT1:SKP1 complex (Fig. 3A). A previous study showed that a version of SKP1 in which the CTR sequence is scrambled (Scrambled6) maintained the high homodimer affinity of the native sequence, in contrast to a version in which the CTR was deleted [18]. These findings indicated the importance of a fuzzy interaction contributed by the CTR to generate the high affinity interaction. To explore a role for the CTR in the GAT1 effect, a concentration series of SKP1ΔCTR in the presence of a 1:10 ratio of GAT1:SKP1 was similarly analyzed (Fig. 8B). The region associated with the SKP1 monomer and dimer species were integrated and deconvoluted into monomer and dimer contributions to yield an apparent *K*_d_ of 900 nM. This matches the homodimer *K*_d_ previously determined for this construct [18] and indicates that SKP1’s exchange toward the monomeric species is dependent on the presence of SKP1’s CTR. In contrast, analysis of Scrambled6 also revealed substantial monomerization of the homodimer (Fig. 8C), supporting a role for the CTR but also suggesting that it acts in a fuzzy manner as in the SKP1 homodimer. This is consistent with the frequent CTR residue contacts with various positions of GAT1 (Figs. 5, S15), which might compete with contacts between SKP1 CTRs. A similar effect was observed with Gly-SKP1 (Fig. 8D). This was, however, less surprising due to the much higher homodimer *K*_d_ of Gly-SKP1 of 400 nM [18], the higher concentration of GAT1 used, and the 2:2 stoichiometry of the GAT1/SKP1 complex (Fig. 3C). Possible explanations for the effect are that GAT1 increases the apparent *K*_d_ of the SKP1 homodimers by stimulating the *k*_off_ rate, or that the nascent monomer possesses a distinct long-lived conformation.

**Figure 8.**
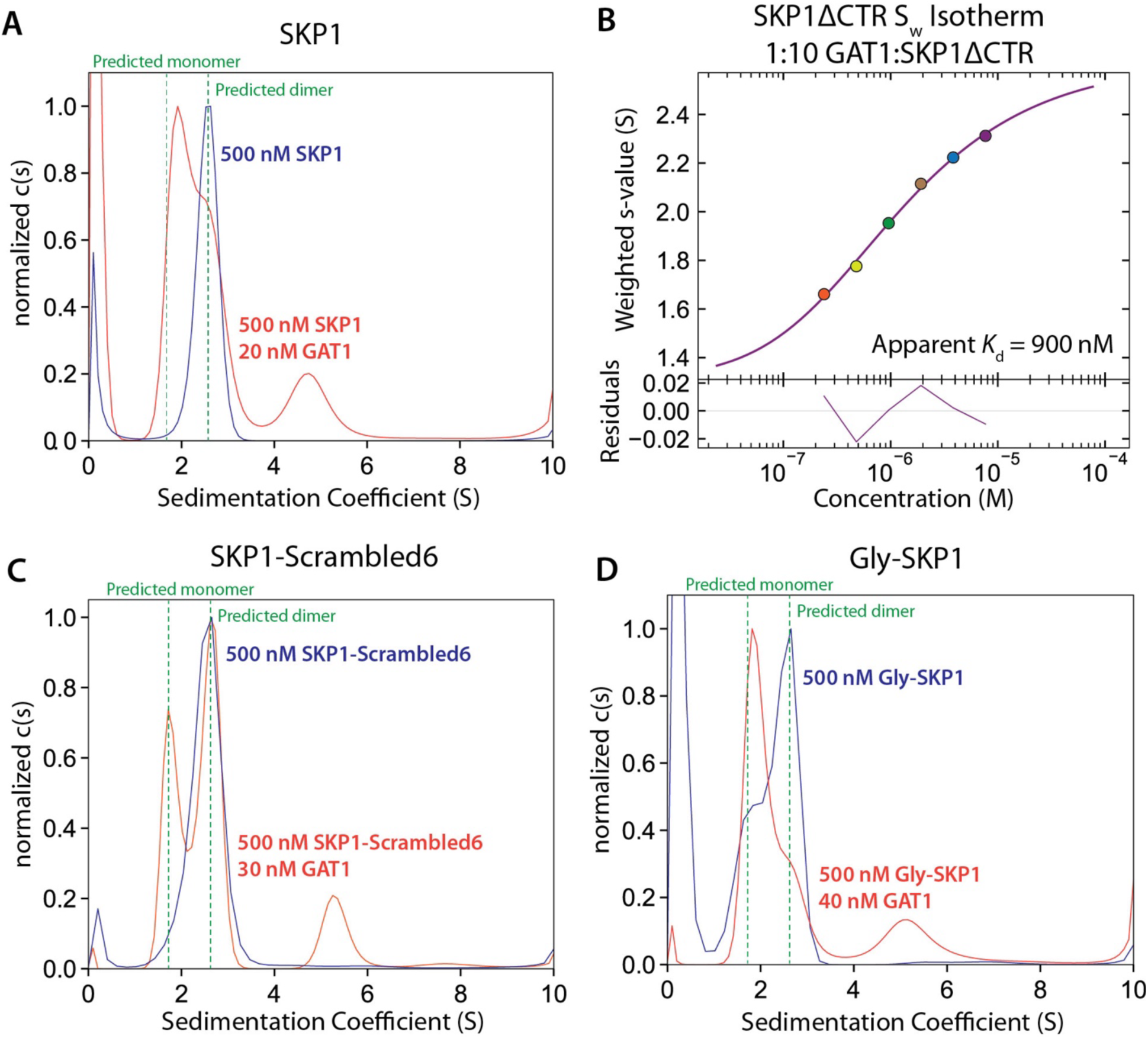
Dissociative Effects of GAT1 on SKP1 Variants. (A) c(s) distributions for 500 nM SKP1 (blue) and 500 nM SKP1 + 20 nM GAT1 (orange). Predicted s-values for the SKP1 monomer and dimer species are shown as vertical dashed lines. See Fig. S3 for concentration series. (B) Isotherm showing the apparent homodimer affinity for SKP1ΔCTR not bound by GAT1 in the presence of a 1:10 ratio of GAT1:SKP1ΔCTR. (C) Same as in (A) using 500 nM SKP1-Scrambled6 and 500 nM SKP1-Scrambled6 + 30 nM GAT1. (D) Same as in (A) using 500 nM Gly-SKP1 and 500 nM Gly-SKP1 + 40 nM GAT1. Supporting data are in Figs. S17-S18.

## DISCUSSION

GAT1 is a conventional sugar nucleotide-dependent, GT-A fold GT that exists as a soluble homodimer in the cytoplasm. GAT1 alone can catalyze the addition of αGal from UDP-Gal to the 3-position of small αGlc-acceptors. However, it is more active toward the GlFGGn-tetrasaccharide that it modifies on SKP1 in cells when coupled to the pNP aglycon and is again substantially more active when GlFGGn-is linked to SKP1 itself [25]. Here we show the structural basis that likely explains the observation that SKP1 is the only natural acceptor for GAT1 in cells. The primary observation is that GAT1 modifies SKP1 monomers despite SKP1’s proclivity to dimerize. Importantly, GAT1 also monomerizes SKP1 dimers even when they are not modified and thus not substrates. We propose this effect tunes the availability of SKP1 in cells as manifested by the altered profile of FBPs in the SKP1-SF interactome of *gat1*-KO cells (Fig. 2).

Computational and mutagenesis studies, together with SV-AUC results, show that the high affinity GAT1 homodimer binds a single unmodified SKP1 monomer via a core interface that buries ∼3300 A^2^ (Fig. 9A). This interface includes α-helices-5 and -6, which overlap with the core SKP1 homodimer interface, and a region including α-helix-2 and a loop up to α-helix-3, which contributes to CUL1 binding (Fig. 1B). Thus, the GAT1 interaction dissolves the SKP1 homodimer but at the same time prevents interactions with FBPs and CUL1. Furthermore, with an estimated 80 nM *K*_d_, the GAT1/SKP1 interaction is expected to be competitive with the SKP1 homodimer whose *K*_d_ was estimated at 50 nM [18]. The intrinsically disordered CTR also contributed to binding GAT1 based on effects of its deletion. Its role appears to involve dynamic charged based interactions with many positions on GAT1 subunits 1 and 2, based on all-atom molecular dynamics simulations regardless of the staring conformation. However, the presence of the tetrasaccharide acceptor, which was previously modeled into the active site of the GAT1 ortholog from *Pythium* [25], is accessible to the active site (Fig. S11) and likely to influence CTR dynamics. A clue may be provided by the 2:2 GAT1:Gly-SKP1 structure detected for the pentasaccharide product of the GAT1 reaction by SV-AUC. In this case separate copies of SKP1 are modeled to symmetrically associate with individual subunits of the GAT1 homodimer. Because glycosylation showed evidence of organizing the CTR [16,17,25], consequent reduced excursions of each CTR may minimize excluded volume effects allowing for dual occupancy (Fig. 9B). There was a roughly 2-fold reduction in the number of productive electrostatic contacts observed when a folded CTR was used as a starting point for MD simulations (e.g., Fig. S16). This correlates with the reduced motility of the CTR of Gly-Skp1 relative to non-glycosylated SKP1 (Fig. S19). This indicates that a bias toward an even partially more well-ordered structure in this region would be expected to weaken an electrostatically mediated fuzzy interaction made with GAT1. The ∼2.5-fold increased *K*_d_ for the 2:2 model (Fig. 3) is consistent with reduced but residual interference. However, the 2:2 ratio allowed for the GAT1/SKP1-Scrambled6 complex (Fig. S18) suggests some sequence dependence for the native CTR interaction. In summary, GAT1 modifies monomeric rather than dimeric SKP1, and its specificity toward GlFGGn-SKP1 is based on an interface also employed by SKP1 to bind itself and FBPs. Significantly, this mechanism limits modification to free SKP1 not complexed with FBPs and influences the monomer/dimer equilibrium of all modification isoforms of SKP1, not just the substrate GlFGGn-SKP1.

**Figure 9.**
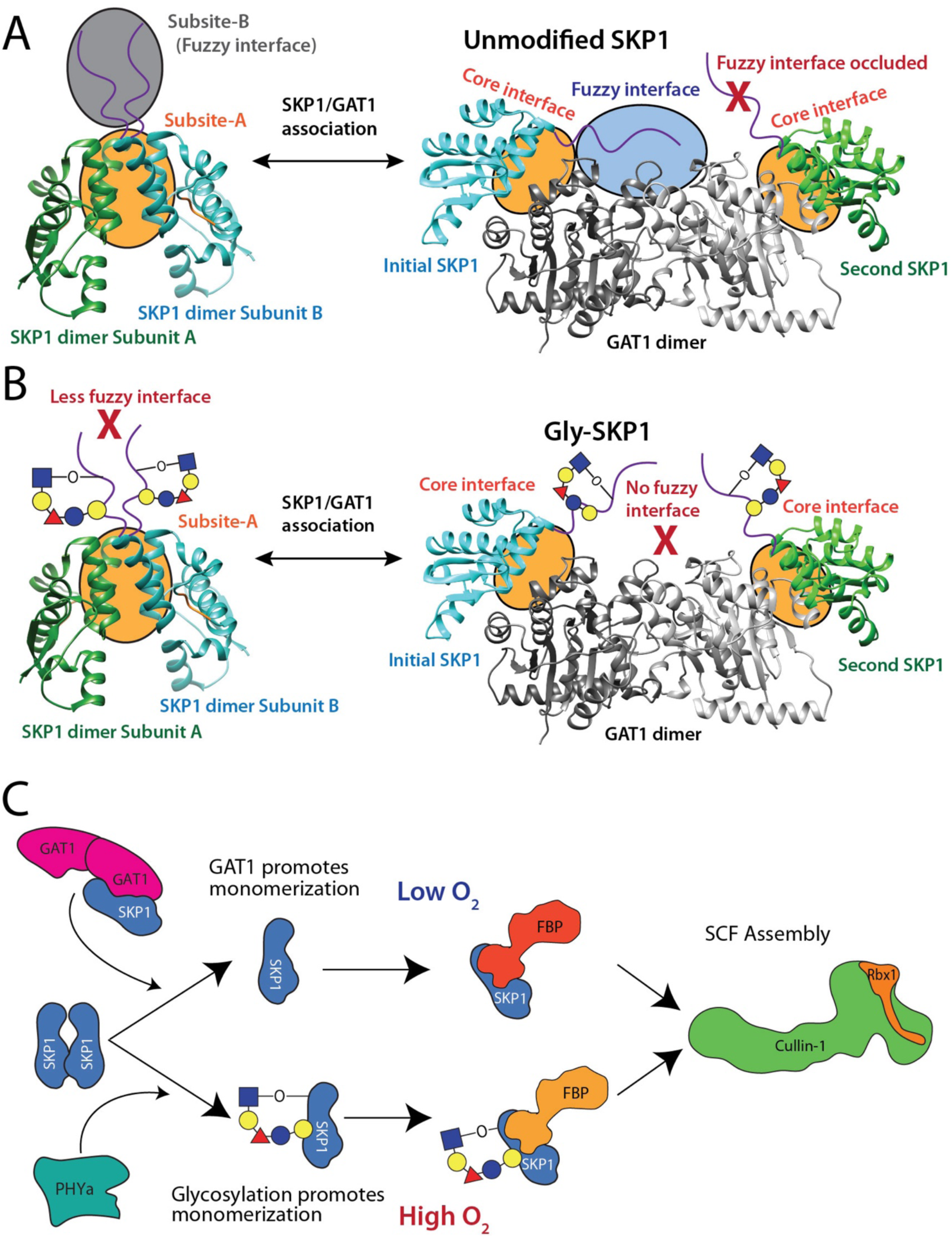
The Non-Enzymatic Role of GAT1 in Binding SKP1. (A) Model for the 2:1 GAT1:SKP1 complex. The SKP1 homodimer forms a two-component interface composed of a core Subsite-A and a fuzzy Subsite-B. These two interfaces exchange with similar interactions occurring in the 2:1 GAT1:SKP1 complex. A second binding event with the other GAT1 subunit is disfavored by excluded volume and competitive binding effects with the intrinsically disordered SKP1 CTR. This 2:2 GAT1:SKP1 complex may, however, act as a transition state allowing for more rapid exchange of SKP1 subunits. (B) Model for the 2:2 GAT1:Gly-SKP1 complex. Glycosylation promotes a disorder to order transition in SKP1’s CTR reducing the fuzzy interactions. The lack of fuzzy interactions between SKP1 and GAT1 allows for symmetrical binding of a second Gly-SKP1 to the other GAT1 subunit. Additionally, the SKP1 homodimer, weakened by the loss of Subsite-B, is less able to effectively compete with the GAT1:SKP1 complex for free SKP1. (C) Model for GAT1’s role as a SKP1 exchange factor. PHYa acts in the presence of sufficient oxygen to glycosylate SKP1 and weaken SKP1 homodimerization by disrupting Subsite-B, allowing free SKP1 to more readily engage with FBPs. Under low oxygen conditions, GAT1 acts as an exchange factor promoting formation of monomeric SKP1. This enhances a free SKP1 monomer pool even in the absence of glycosylation. Note, however, the expectation that Gly-SKP1 and unmodified SKP1 may have distinct FBP preferences as suggested by differential shading.

Surprisingly, there was a persistence of monomeric SKP1 in the presence of substoichiometric GAT1, even at 500 nM SKP1, far above its ∼50 nM *K*_d_ (Fig. 8). This suggests a novel model in which the interacting partners have the following characteristics. i) The GAT1 homodimer initially encounters the SKP1 homodimer rather than free SKP1 monomers that are in equilibrium with the dimers. ii) The SKP1 homodimer achieves its low *K*_d_ value due to relatively slow *k*_on_ and *k*_off_ rate constants. iii) The *K*_d_ that describes the GAT1/SKP1 interaction (84 nM) (Fig. 3) is characterized by a ratio of fast *k*_off_ and *k*_on_ rate constants. With these assumptions, GAT1 might increase the SKP1 monomer pool by either increasing SKP1’s homodimer *k*_off_ or decreasing its *k*_on_. The former might occur as a result of transient interactions of the two SKP1 subunits with the GAT1 homodimer, i.e, a 2:2 complex transition state that rapidly dissociates due to repulsion of their CTRs. The proposed rapid kinetics of the GAT1/SKP1 complex relative to that of SKP1 homodimerization could explain how substoichiometric GAT1 promotes monomerization of a >10-fold molar excess of SKP1. As a result, a pool of monomeric SKP1 would become available to bind other high affinity partners such as FBPs. This hypothesized kinetic exchange model warrants further studies.

Previous findings indicate that fuzzy protein interactions can facilitate the rapid exchange of high affinity binding partners, as well as allow for more facile addition of post translational modifications [20–23]. Given that GAT1’s catalytic effect on SKP1 homodimerization depends on the presence of its CTR (Fig. 8), GAT1 may leverage this property of fuzzy interactions to exchange SKP1/SKP1 fuzzy interactions with SKP1/GAT1 interactions, and thus act like a SKP1 exchange factor. The effect would be less significant for Gly-SKP1 because, with a *K*_d_ almost an order of magnitude greater than that of SKP1, the monomer is already more abundant. However, the lower apparent affinities of the two Gly-SKP1’s to the GAT1 dimer (Fig. 3B), perhaps associated with mutual interference between their CTRs, may increase the monomer pool (Fig. 8). If there is an affinity hierarchy of FBPs, owing to distinct F-box sequences, for limiting amounts of the single SKP1, subtle changes in SKP1 availability may affect the profile of FBP/SKP1 complexes *in vivo* [12].

The influence of GAT1 on TgSKP1 shows parallels with the previously reported influence of the terminal GT AgtA in *Dictyostelium* to act as a buffer for SKP1 also via a sequestration mechanism [64]. AgtA also binds strongly to unmodified Skp1 *in vitro*. *In vivo*, the *agtA*-KO is much more deleterious for the O_2_-dependence of culmination than expected based on comparison with effects of KOs of preceding GTs in the modification pathway, a double-KO with *gnt1* is more deleterious than the *gnt1*-KO, and overexpression of catalytically dead mutant AgtA inhibited Skp1 modification and terminal differentiation [reviewed in 12]. Strikingly, AgtA is evolutionarily unrelated and its effect on *Dictyostelium* Skp1 involves a separate WD40-repeat domain expected to fold as a β-propeller. This mechanism may ensure specificity of AgtA for FGGn-Skp1 relative to the abundant O-Fuc residues on other proteins in the cytoplasm [69]. However, the relative influence is distinct given the considerably higher *K*_d_ (∼2 µM) for the *Dictyostelium* SKP1 homodimer [52]. Through apparent adaptations of their substrate binding activities, these two terminal GTs appear to have evolved related new secondary roles proposed to regulate the availability of their SKP1 substrates and non-substrate isoforms for FBP interactions.

In summary, GAT1 may act to complement or buffer the action of PHYa on SKP1 monomer generation through non-enzymatic means (Fig. 9C). Homodimerization competes with FBPs to form functional 1:1 FBP:SKP1 subcomplexes [52]. Under normoxic conditions, PHYa will efficiently hydroxylate SKP1 allowing for its glycosylation. Upon glycosylation, the SKP1 homodimer will substantially weaken to free more SKP1 for incorporation into FBP complexes [18]. Under these conditions, GAT1 may tune SKP1 availability itself through dual binding events on the GAT1 homodimer. Under hypoxic conditions, however, PHYa will not as efficiently hydroxylate SKP1 resulting in reduced glycosylation. Unmodified SKP1 forms a tight homodimer that can effectively compete with FBP binding [18]. Under these conditions, GAT1 is proposed to access the SKP1 homodimer and act as an exchange factor to promote the transient accumulation of monomeric SKP1. The resulting monomeric SKP1 will then be available for FBP binding allowing for sufficient SCF activity even under low oxygen conditions. Evidence of physiological significance comes from an effect of the loss of GAT1 on the SKP1 interactome. Whereas some Group 2 FBPs show intermediate abundances in *gat1*-KO parasites relative to *phyA*-KO and normal (RHΔΔ) parasites as previously observed for *pgtA*-KO parasites (Fig. 2), some Group 1 FBPs, whose presence in the SKP1 interactome of normal parasites was not affected in *phyA*-, *gnt1*- and *pgtA*-KO parasites, showed highly enriched presence in *gat1*-KO parasites. One interpretation is that in the absence of the monomerization activity of GAT1, Group 3 FBPs have an advantage in accessing SKP1 owing to intrinsically high affinity for SKP1 that can compete with its homodimer [12]. However, further studies are required to rule out an alternative interpretation that Group 3 FBPs simply have a relatively strong preference for the tetrasaccharide (GlFGGn-) glycoform of SKP1 compared to other FBPs.

## Supporting information

Supplement, full

## ACKNOWLEDGMENTS

We are grateful to Dr. Msano Mandalasi for generating the *gat1*-KO/SKP1-SF strain, and Dr. Hyunwoo Kim and Dr. Rob Woods for their suggestions.

## DATA AVAILABILITY

The MS proteomics data are deposited in the ProteomeXchange Consortium via the PRIDE [99] partner repository with the dataset identifiers PXD079463 and PXD079410.

## SUPPORTING INFORMATION

This article contains supporting information, with 19 figures.

## ABBREVIATIONS

BSA: bovine serum albumin
co-IP: co-immunoprecipitation
CTR: partially intrinsically disordered C-terminal region of SKP1
FBP: F-box protein
GGlFGGn-SKP1: notation that identifies the individual monosaccharides of the SKP1 glycan, including galactose (G), glucose (Gl), fucose (F), N-acetylglucosamine (Gn)
Gly-SKP1: abbreviated notation for fully glycosylated SKP1 (GGlFGGn-Skp1)
GT: glycosyltransferase
KO: knock-out or inactivation of the protein coding capacity of a gene; MD, molecular dynamics simulations
MMGBSA: Molecular Mechanics/Generalized Born Surface Area
MS: mass spectrometry
pNP: paranitrophenol
SV-AUC: sedimentation velocity analytical ultracentrifugation

