## Supplement, full for "Potential for the Terminal SKP1 Glycosyltransferase to Exert Non-Enzymatic Control of SKP1 in Toxoplasma gondii"

### Table of Contents

Figure S1. Replacement of the *gat1* coding region with a DHFR resistance cassette in a SKP1-SF strain
Figure S2. Sequences of expressed GAT1 variants
Figure S3. Residuals and c(s) distributions for GAT1:SKP1 association experiments
Figure S4. Residuals and c(s) distributions for GAT1:Gly-SKP1 association experiments
Figure S5. Residuals and c(s) distributions for GAT1:SKP1ΔCTR association experiments
Figure S6. Overlay of distributions utilized for Isotherms in Figure 3
Figure S7. Alignment of GAT1:SKP1 AlphaFold3 with the *Pythium* GAT1 crystal structure
Figure S8. pLDDT scores for the 2:1 GAT1:SKP1 complex
Figure S9. Competition assays between GAT1 and Fbs1 for limiting SKP1 concentrations
Figure S10. Inhibition of PHYa in the presence of varying concentrations of GAT1
Figure S11. Overlay of an AlphaFold2 structure of a 2:1 GAT1:SKP1 complex with a glycan docked PuGAT1 homodimer structure
Figure S12. Per residue MMGBSA scores for GAT1/SKP1 complexes
Figure S13. Predicted changes in binding energy of GAT1 to SKP1
Figure S14. Predicted change in stability of the GAT1 homodimer for each cluster
Figure S15. pLDDT scores for AlphaFold3 models of mutant GAT1/SKP1 complexes
Figure S16. A sample of distances between oppositely charged residues between SKP1 and GAT1
Figure S17. Residuals and c(s) distributions for SKP1ΔCTR homodimerization with GAT1
Figure S18. Residuals and c(s) distributions for SKP1-Scrambled6 with GAT1
Figure S19. Mobility of residues in complex with glycosylated and unmodified SKP1

**Figure S1.** Replacement of the *gat1* coding region with a DHFR resistance cassette in a SKP1-SF strain. As previously described [25], dual guide RNAs (gRNA) were expressed by transient transfection with a plasmid that also encoded Cas9, and a repair DNA encoding DHFR. Clones were screened by PCR using primers Fh and Rh that hybridize to sequences flanking the gRNA sites and DHFR homology arms. Four clones indicate replacement of the locus with the longer DHFR-resistance cassette, and clone E8 (strain MM18) was chosen for analysis.

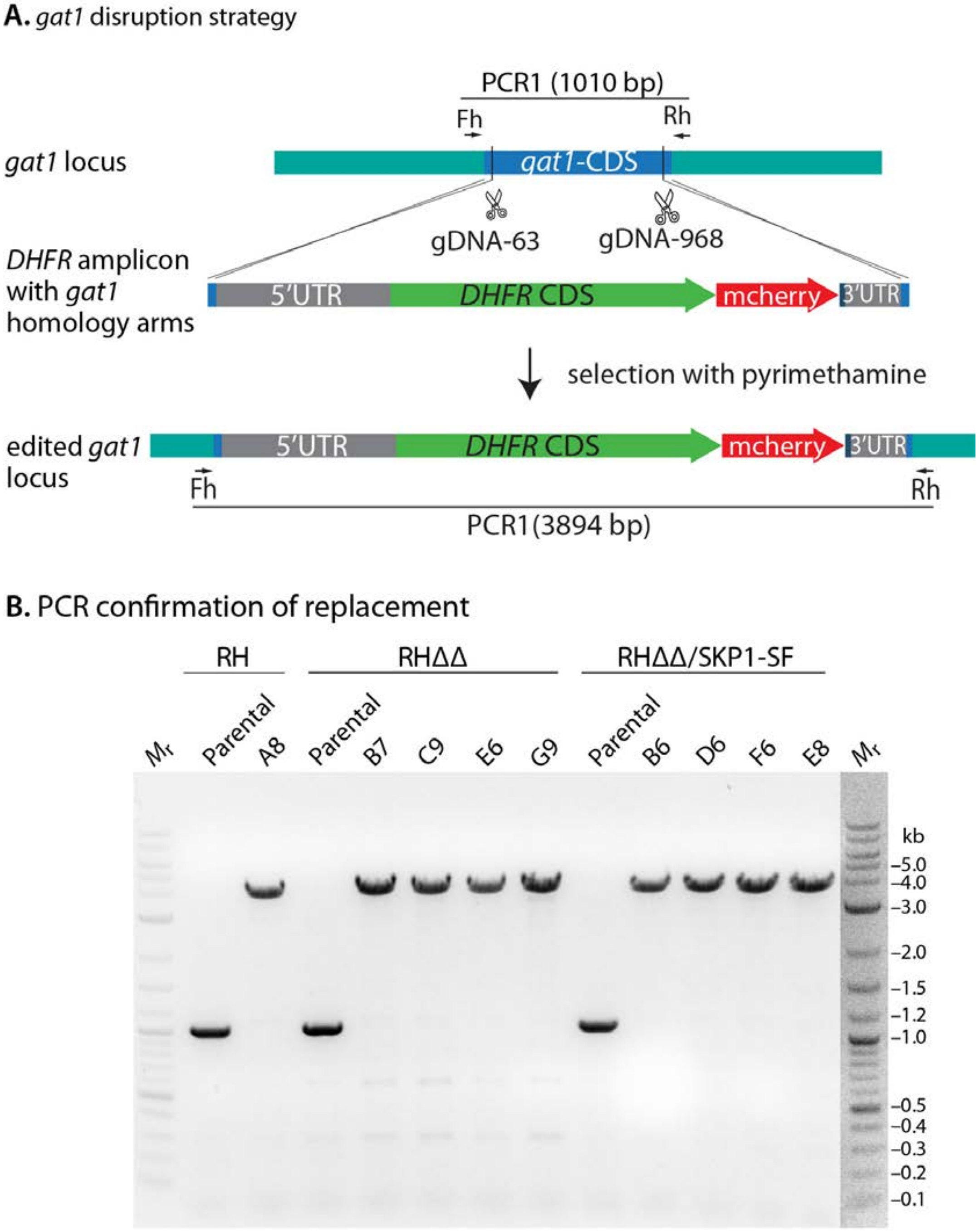

**Figure S2.** Sequences of expressed GAT1 variants. The pET15-HisTEVTgGat1 plasmid [18] was used as the template for site-directed mutagenesis. (A) Primer pairs used to introduce the Y18K and A51K mutations into GAT1. (B) The coding region for the TgGat1 gene used to express GAT1 in *E. coli*. The mutated DNA and aa sequences are given in purple. The $His_6$-tag is highlighted green, the TEV protease recognition site is highlighted yellow, the cleavage site is highlighted blue, and mutated residues/codons are highlighted in grey.

**A.**

| Primer | Sequence |
|---|---|
| Y18K-F | 5′-TTCTTTCTACAAGGGTGTCGAGGCACTGCTCA-3′ |
| Y18K-R | 5′-ACCCTTGTAGAAAGAATTGTCCGTCAACAGGGT-3′ |
| A51K-F | 5′-AATAAAAAAGTTGGTTTATCAGCGTCGAAAAGC-3′ |
| A51K-R | 5′-AACCAACTTTTTTATTGTACTCTGAGAAACATCAGATGTGTG-3′ |

His tag
Cleavage site
TEV protease recognition site
Y18K-A51K
Mutated residues

**B.**

```
>pET15-HisTEVTgGat1
TAATACGACTCACTATAGGGGAATTGTGAGCGGATAACAATTCCCCTCTAGAAATAATTTTGTTTAACTTTAAGA
AGGAGATATACC
M  G  S  S  H  H  H  H  H  H  S  S  G  R  E  N  L  Y  F  Q><
ATGGGCAGCAGCCATCATCATCATCATCACAGCAGCGGCAGAGAAAACTTGTATTTCCAG

G  H  M  A  S  M  S  P  R  Y  A  Y  A  T  L  L  T  D  N  S
GGCCATATGGCTAGCATGTCTCCTCGGTACGCGTACGCTACCCTGTTGACGGACAATTCT
                                                   5′-TTCT (Y18K-F)
                                     3′-TGGGACAACTGCCTGTTAAGA (Y18K-R)

F  Y  Y  G  V  E  A  L  L  K  S  L  E  A  T  K  T  P  Y  P
TTCTACTATGGTGTCGAGGCACTGCTCAAGTCACTGGAGGCTACGAAGACGCCTTACCCC
F  Y  K  G  V  E  A  L  L  K  S  L  E  A  T  K  T  P  Y  P
TTCTACAAGGGTGTCGAGGCACTGCTCAAGTCACTGGAGGCTACGAAGACGCCTTACCCC
TTCTACAAGGGTGTCGAGGCACTGCTCA-3′
AAGATGTTCCCA-5′

V  L  L  L  H  T  S  D  V  S  Q  S  T  I  K  A  L  V  Y  Q
GTGCTTCTTTTGCACACATCTGATGTTTCTCAGAGTACAATAAAAGCGTTGGTTTATCAG
V  L  L  L  H  T  S  D  V  S  Q  S  T  I  K  K  L  V  Y  Q
GTGCTTCTTTTGCACACATCTGATGTTTCTCAGAGTACAATAAAAAAGTTGGTTTATCAG
                                  5′-AATAAAAAAGTTGGTTTATCAG (A51K-F)
      3′-GTGTGTAGACTACAAAGAGTCTCATGTTATTTTTTCAACCAA-5′     (A51K-R)

R  R  K  A  P  A  S  E  D  A  G  T  T  G  K  E  M  K  T  G
CGTCGAAAAGCCCCGGCGAGTGAGGATGCGGGAACTACAGGGAAGGAAATGAAAACAGGG
CGTCGAAAAGC-3′

Q  E  V  I  P  S  S  Q  C  P  E  H  T  P  G  R  N  L  H  S
CAGGAAGTCATCCCAAGTTCACAGTGTCCAGAACACACCCCAGGTAGAAACTTGCACTCC

P  I  G  R  K  G  V  N  P  V  S  C  S  V  T  Q  D  E  T  R
```

CCCATTGGCAGGAAAGGGGTAAACCCTGTGAGTTGCTCCGTCACACAAGACGAGACTAGG

**V R T D S D H I E E A E R R A S A R T S**
GTTCGTACTGATTCAGATCATATAGAAGAAGCAGAGCGTCGAGCCTCAGCGAGAACCTCG

**E R A R A G G T E E Q G I C V I P R L V**
GAGCGAGCGAGAGCTGGGGGAACAGAGGAACAGGGCATTTGCGTTATTCCCCGACTCGTT

**G S V A Y P K A E R D T C P V E G W K D**
GGTTCTGTCGCGTACCCTAAAGCGGAACGGGACACGTGCCCTGTTGAAGGGTGGAAGGAC

**C F T K L R V W E Q V D F D V I V Y V D**
TGTTTCACCAAACTGCGTGTGTGGGAGCAGGTTGACTTCGATGTGATTGTGTATGTCGAC

**A D C I V L R P V D E L F L R Q P L P A**
GCGGACTGTATAGTTTTGCGGCCGGTAGACGAGCTTTTTCTTAGGCAGCCACTACCCGCC

**F A P D I F P P D K F N A G V A V L K P**
TTTGCACCAGATATCTTCCCTCCCGATAAATTTAACGCGGGAGTCGCAGTGCTGAAGCCC

**D L G E Y G N M V A A V E R L P S Y D G**
GACCTCGGCGAATACGGAAATATGGTAGCCGCGGTCGAGCGTTTACCTTCATATGACGGA

**G D T G F L N A Y F S S W Y E N A A G A**
GGCGACACAGGGTTTTTGAACGCGTATTTCTCATCGTGGTATGAAAACGCCGCTGGCGCC

**R L P F R Y N A L R T L Y H M T Y S S R**
CGTTTGCCCTTTCGGTACAATGCTCTGCGCACACTGTATCACATGACGTACTCCAGTCGA

**K G Y W D A V K P I K I L H F C S S P K**
AAAGGATACTGGGATGCCGTCAAGCCGATCAAAATCCTGCACTTCTGCTCCTCCCCGAAG

**P W E Q P A K T D L E E L W W K V F L T**
CCTTGGGAACAACCAGCAAAGACCGACCTCGAGGAACTATGGTGGAAAGTCTTCCTTACG

**G T V P T D S D I V ***
GGCACTGTGCCAACTGATTCTGATATCGTGTAGGGATCCTAATAACTAAGTAAACTAGTG

CTGAGCAATAACTAGCATAACCCCTTGGGGCCTCTAAACGGGTCTTGAGGGGTTTTTTG

**Figure S3.** Residuals and c(s) distributions for GAT1/SKP1 association experiments. c(s) distributions and associated sedimentation data used in modeling the association of GAT1 and SKP1. Sedfit models the SKP1 monomer/dimer equilibrium with either two discrete species or a single averaged peak depending on the replicate and Gat1 concentration. The data support Fig. 3A.

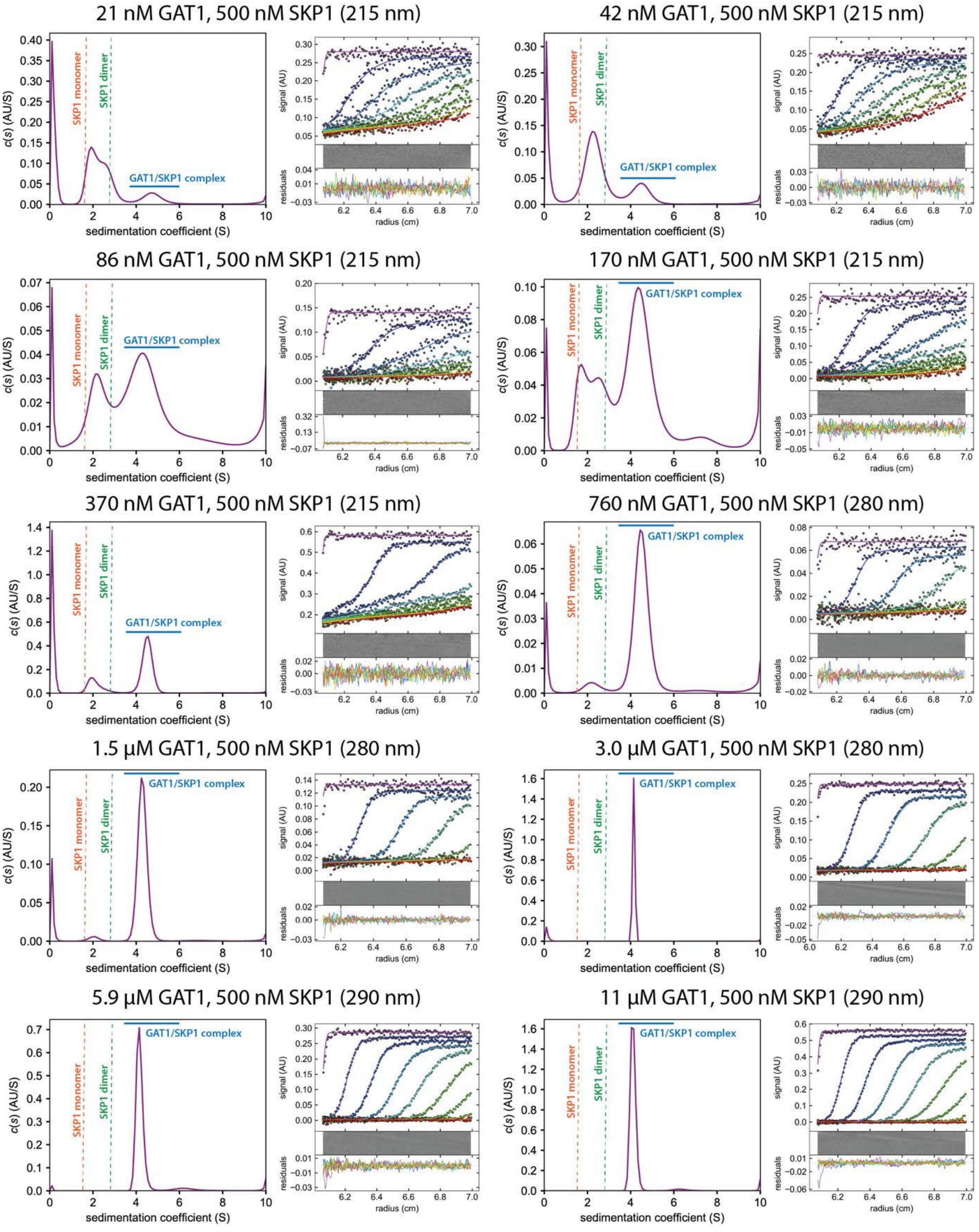

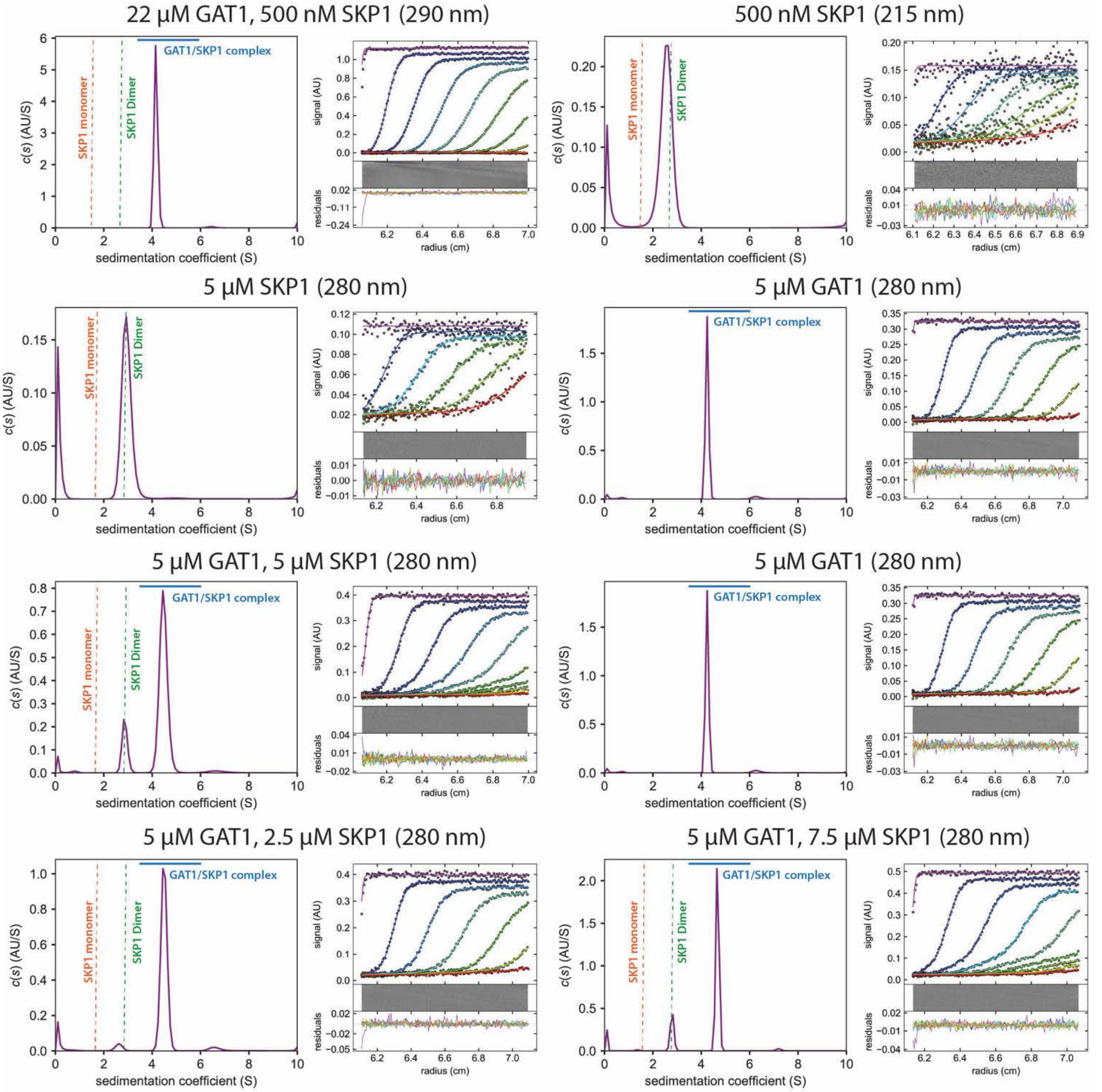
22 µM GAT1, 500 nM SKP1 (290 nm)
500 nM SKP1 (215 nm)
5 µM SKP1 (280 nm)
5 µM GAT1 (280 nm)
5 µM GAT1, 5 µM SKP1 (280 nm)
5 µM GAT1 (280 nm)
5 µM GAT1, 2.5 µM SKP1 (280 nm)
5 µM GAT1, 7.5 µM SKP1 (280 nm)
SKP1 monomer
SKP1 Dimer
GAT1/SKP1 complex
c(s) (AU/S)
sedimentation coefficient (S)
signal (AU)
residuals
radius (cm)

**Figure S4.** Residuals and c(s) distributions for GAT1/Gly-SKP1 association experiments. c(s) distributions and associated sedimentation data used in modeling the association of GAT1 and Gly-SKP1. The 420 nM GAT1 data point is missing due to sample aggregation. The data support Fig. 3B.

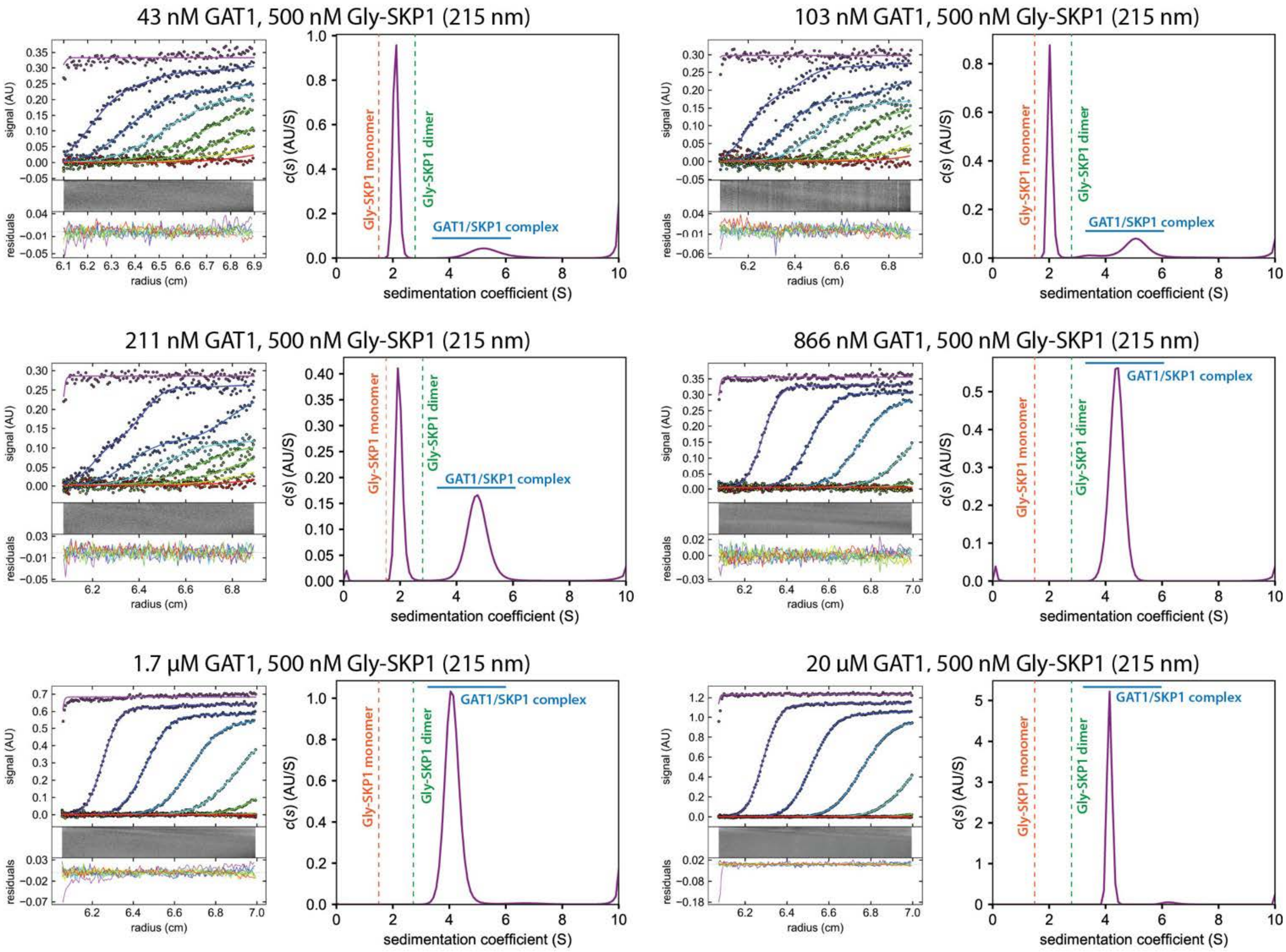

**Figure S5.** Residuals and c(s) distributions for GAT1/SKP1ΔCTR association experiments. c(s) distributions and associated sedimentation data used in modeling the association of GAT1 and SKP1ΔCTR. The data support Fig. 3C.

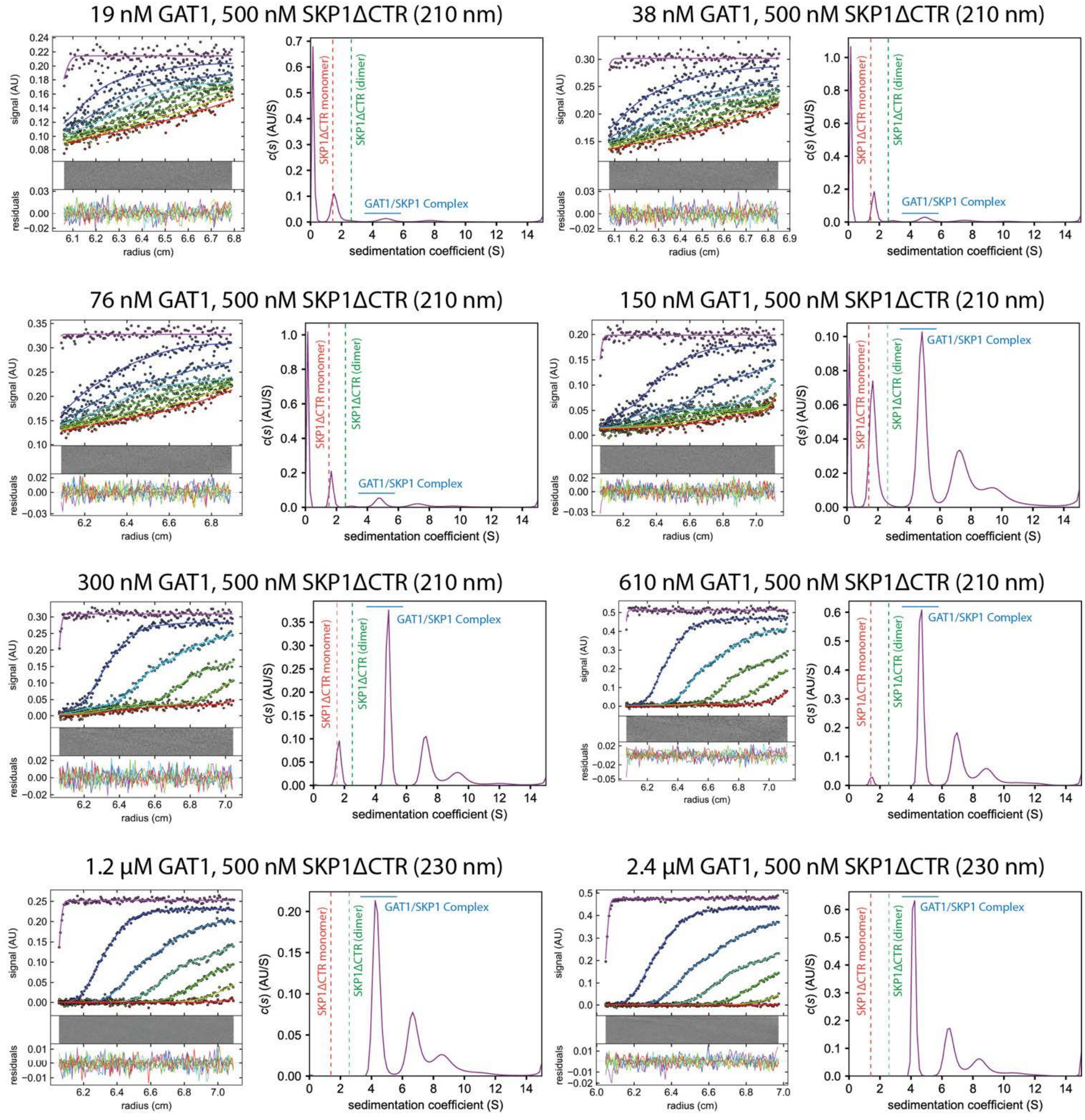

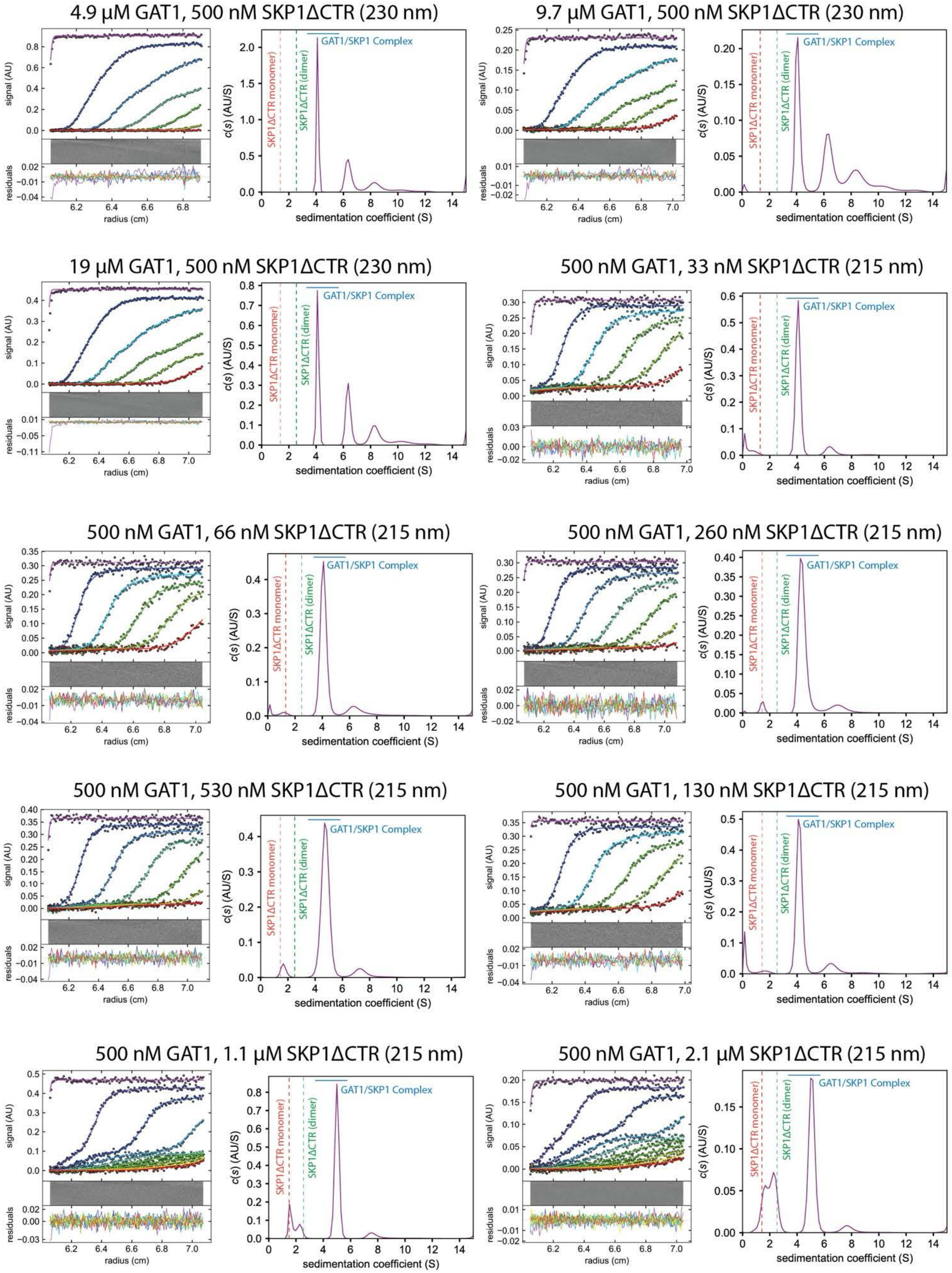
4.9 μM GAT1, 500 nM SKP1ΔCTR (230 nm)
9.7 μM GAT1, 500 nM SKP1ΔCTR (230 nm)
19 μM GAT1, 500 nM SKP1ΔCTR (230 nm)
500 nM GAT1, 33 nM SKP1ΔCTR (215 nm)
500 nM GAT1, 66 nM SKP1ΔCTR (215 nm)
500 nM GAT1, 260 nM SKP1ΔCTR (215 nm)
500 nM GAT1, 530 nM SKP1ΔCTR (215 nm)
500 nM GAT1, 130 nM SKP1ΔCTR (215 nm)
500 nM GAT1, 1.1 μM SKP1ΔCTR (215 nm)
500 nM GAT1, 2.1 μM SKP1ΔCTR (215 nm)
signal (AU)
residuals
radius (cm)
c(s) (AU/S)
sedimentation coefficient (S)
SKP1ΔCTR monomer
SKP1ΔCTR (dimer)
GAT1/SKP1 Complex

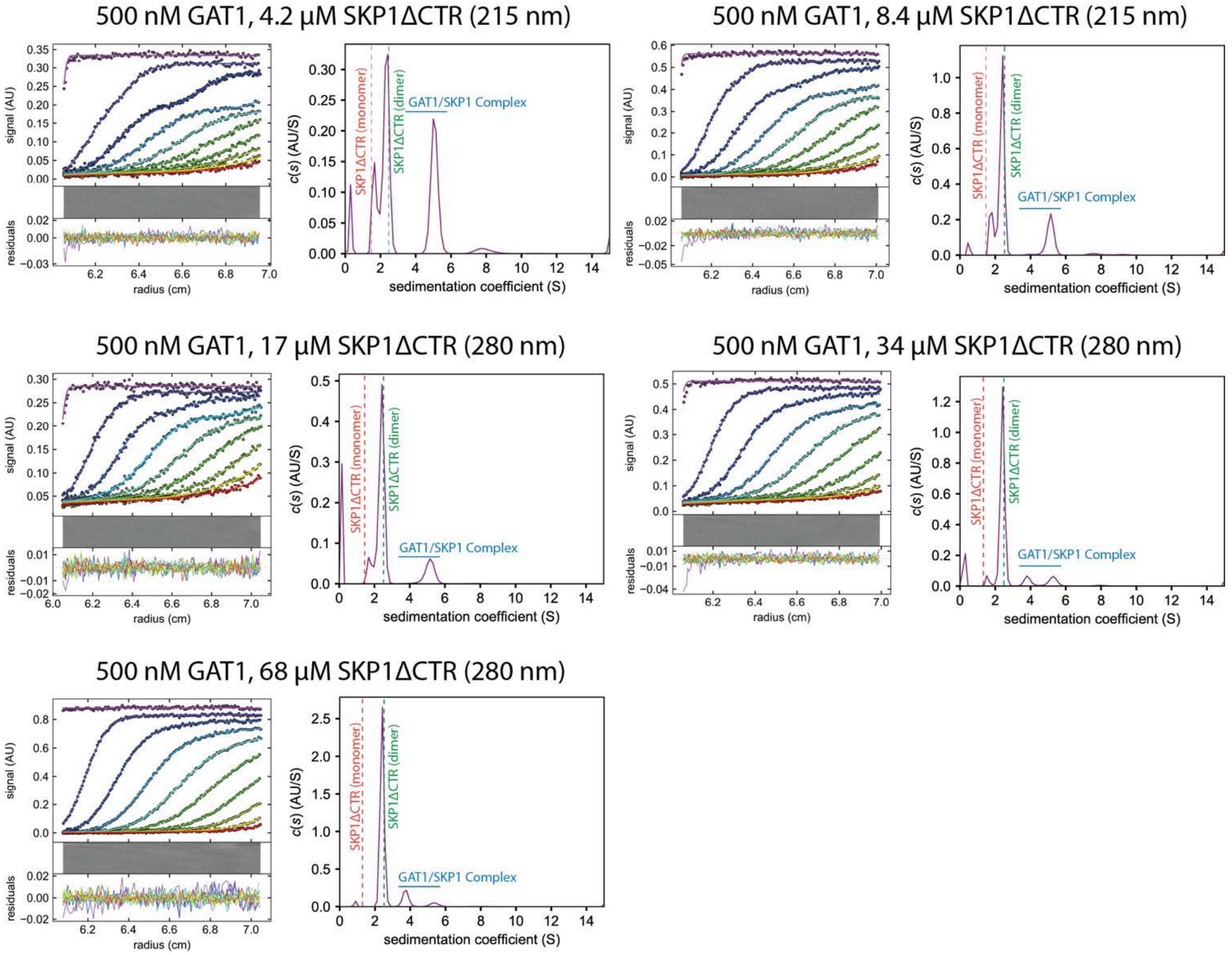
500 nM GAT1, 4.2 μM SKP1ΔCTR (215 nm)
500 nM GAT1, 8.4 μM SKP1ΔCTR (215 nm)
500 nM GAT1, 17 μM SKP1ΔCTR (280 nm)
500 nM GAT1, 34 μM SKP1ΔCTR (280 nm)
500 nM GAT1, 68 μM SKP1ΔCTR (280 nm)
signal (AU)
residuals
radius (cm)
c(s) (AU/S)
sedimentation coefficient (S)
SKP1ΔCTR (monomer)
SKP1ΔCTR (dimer)
GAT1/SKP1 Complex

**Figure S6.** Overlay of Distributions Utilized for Isotherms in Figure 3. Overlay of c(s) distributions used in modeling the association of GAT1 and SKP1 variant isotherms. The SKP1 and Complex peaks were separately integrated, normalized by extinction coefficients, and plotted into a c(s) isotherm. The integration of each complex peak with a constant SKP1 concentration was used in modeling the stoichiometry of each complex. The data support Fig. 3.

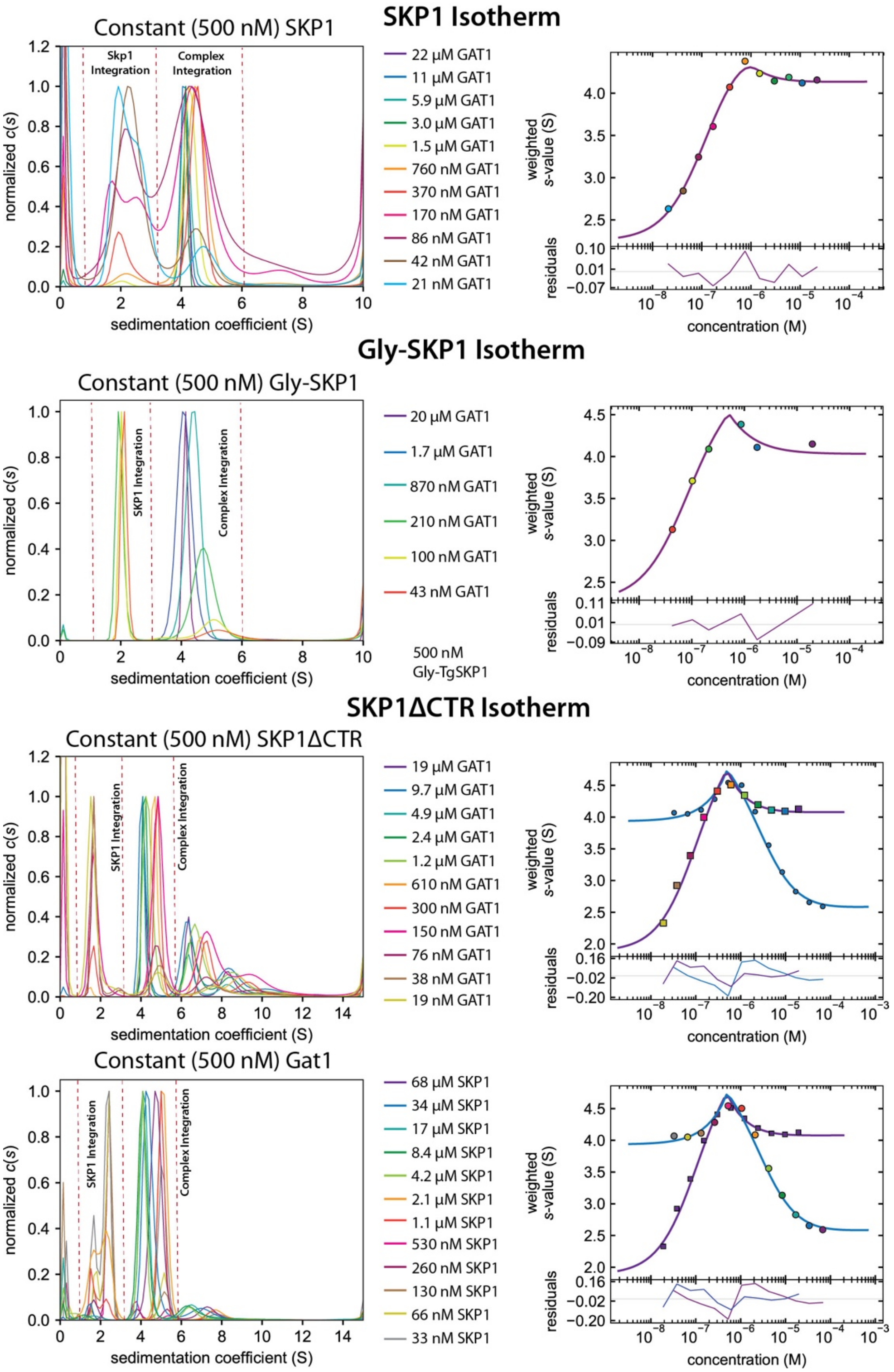

**Figure S7.** Alignment of the *Toxoplasma* GAT1/SKP1 AlphaFold3 structure with the *Pythium* GAT1 crystal structure. Overlay of the top scoring 2:1 GAT1:SKP1 complex backbone trace from AlphaFold3 with that of the PuGAT1 crystal structure (PDB: 6MW8).

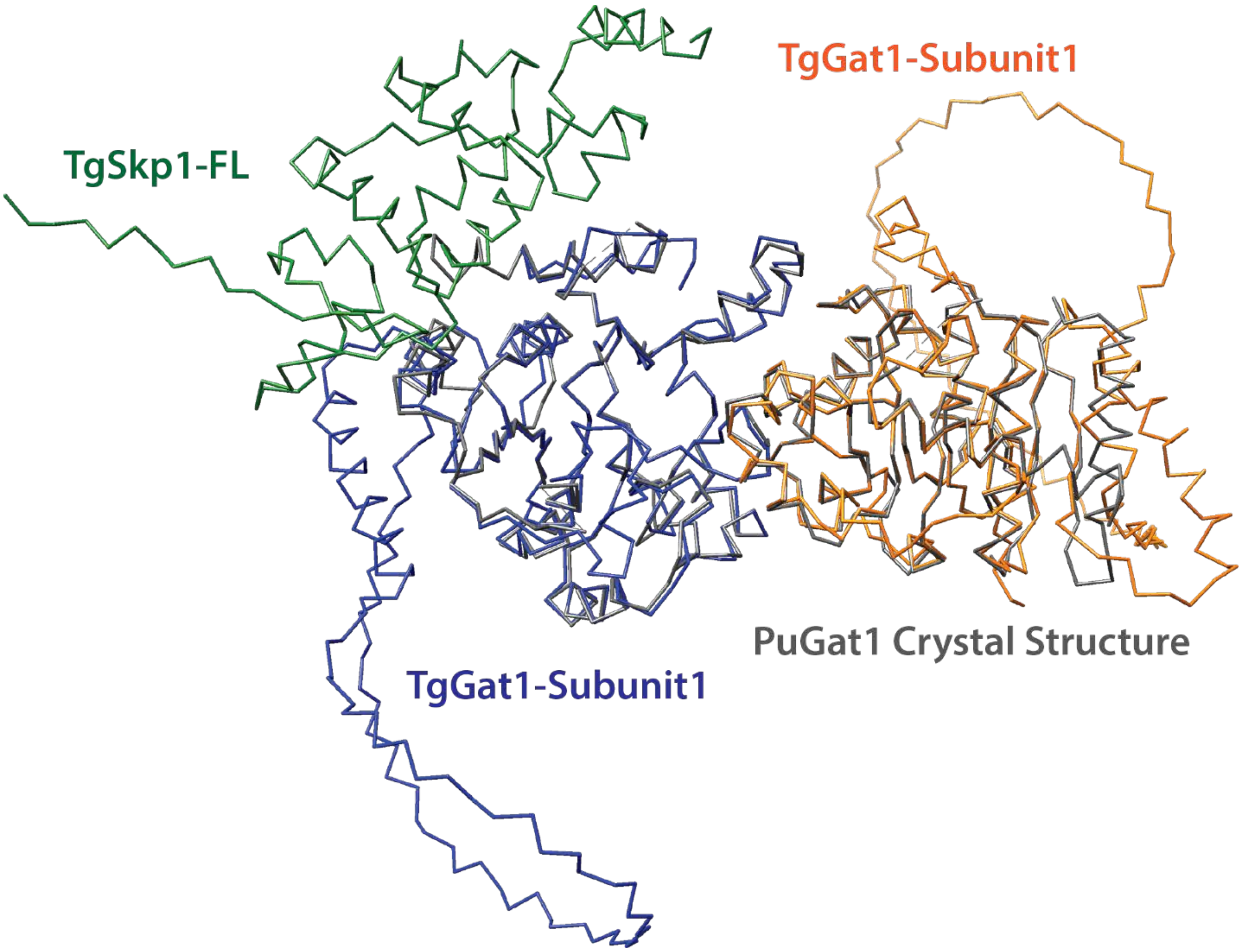

**Figure S8.** pLDDT scores for the 2:1 GAT1:SKP1 complex. Average per residue pLDDT score for each AlphaFold3 model for the 2:1 GAT1:SKP1 complex. These scores were averaged across all 5 structures generated by AlphaFold3 and plotted onto the top scoring model.

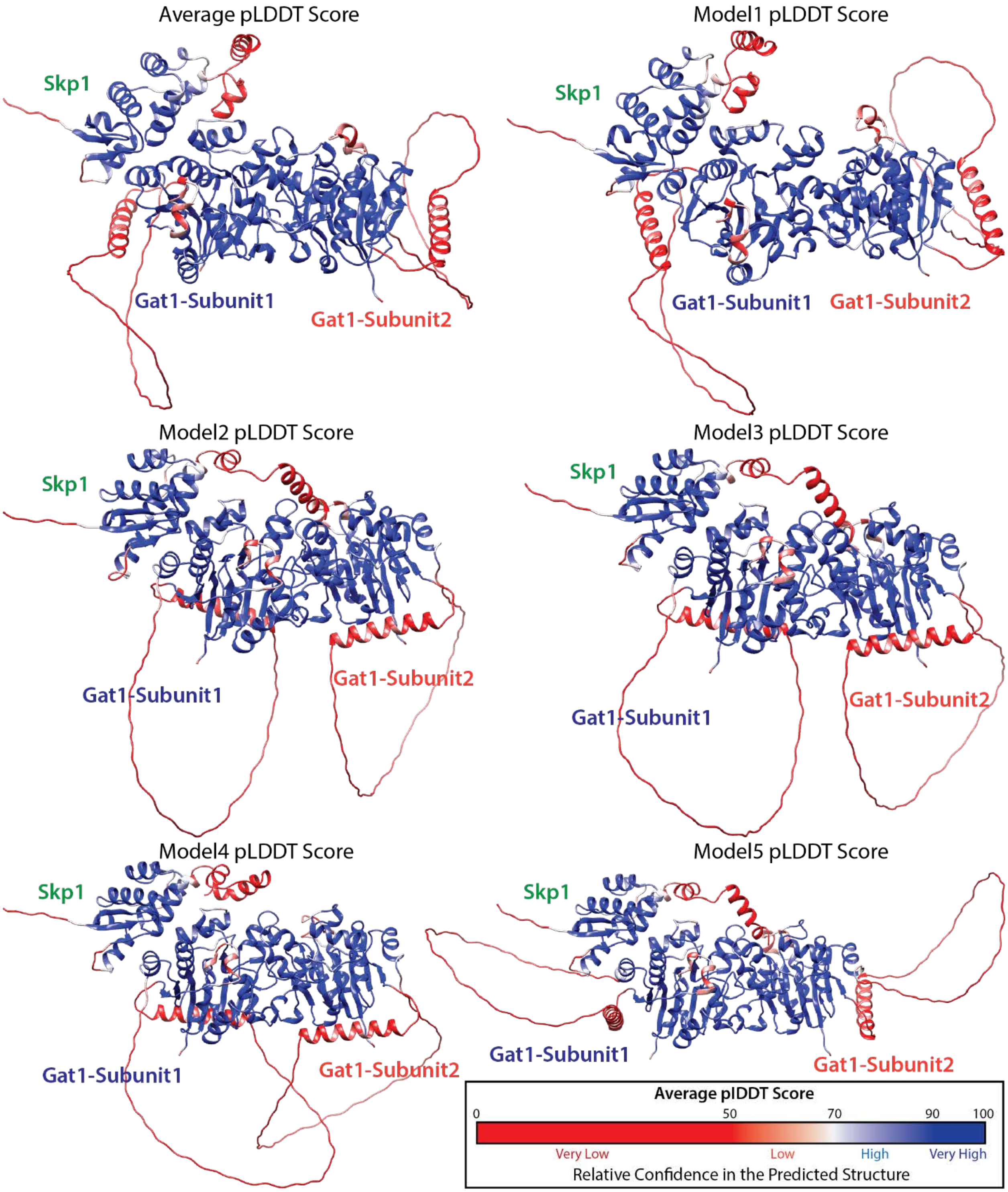

**Figure S9.** Competition assays between GAT1 and Fbs1 for limiting SKP1 concentrations. (A) The indicated mixtures of 10 µM GAT1, 5 µM Fbs1 and/or 5 µM SKP1 were applied to an S200 size exclusion column. Elution was monitored as $A_{280}$. (B) Fractions were analyzed by SDS-PAGE and Coomassie blue staining, and by Western blotting using pAb UOK75 for SKP1 and anti-His Ab for Fbs1. (C) densitometry was used to observe the elution profiles of GAT1 and Fbs1. Both Fbs1 and GAT1 showed an earlier elution profile in the presence of SKP1. When GAT1 and Fbs1 are mixed with limiting SKP1, Fbs1 shows an elution profile matching that of when Fbs1 is mixed with SKP1 alone. In the same condition, GAT1's elution profile matches that of GAT1 in the absence of SKP1. This supports a conclusion where GAT1 is excluded from a complex with SKP1 in the presence of Fbs1 in line with a competitive binding model.

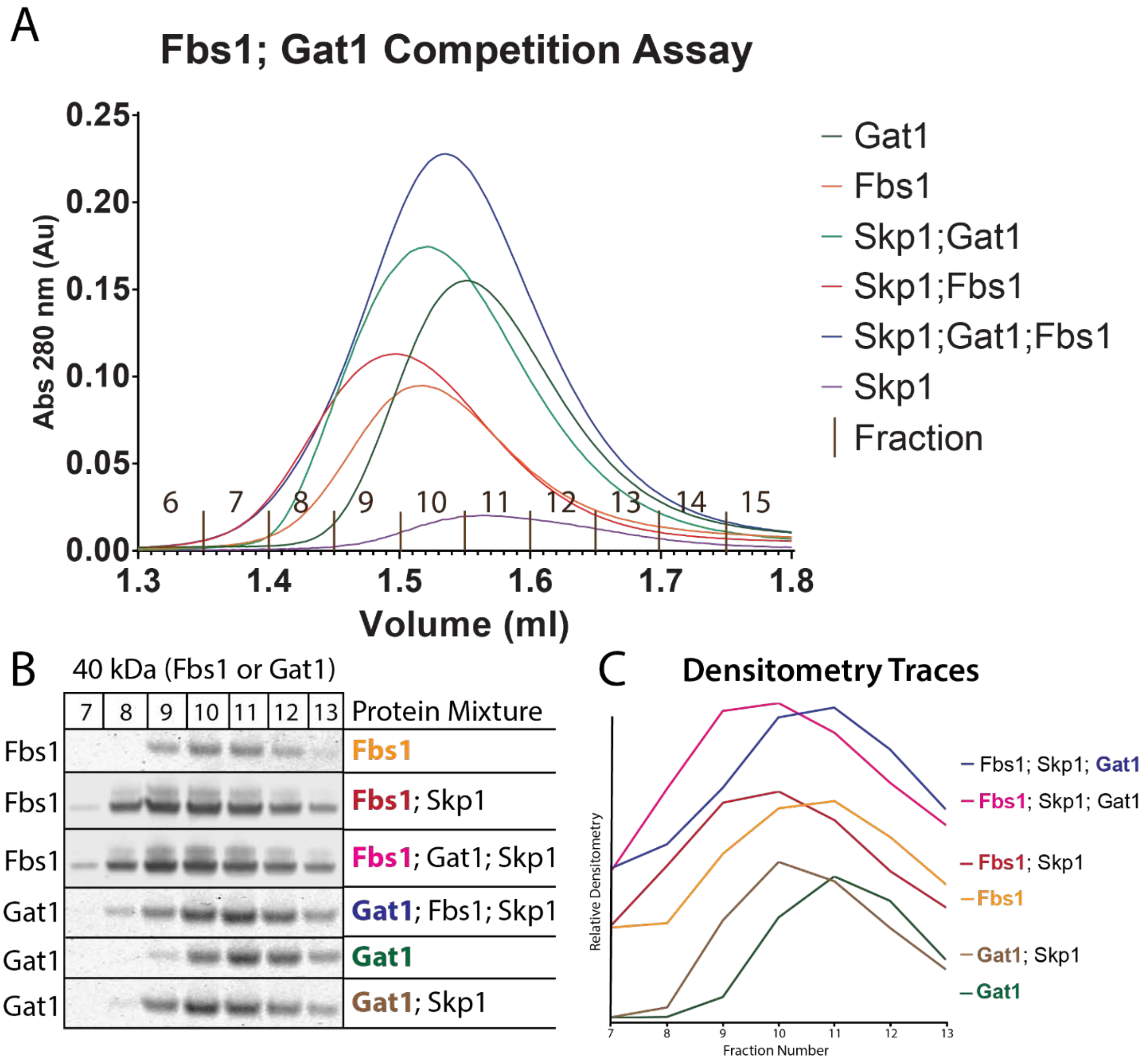

**Figure S10.** Inhibition of PHYa in the presence of varying concentrations of GAT1. (A) PHYa inhibition assays were conducted according to the following framework. Premade mixtures of SKP1 with inhibitors were mixed with PHYa to start the reaction giving a final SKP1 concentration of 100 nM. Reactions were quenched at different times with SDS and subjected to SDS-PAGE and Western blotting and probed with pAb UOK85 which is specific for the reaction product HO-SKP1 relative to the unmodified substrate. Densitometry readings were normalized to a time zero control on the same gel. (B) Varying ratios of HO-SKP1 to SKP1 at 100 nM were analyzed. (C) Densitometry data were used to plot a calibration curve. (D) Percent hydroxylation for reactions quenched at 23 min were plotted to show the effect of GAT1 on SKP1's hydroxylation efficiency. Error bars represent standard deviation of values from 2-3 replicates. (E) Western blots used to calculate hydroxylation efficiency under each condition include effects of chemical inhibitors 500 µM protocatechuic acid (PCA) or 500 µM folate.

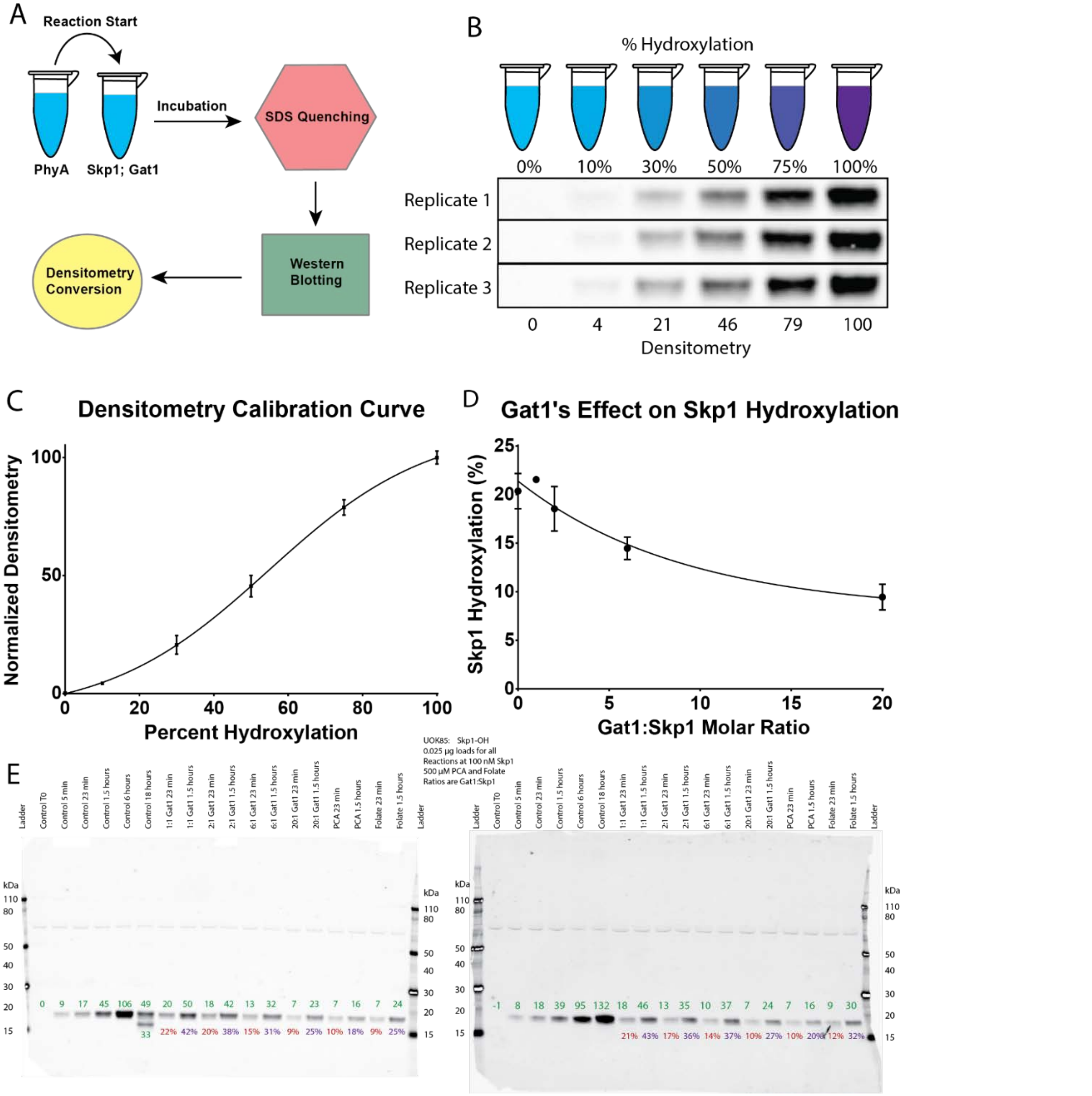

**Figure S11.** Overlay of an AlphaFold2 structure of a 2:1 GAT1:SKP1 complex with a glycan docked PuGAT1 homodimer structure. Overlay of a 2:1 GAT1:SKP1 complex generated by AlphaFold2 with the PuGAT1 crystal structure with a docked tetrasaccharide as reported by (Mandalasi et al, 2020). SKP1's CTR is positioned in such a way as to allow its tetrasaccharide to bind into the active site of GAT1. SKP1 is shown as green with Pro154 of SKP1 shown in red. *Pythium ultimum* GAT1 Subunit1 is shown as blue with its GAT1 Subunit2 shown in orange. The tetrasaccharide docks into the active site of the opposite dimer subunit allowing for the addition of the terminal Gal moiety of the SKP1 pentasaccharide.

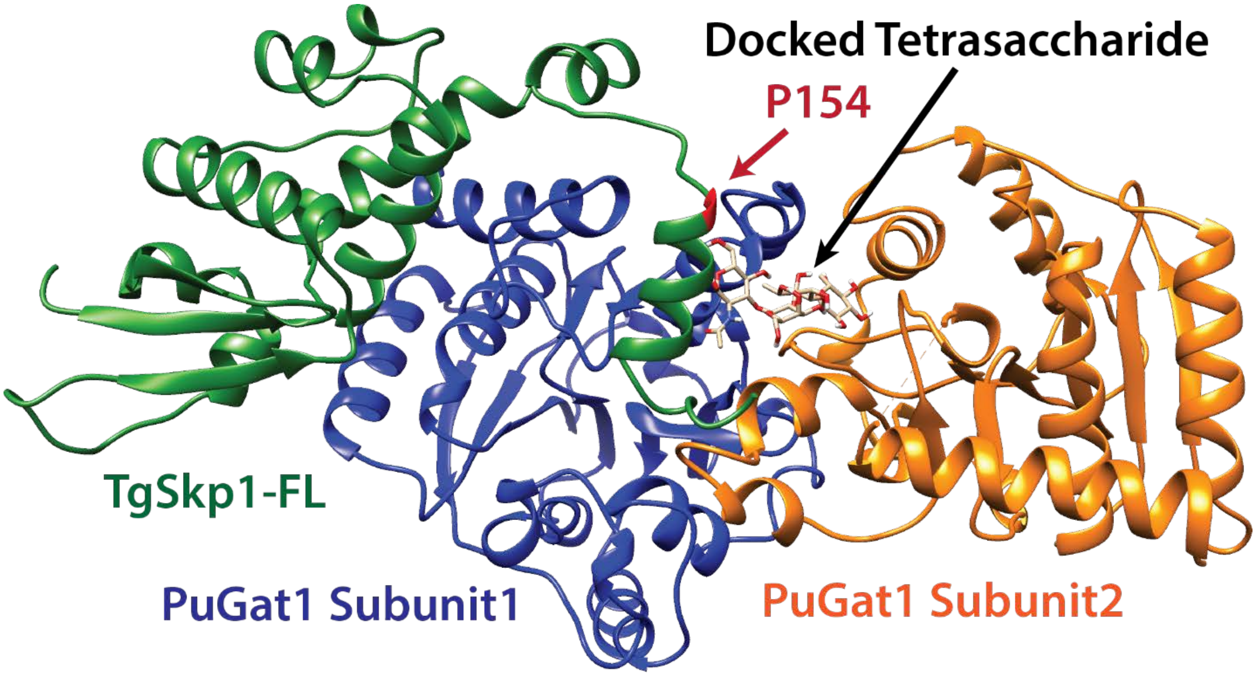

**Figure S12.** Per residue MMGBSA scores for GAT1/SKP1 complexes. Scores for GAT1 are the sum of energetic contributions from each homodimer subunit. SKP1's disordered N-terminus, and the disordered internal loops of both SKP1 and GAT1 are highlighted. Subsites A and B of the SKP1 homodimer are indicated. Coloring of the predicted energetic contributions and standard error of the mean across all 5 simulations are as indicated.

(A) Calculated with a folded SKP1-CTR.

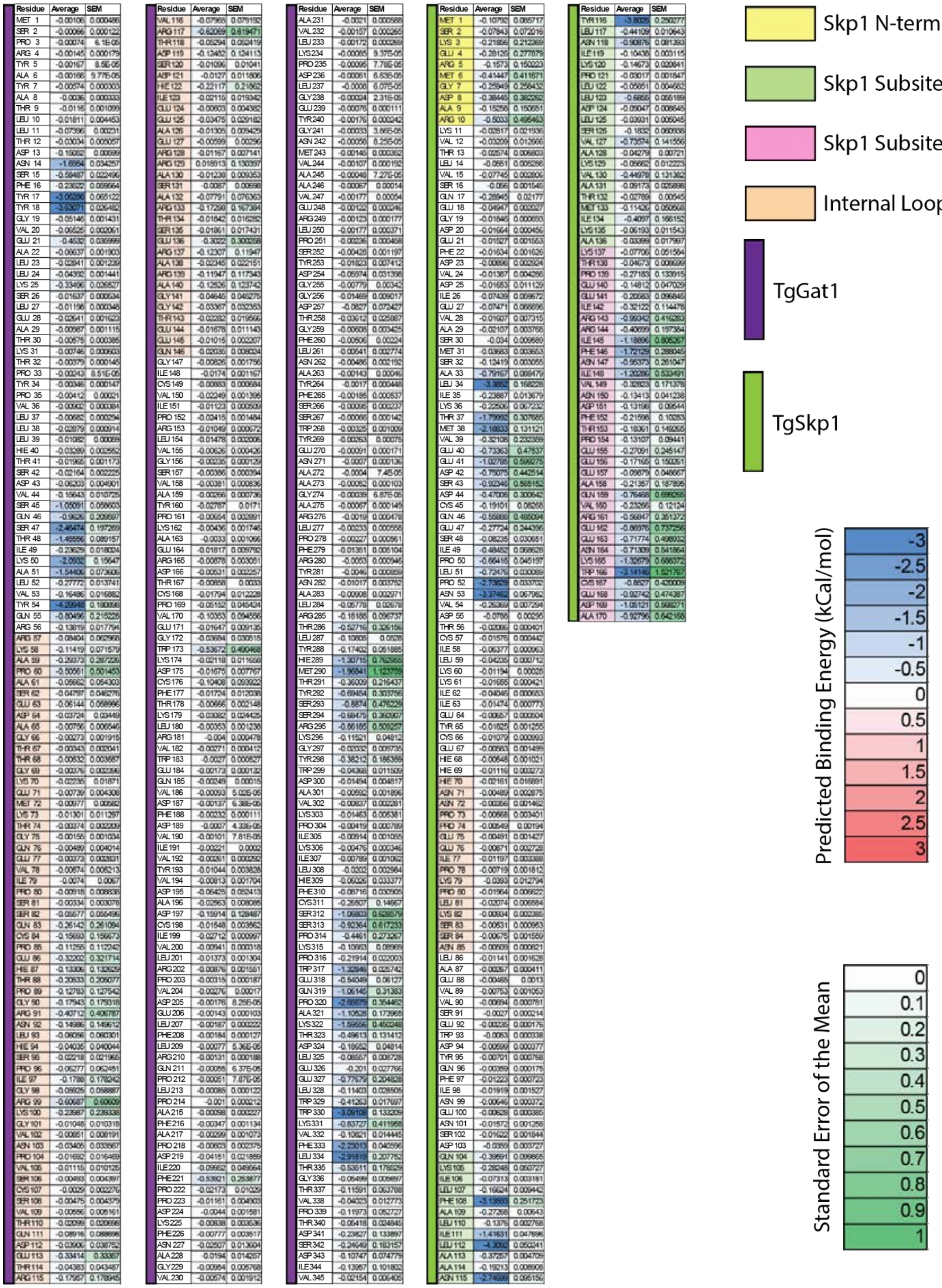

(B) Calculated with a disordered SKP1-CTR.

| Residue | Average | SEM |
|---|---|---|
| MET 1 | -0.06864 | 0.067817 |
| SER 2 | -0.05873 | 0.058029 |
| PRO 3 | -0.01143 | 0.010502 |
| ARG 4 | -0.00591 | 0.003646 |
| TYR 5 | -0.00689 | 0.005054 |
| ALA 6 | -0.00315 | 0.001535 |
| TYR 7 | -0.01378 | 0.00817 |
| ALA 8 | -0.00382 | 0.0004 |
| THR 9 | -0.01035 | 0.000787 |
| LEU 10 | -0.01692 | 0.00626 |
| LEU 11 | -0.05575 | 0.011445 |
| THR 12 | -0.07058 | 0.051303 |
| ASP 13 | -0.1381 | 0.036916 |
| ASN 14 | -1.43001 | 0.372171 |
| SER 15 | -0.72017 | 0.234303 |
| PHE 16 | -1.18623 | 1.038098 |
| TYR 17 | -2.52976 | 0.421957 |
| TYR 18 | -3.28781 | 0.752679 |
| GLY 19 | -0.13538 | 0.088504 |
| VAL 20 | -0.09459 | 0.03411 |
| GLU 21 | -0.51254 | 0.120013 |
| ALA 22 | -0.2136 | 0.13487 |
| LEU 23 | -0.0958 | 0.083988 |
| LEU 24 | -0.06187 | 0.010414 |
| LYS 25 | -0.67063 | 0.281726 |
| SER 26 | -0.11723 | 0.083669 |
| LEU 27 | -0.04312 | 0.018788 |
| GLU 28 | -0.12247 | 0.089798 |
| ALA 29 | -0.20287 | 0.167935 |
| THR 30 | -0.22796 | 0.199205 |
| LYS 31 | -0.39085 | 0.351107 |
| THR 32 | -0.09236 | 0.086079 |
| PRO 33 | -0.2196 | 0.215143 |
| TYR 34 | -0.03872 | 0.034754 |
| PRO 35 | -0.0129 | 0.008573 |
| VAL 36 | -0.01133 | 0.003236 |
| LEU 37 | -0.00929 | 0.00187 |
| LEU 38 | -0.02365 | 0.00282 |
| LEU 39 | -0.02968 | 0.020776 |
| HIE 40 | -0.04164 | 0.015972 |
| THR 41 | -0.02734 | 0.012972 |
| SER 42 | -0.0347 | 0.018979 |
| ASP 43 | -0.04747 | 0.011027 |
| VAL 44 | -0.21193 | 0.103889 |
| SER 45 | -0.68361 | 0.195111 |
| GLN 46 | -1.4113 | 0.696683 |
| SER 47 | -1.8069 | 0.520243 |
| THR 48 | -1.71978 | 0.641524 |
| ILE 49 | -0.59906 | 0.407682 |
| LYS 50 | -1.42091 | 0.411532 |
| ALA 51 | -1.2453 | 0.193318 |
| LEU 52 | -0.88152 | 0.631606 |
| VAL 53 | -0.26582 | 0.111643 |
| TYR 54 | -3.52476 | 0.800665 |
| GLN 55 | -1.66094 | 0.708771 |
| ARG 56 | -0.58585 | 0.352445 |
| ARG 57 | -0.26438 | 0.213282 |
| LYS 58 | -0.18496 | 0.080207 |
| ALA 59 | -0.02603 | 0.013233 |
| PRO 60 | -0.03952 | 0.022572 |
| ALA 61 | -0.18282 | 0.168928 |
| SER 62 | -0.02575 | 0.017567 |
| GLU 63 | -0.01955 | 0.009621 |
| ASP 64 | -0.03407 | 0.030704 |
| ALA 65 | -0.03634 | 0.032488 |
| GLY 66 | 0.017523 | 0.034446 |
| THR 67 | -0.47102 | 0.43475 |
| THR 68 | -0.33778 | 0.316302 |
| GLY 69 | -0.36607 | 0.3467 |
| LYS 70 | -0.16274 | 0.112372 |
| GLU 71 | -0.45324 | 0.410402 |
| MET 72 | -0.29818 | 0.28068 |
| LYS 73 | -0.85756 | 0.847842 |
| THR 74 | -0.22387 | 0.206344 |
| GLY 75 | -0.03616 | 0.022158 |
| GLN 76 | -0.37591 | 0.252987 |
| GLU 77 | -0.45968 | 0.288532 |
| VAL 78 | -0.20176 | 0.188336 |
| ILE 79 | -0.1698 | 0.158105 |
| PRO 80 | -0.03541 | 0.032114 |
| SER 81 | -0.09877 | 0.094911 |
| SER 82 | -0.3315 | 0.329528 |
| GLN 83 | -0.1963 | 0.194342 |
| CYS 84 | -0.00722 | 0.005753 |
| PRO 85 | -0.16656 | 0.164643 |
| GLU 86 | -0.31998 | 0.310608 |
| HIE 87 | -0.15452 | 0.139083 |
| THR 88 | -0.20373 | 0.162133 |
| PRO 89 | -0.71947 | 0.648592 |
| GLY 90 | -0.4324 | 0.366649 |
| ARG 91 | -0.78486 | 0.628613 |
| ASN 92 | -0.41855 | 0.336721 |
| LEU 93 | -0.09024 | 0.057133 |
| HIE 94 | -0.70167 | 0.563158 |
| SER 95 | -0.27316 | 0.246574 |
| PRO 96 | -0.61168 | 0.371098 |
| ILE 97 | -0.44661 | 0.307454 |
| GLY 98 | -0.08092 | 0.04882 |
| ARG 99 | -0.28174 | 0.236262 |
| LYS 100 | -0.07829 | 0.055271 |
| GLY 101 | -0.01522 | 0.008334 |
| VAL 102 | -0.04402 | 0.021902 |
| ASN 103 | -0.01533 | 0.007027 |
| PRO 104 | -0.03152 | 0.022597 |
| VAL 105 | -0.02732 | 0.012974 |
| SER 106 | -0.00902 | 0.004001 |
| CYS 107 | -0.01489 | 0.007937 |
| SER 108 | -0.01367 | 0.009828 |
| VAL 109 | -0.06443 | 0.059596 |
| THR 110 | -0.06565 | 0.061749 |
| GLN 111 | -0.07595 | 0.073521 |
| ASP 112 | -0.1907 | 0.188876 |
| GLU 113 | -0.22237 | 0.22093 |
| THR 114 | -0.29553 | 0.294814 |
| ARG 115 | -0.6136 | 0.611293 |

| Residue | Average | SEM |
|---|---|---|
| VAL 116 | -0.57631 | 0.575344 |
| ARG 117 | -0.09455 | 0.091486 |
| THR 118 | -0.2387 | 0.236064 |
| ASP 119 | -0.05496 | 0.051681 |
| SER 120 | -0.08249 | 0.076826 |
| ASP 121 | -0.01417 | 0.009072 |
| HIE 122 | -0.04108 | 0.021565 |
| ILE 123 | -0.0687 | 0.026436 |
| GLU 124 | -0.02206 | 0.009609 |
| GLU 125 | -0.04324 | 0.032486 |
| ALA 126 | -0.06953 | 0.040313 |
| GLU 127 | -0.20938 | 0.190849 |
| ARG 128 | -0.03941 | 0.025947 |
| ARG 129 | -0.19067 | 0.150711 |
| ALA 130 | -0.25153 | 0.205549 |
| SER 131 | -0.20001 | 0.190078 |
| ALA 132 | -0.05571 | 0.043007 |
| ARG 133 | -0.40697 | 0.260043 |
| THR 134 | -0.57526 | 0.553124 |
| SER 135 | -0.23921 | 0.233606 |
| GLU 136 | -0.31244 | 0.296128 |
| ARG 137 | -1.00287 | 0.985274 |
| ALA 138 | -0.68332 | 0.677889 |
| ARG 139 | -1.18386 | 1.17687 |
| ALA 140 | -0.7456 | 0.73797 |
| GLY 141 | -0.43444 | 0.429638 |
| GLY 142 | -0.304 | 0.296999 |
| THR 143 | -0.60511 | 0.588277 |
| GLU 144 | -0.73025 | 0.704821 |
| GLU 145 | -0.77557 | 0.765644 |
| GLN 146 | -0.29729 | 0.273393 |
| GLY 147 | -0.06121 | 0.060172 |
| ILE 148 | -0.09047 | 0.045609 |
| CYS 149 | -0.05718 | 0.030436 |
| VAL 150 | -0.03039 | 0.00515 |
| ILE 151 | -0.01976 | 0.006007 |
| PRO 152 | -0.02134 | 0.002288 |
| ARG 153 | -0.03041 | 0.020662 |
| LEU 154 | -0.01249 | 0.001956 |
| VAL 155 | -0.00655 | 0.001691 |
| GLY 156 | -0.00263 | 0.000952 |
| SER 157 | -0.00275 | 0.000571 |
| VAL 158 | -0.00224 | 0.000631 |
| ALA 159 | -0.0013 | 0.000415 |
| TYR 160 | -0.00361 | 0.001472 |
| PRO 161 | -0.00355 | 0.002616 |
| LYS 162 | -0.01361 | 0.011832 |
| ALA 163 | -0.00989 | 0.008436 |
| GLU 164 | -0.06384 | 0.062233 |
| ARG 165 | -0.08706 | 0.084869 |
| ASP 166 | -0.04038 | 0.036409 |
| THR 167 | -0.085 | 0.078812 |
| CYS 168 | -0.09175 | 0.08797 |
| PRO 169 | -0.06763 | 0.059696 |
| VAL 170 | -0.43117 | 0.42819 |
| GLU 171 | -0.48955 | 0.479244 |
| GLY 172 | -0.07787 | 0.072896 |
| TRP 173 | -0.19545 | 0.179854 |
| LYS 174 | -0.04971 | 0.042226 |
| ASP 175 | -0.0535 | 0.047687 |
| CYS 176 | -0.01688 | 0.012622 |
| PHE 177 | -0.00801 | 0.005669 |
| THR 178 | -0.00795 | 0.004759 |
| LYS 179 | -0.05047 | 0.047017 |
| LEU 180 | -0.00392 | 0.002304 |
| ARG 181 | -0.00443 | 0.001529 |
| VAL 182 | -0.00371 | 0.001642 |
| TRP 183 | -0.00668 | 0.004793 |
| GLU 184 | -0.00178 | 0.000296 |
| GLN 185 | -0.00259 | 0.000212 |
| VAL 186 | -0.0015 | 0.000476 |
| ASP 187 | -0.00192 | 0.000448 |
| PHE 188 | -0.00404 | 0.00159 |
| ASP 189 | -0.00153 | 0.000637 |
| VAL 190 | -0.00185 | 0.000836 |
| ILE 191 | -0.00287 | 0.000715 |
| VAL 192 | -0.00394 | 0.001378 |
| TYR 193 | -0.00567 | 0.000154 |
| VAL 194 | -0.00661 | 0.000518 |
| ASP 195 | -0.00796 | 0.000935 |
| ALA 196 | -0.01233 | 0.002852 |
| ASP 197 | -0.01387 | 0.001865 |
| CYS 198 | -0.00996 | 0.0008 |
| ILE 199 | -0.026 | 0.005434 |
| VAL 200 | -0.02424 | 0.01256 |
| LEU 201 | -0.07682 | 0.067811 |
| ARG 202 | -0.26943 | 0.216072 |
| PRO 203 | -0.38092 | 0.37159 |
| VAL 204 | -0.03131 | 0.024719 |
| ASP 205 | 0.098602 | 0.101856 |
| GLU 206 | -0.04184 | 0.038718 |
| LEU 207 | -0.00636 | 0.004195 |
| PHE 208 | -0.00849 | 0.006616 |
| LEU 209 | -0.00529 | 0.003872 |
| ARG 210 | -0.00333 | 0.001536 |
| GLN 211 | -0.00118 | 0.000387 |
| PRO 212 | -0.00073 | 0.000166 |
| LEU 213 | -0.00106 | 0.000288 |
| PRO 214 | -0.0009 | 0.000113 |
| ALA 215 | -0.0008 | 0.000168 |
| PHE 216 | -0.00187 | 0.000237 |
| ALA 217 | -0.00124 | 0.00016 |
| PRO 218 | -0.00189 | 0.000381 |
| ASP 219 | -0.00215 | 0.000443 |
| ILE 220 | -0.00371 | 0.001074 |
| PHE 221 | -0.00431 | 0.001166 |
| PRO 222 | -0.00335 | 0.001246 |
| PRO 223 | -0.0048 | 0.002647 |
| ASP 224 | -0.00256 | 0.000956 |
| LYS 225 | -0.00244 | 0.000759 |
| PHE 226 | -0.01449 | 0.012796 |
| ASN 227 | -0.00362 | 0.001515 |
| ALA 228 | -0.00479 | 0.003239 |
| GLY 229 | -0.00426 | 0.002538 |
| VAL 230 | -0.00735 | 0.004082 |

| Residue | Average | SEM |
|---|---|---|
| ALA 231 | -0.00344 | 0.001023 |
| VAL 232 | -0.00293 | 0.001165 |
| LEU 233 | -0.00205 | 0.000509 |
| LYS 234 | -0.00142 | 0.000429 |
| PRO 235 | -0.0011 | 0.000185 |
| ASP 236 | -0.00083 | 7.72E-05 |
| LEU 237 | -0.00073 | 3.21E-05 |
| GLY 238 | -0.00023 | 1.8E-05 |
| GLU 239 | -0.00058 | 3.4E-05 |
| TYR 240 | -0.00122 | 5.55E-05 |
| GLY 241 | -0.00028 | 3.68E-05 |
| ASN 242 | -0.00065 | 0.000202 |
| MET 243 | -0.00126 | 0.000433 |
| VAL 244 | -0.00118 | 0.000522 |
| ALA 245 | -0.00051 | 0.000163 |
| ALA 246 | -0.00062 | 0.000197 |
| VAL 247 | -0.00145 | 0.000667 |
| GLU 248 | -0.00119 | 0.00051 |
| ARG 249 | -0.00189 | 0.00087 |
| LEU 250 | -0.00172 | 0.00068 |
| PRO 251 | -0.00324 | 0.001741 |
| SER 252 | -0.003 | 0.001566 |
| TYR 253 | -0.0094 | 0.007112 |
| ASP 254 | -0.01428 | 0.009718 |
| GLY 255 | -0.00648 | 0.005382 |
| GLY 256 | -0.00788 | 0.006935 |
| ASP 257 | -0.02932 | 0.027411 |
| THR 258 | -0.07503 | 0.073465 |
| GLY 259 | -0.00593 | 0.005223 |
| PHE 260 | -0.00501 | 0.003667 |
| LEU 261 | -0.00502 | 0.003751 |
| ASN 262 | -0.00323 | 0.002097 |
| ALA 263 | -0.00138 | 0.000706 |
| TYR 264 | -0.00171 | 0.000749 |
| PHE 265 | -0.00197 | 0.000977 |
| SER 266 | -0.00095 | 0.000325 |
| SER 267 | -0.00089 | 0.000298 |
| TRP 268 | -0.00192 | 0.000411 |
| TYR 269 | -0.00372 | 0.00228 |
| GLU 270 | -0.00186 | 0.000846 |
| ASN 271 | -0.00071 | 0.000163 |
| ALA 272 | -0.0005 | 0.000104 |
| ALA 273 | -0.00059 | 0.000142 |
| GLY 274 | -0.0006 | 0.000158 |
| ALA 275 | -0.00071 | 0.000146 |
| ARG 276 | -0.00192 | 0.000451 |
| LEU 277 | -0.00266 | 0.00068 |
| PRO 278 | -0.00581 | 0.003003 |
| PHE 279 | -0.00529 | 0.001287 |
| ARG 280 | -0.01618 | 0.005834 |
| TYR 281 | -0.01942 | 0.009782 |
| ASN 282 | -0.00783 | 0.004167 |
| ALA 283 | -0.00566 | 0.001838 |
| LEU 284 | -0.02474 | 0.019728 |
| ARG 285 | -0.02372 | 0.004796 |
| THR 286 | -0.01906 | 0.013602 |
| LEU 287 | -0.01868 | 0.013042 |
| TYR 288 | -0.04192 | 0.014651 |
| HIE 289 | -0.11201 | 0.076302 |
| MET 290 | -0.0388 | 0.030804 |
| THR 291 | -0.01361 | 0.006844 |
| TYR 292 | -0.12229 | 0.06929 |
| SER 293 | -0.07024 | 0.038152 |
| SER 294 | -0.0637 | 0.040961 |
| ARG 295 | -0.08284 | 0.053208 |
| LYS 296 | -0.27175 | 0.163727 |
| GLY 297 | -0.14973 | 0.124456 |
| TYR 298 | -1.1385 | 1.121497 |
| TRP 299 | -0.06626 | 0.036283 |
| ASP 300 | 0.015506 | 0.061028 |
| ALA 301 | -0.1074 | 0.095765 |
| VAL 302 | -0.39584 | 0.387199 |
| LYS 303 | -0.18241 | 0.122633 |
| PRO 304 | -0.2458 | 0.201366 |
| ILE 305 | -0.06714 | 0.046673 |
| LYS 306 | -0.0798 | 0.046173 |
| ILE 307 | -0.01321 | 0.004044 |
| LEU 308 | -0.01559 | 0.002216 |
| HIE 309 | -0.01362 | 0.004793 |
| PHE 310 | -0.02641 | 0.005344 |
| CYS 311 | -0.00813 | 0.002 |
| SER 312 | -0.01199 | 0.004555 |
| SER 313 | -0.02086 | 0.006317 |
| PRO 314 | -0.07441 | 0.025062 |
| LYS 315 | -0.10623 | 0.040135 |
| PRO 316 | -0.39084 | 0.225236 |
| TRP 317 | -1.46787 | 0.450037 |
| GLU 318 | -0.69254 | 0.269163 |
| GLN 319 | -0.63395 | 0.247477 |
| PRO 320 | -1.69424 | 0.739062 |
| ALA 321 | -0.84368 | 0.316871 |
| LYS 322 | -0.57165 | 0.224045 |
| THR 323 | -0.19851 | 0.063307 |
| ASP 324 | -0.46312 | 0.368854 |
| LEU 325 | -0.23524 | 0.181329 |
| GLU 326 | -0.16312 | 0.050676 |
| GLU 327 | -0.9923 | 0.35964 |
| LEU 328 | -0.95099 | 0.828236 |
| TRP 329 | -0.38234 | 0.079921 |
| TRP 330 | -2.38934 | 0.904458 |
| LYS 331 | -1.38258 | 0.527736 |
| VAL 332 | -0.51512 | 0.366042 |
| PHE 333 | -1.86617 | 0.398451 |
| LEU 334 | -3.39666 | 0.320711 |
| THR 335 | -1.58964 | 0.602786 |
| GLY 336 | -0.20384 | 0.086355 |
| THR 337 | -0.74921 | 0.437236 |
| VAL 338 | -0.68921 | 0.35672 |
| PRO 339 | -0.67516 | 0.264112 |
| THR 340 | -0.78387 | 0.434286 |
| ASP 341 | -0.92384 | 0.518908 |
| SER 342 | -0.64583 | 0.371553 |
| ASP 343 | -0.45942 | 0.299355 |
| ILE 344 | -0.52569 | 0.445969 |
| VAL 345 | -0.2649 | 0.114653 |

| Residue | Average | SEM |
|---|---|---|
| MET 1 | -2.05332 | 1.289112 |
| SER 2 | -1.04404 | 0.773047 |
| LYS 3 | -0.90106 | 0.571718 |
| GLU 4 | -0.28177 | 0.225568 |
| ARG 5 | -0.15229 | 0.10506 |
| MET 6 | -0.8757 | 0.830205 |
| GLY 7 | -0.27574 | 0.250968 |
| ASP 8 | -0.06831 | 0.044141 |
| ALA 9 | -0.03364 | 0.0275 |
| ARG 10 | -0.03315 | 0.017077 |
| LYS 11 | -0.07317 | 0.065274 |
| VAL 12 | -0.02396 | 0.006264 |
| THR 13 | -0.02689 | 0.008781 |
| LEU 14 | -0.05401 | 0.014075 |
| VAL 15 | -0.06682 | 0.014108 |
| SER 16 | -0.07643 | 0.021606 |
| GLN 17 | -0.4083 | 0.210659 |
| GLU 18 | -0.1422 | 0.061276 |
| GLY 19 | -0.04577 | 0.026908 |
| ASP 20 | -0.06522 | 0.045131 |
| GLU 21 | -0.0254 | 0.012002 |
| PHE 22 | -0.0428 | 0.02682 |
| ASP 23 | -0.00969 | 0.003598 |
| VAL 24 | -0.01072 | 0.002334 |
| ASP 25 | -0.00877 | 0.003049 |
| ILE 26 | -0.02202 | 0.009586 |
| GLU 27 | 0.007582 | 0.013617 |
| VAL 28 | -0.01738 | 0.009134 |
| ALA 29 | -0.02439 | 0.010468 |
| SER 30 | -0.12146 | 0.099149 |
| MET 31 | -0.23962 | 0.211973 |
| SER 32 | -0.09515 | 0.022963 |
| ALA 33 | -0.56177 | 0.190272 |
| LEU 34 | -2.64358 | 0.667453 |
| ILE 35 | -0.67818 | 0.517178 |
| LYS 36 | -0.15283 | 0.046739 |
| THR 37 | -1.18529 | 0.313198 |
| MET 38 | -1.9048 | 0.586436 |
| VAL 39 | -0.32108 | 0.196956 |
| GLU 40 | -0.29018 | 0.188209 |
| GLU 41 | -0.67971 | 0.332086 |
| ASP 42 | -0.88037 | 0.566937 |
| SER 43 | -0.44085 | 0.306846 |
| ASP 44 | -0.3145 | 0.19328 |
| CYS 45 | -0.42983 | 0.266982 |
| GLN 46 | -0.7069 | 0.4339 |
| GLU 47 | -0.27883 | 0.170342 |
| SER 48 | -0.28243 | 0.185665 |
| ILE 49 | -0.47188 | 0.147371 |
| PRO 50 | -0.89417 | 0.282404 |
| LEU 51 | -1.28476 | 0.771919 |
| PRO 52 | -2.01803 | 0.593283 |
| ASN 53 | -2.19167 | 0.654619 |
| VAL 54 | -1.11142 | 0.704133 |
| ASP 55 | -0.36899 | 0.248505 |
| THR 56 | -0.3016 | 0.269863 |
| CYS 57 | -0.0867 | 0.066147 |
| ILE 58 | -0.15466 | 0.057623 |
| LEU 59 | -0.10374 | 0.061495 |
| LYS 60 | -0.02418 | 0.010774 |
| LYS 61 | -0.02288 | 0.003299 |
| ILE 62 | -0.04815 | 0.007487 |
| ILE 63 | -0.03647 | 0.023769 |
| GLU 64 | -0.01436 | 0.007816 |
| TYR 65 | -0.02066 | 0.003726 |
| CYS 66 | -0.02128 | 0.012659 |
| GLU 67 | -0.10967 | 0.105309 |
| HIE 68 | -0.01225 | 0.007775 |
| HIE 69 | -0.01779 | 0.010698 |
| HIE 70 | -0.08239 | 0.076475 |
| ASN 71 | -0.01322 | 0.010633 |
| ASN 72 | -0.0104 | 0.006584 |
| PRO 73 | -0.01072 | 0.006198 |
| PRO 74 | -0.04298 | 0.027808 |
| GLU 75 | -0.04074 | 0.026189 |
| GLU 76 | -0.05694 | 0.030738 |
| ILE 77 | -0.28893 | 0.273604 |
| PRO 78 | -0.30067 | 0.266654 |
| LYS 79 | -0.64288 | 0.582015 |
| PRO 80 | -0.34637 | 0.314625 |
| LEU 81 | -0.07906 | 0.050266 |
| LYS 82 | -0.13187 | 0.114546 |
| SER 83 | -0.0226 | 0.012501 |
| SER 84 | -0.01754 | 0.005132 |
| ASN 85 | -0.01734 | 0.006924 |
| LEU 86 | -0.03143 | 0.012686 |
| ALA 87 | -0.00495 | 0.001319 |
| GLU 88 | -0.00669 | 0.001421 |
| VAL 89 | -0.00931 | 0.001426 |
| VAL 90 | -0.0076 | 0.001137 |
| SER 91 | -0.00317 | 0.00027 |
| GLU 92 | -0.00361 | 0.000689 |
| TRP 93 | -0.00612 | 0.000603 |
| ASP 94 | -0.0073 | 0.000599 |
| TYR 95 | -0.01243 | 0.002205 |
| GLN 96 | -0.00628 | 0.001314 |
| PHE 97 | -0.01745 | 0.002757 |
| ILE 98 | -0.03132 | 0.00443 |
| ASN 99 | -0.01424 | 0.004299 |
| GLU 100 | -0.01221 | 0.003403 |
| ASN 101 | -0.02901 | 0.00625 |
| SER 102 | -0.06756 | 0.033044 |
| ASP 103 | -0.12087 | 0.058087 |
| GLN 104 | -1.39746 | 0.775209 |
| LYS 105 | -1.26843 | 0.582263 |
| ILE 106 | -0.56543 | 0.407562 |
| LEU 107 | -0.26789 | 0.049371 |
| PHE 108 | -3.28839 | 0.597252 |
| ALA 109 | -1.06902 | 0.497593 |
| LEU 110 | -0.33306 | 0.158 |
| ILE 111 | -1.23102 | 0.267017 |
| LEU 112 | -4.37578 | 0.140265 |
| ALA 113 | -0.59021 | 0.186238 |
| ALA 114 | -0.1861 | 0.024619 |
| ASN 115 | -2.18467 | 0.437791 |

| Residue | Average | SEM |
|---|---|---|
| TYR 116 | -3.78017 | 0.984188 |
| LEU 117 | -0.63624 | 0.295177 |
| ASN 118 | -0.74998 | 0.19957 |
| ILE 119 | -0.57704 | 0.492276 |
| LYS 120 | -0.5399 | 0.438967 |
| PRO 121 | -0.27906 | 0.252705 |
| LEU 122 | -0.10742 | 0.04644 |
| LEU 123 | -0.59039 | 0.128547 |
| ASP 124 | 0.008341 | 0.100371 |
| LEU 125 | -0.08307 | 0.034925 |
| SER 126 | -0.2833 | 0.113201 |
| VAL 127 | -0.86827 | 0.205373 |
| ALA 128 | -0.20249 | 0.145088 |
| LYS 129 | -0.11913 | 0.021981 |
| VAL 130 | -0.86603 | 0.263926 |
| ALA 131 | -0.37395 | 0.227601 |
| THR 132 | -0.20973 | 0.122559 |
| MET 133 | -1.21083 | 0.55667 |
| ILE 134 | -2.14599 | 0.571011 |
| LYS 135 | -0.90839 | 0.279694 |
| ALA 136 | -0.69097 | 0.30768 |
| LYS 137 | -1.11354 | 0.488229 |
| THR 138 | -0.80814 | 0.578144 |
| PRO 139 | -0.25537 | 0.136244 |
| GLU 140 | -0.7986 | 0.67488 |
| GLU 141 | -0.62023 | 0.508829 |
| ILE 142 | -0.62117 | 0.396378 |
| ARG 143 | -1.42584 | 0.820587 |
| ARG 144 | -0.7497 | 0.390007 |
| ILE 145 | -0.67748 | 0.547044 |
| PHE 146 | -1.06579 | 0.8611 |
| ASN 147 | -0.25631 | 0.204679 |
| ILE 148 | -0.64204 | 0.408493 |
| VAL 149 | -0.42219 | 0.240277 |
| ASN 150 | -0.21967 | 0.162406 |
| ASP 151 | -0.05361 | 0.035199 |
| PHE 152 | -0.16126 | 0.130667 |
| THR 153 | -0.11842 | 0.103168 |
| PRO 154 | -0.11182 | 0.080063 |
| GLU 155 | -0.44839 | 0.267236 |
| GLU 156 | -0.17325 | 0.103753 |
| GLU 157 | -0.46614 | 0.362042 |
| ALA 158 | -0.24213 | 0.117802 |
| GLN 159 | -0.77853 | 0.397164 |
| VAL 160 | -0.84605 | 0.358523 |
| ARG 161 | -1.2895 | 0.58066 |
| GLU 162 | -0.27057 | 0.177241 |
| GLU 163 | -1.22016 | 0.49551 |
| ASN 164 | -0.91622 | 0.457806 |
| LYS 165 | -0.92024 | 0.634109 |
| TRP 166 | -2.89177 | 1.603655 |
| CYS 167 | -1.08289 | 0.410843 |
| GLU 168 | -0.97775 | 0.613061 |
| ASP 169 | -0.7047 | 0.296386 |
| ALA 170 | -0.48901 | 0.248246 |

Skp1 N-terminus

Skp1 Subsite-A

Skp1 Subsite-B

Internal Loop

TgGat1

TgSkp1

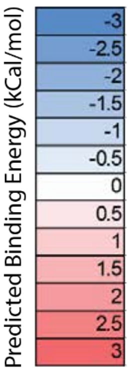


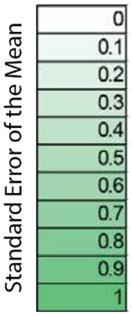

**Figure S13.** Predicted changes in binding energy of GAT1 to SKP1. Predicted complex ΔΔG values from Rosetta for each GAT1 aa substitution in the predicted interface for each cluster are indicated. The weighted average of the lowest 3 scoring clusters was used as the final predicted ΔΔG for each point mutation (bottom panel).

(A) Calculated with a folded SKP1 CTR. Supports Fig. 6C.

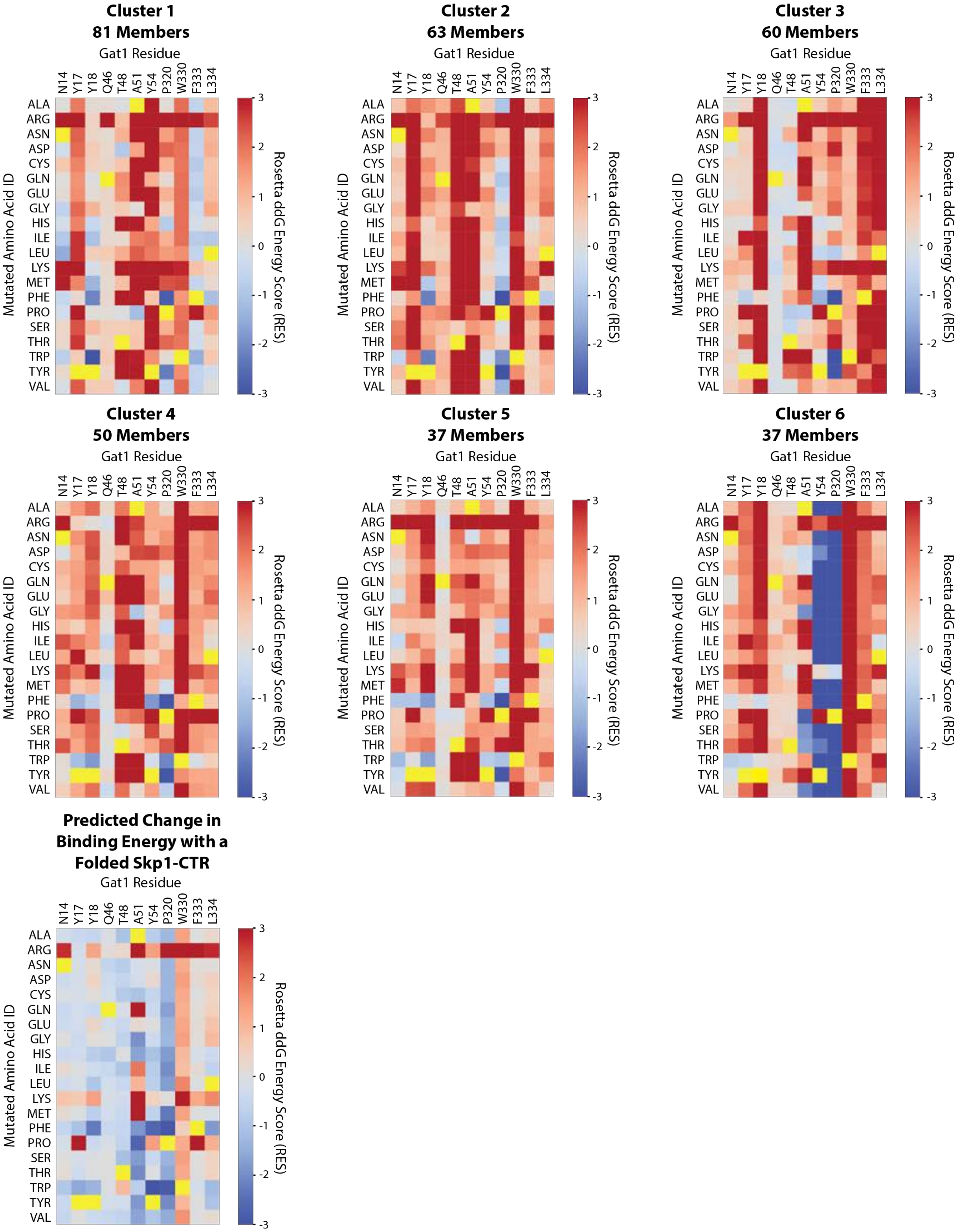

(B) Calculated with a disordered SKP1 CTR. Supports Fig. 6D.

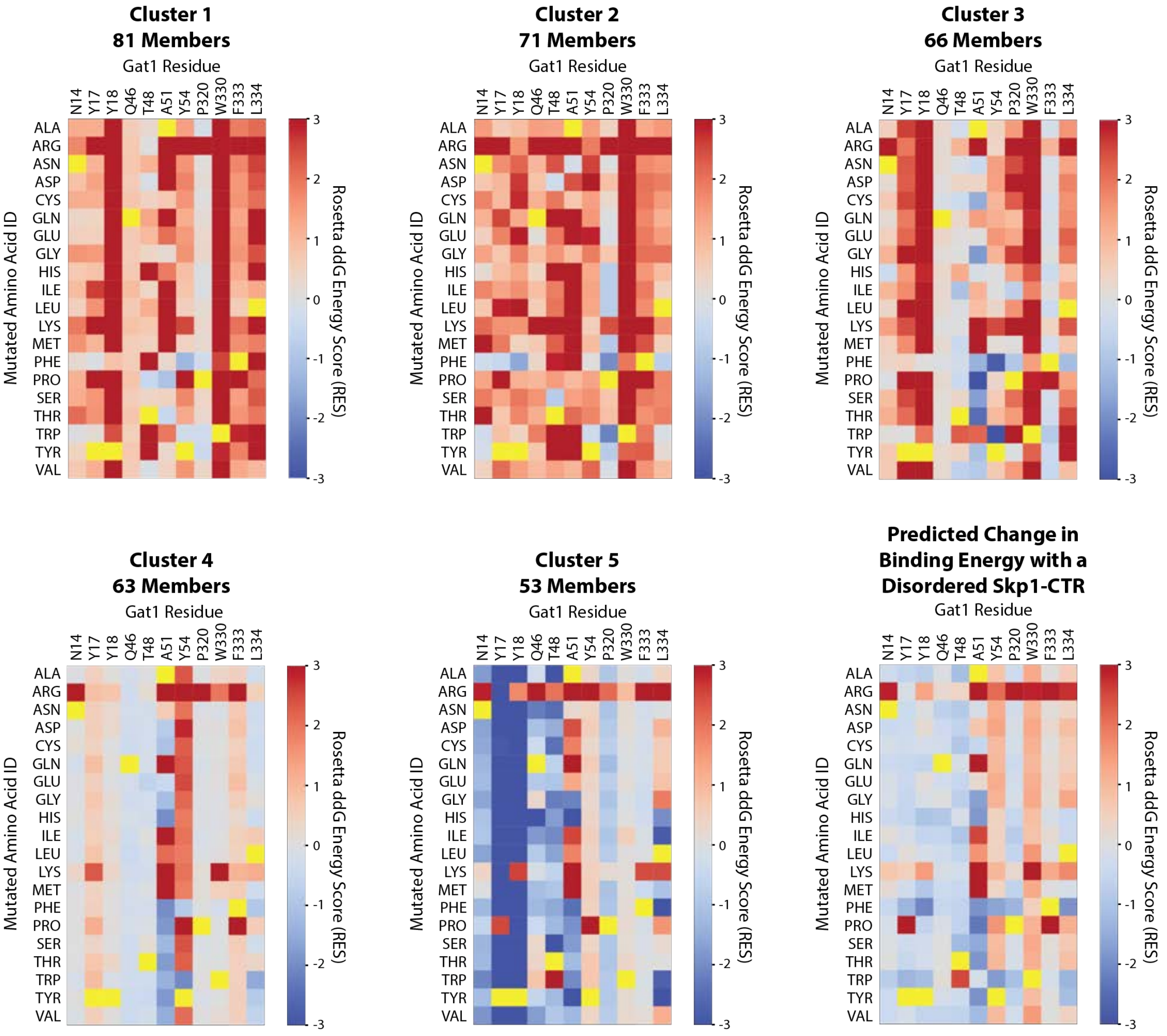

**Figure S14.** Predicted change in stability of the GAT1 homodimer for each cluster. Predicted GAT1 ΔΔG values for each amino acid substitution in each cluster are indicated. The weighted average of the lowest 3 scoring clusters was used as the final predicted ΔΔG for each point mutation. Supports Fig. 6E.

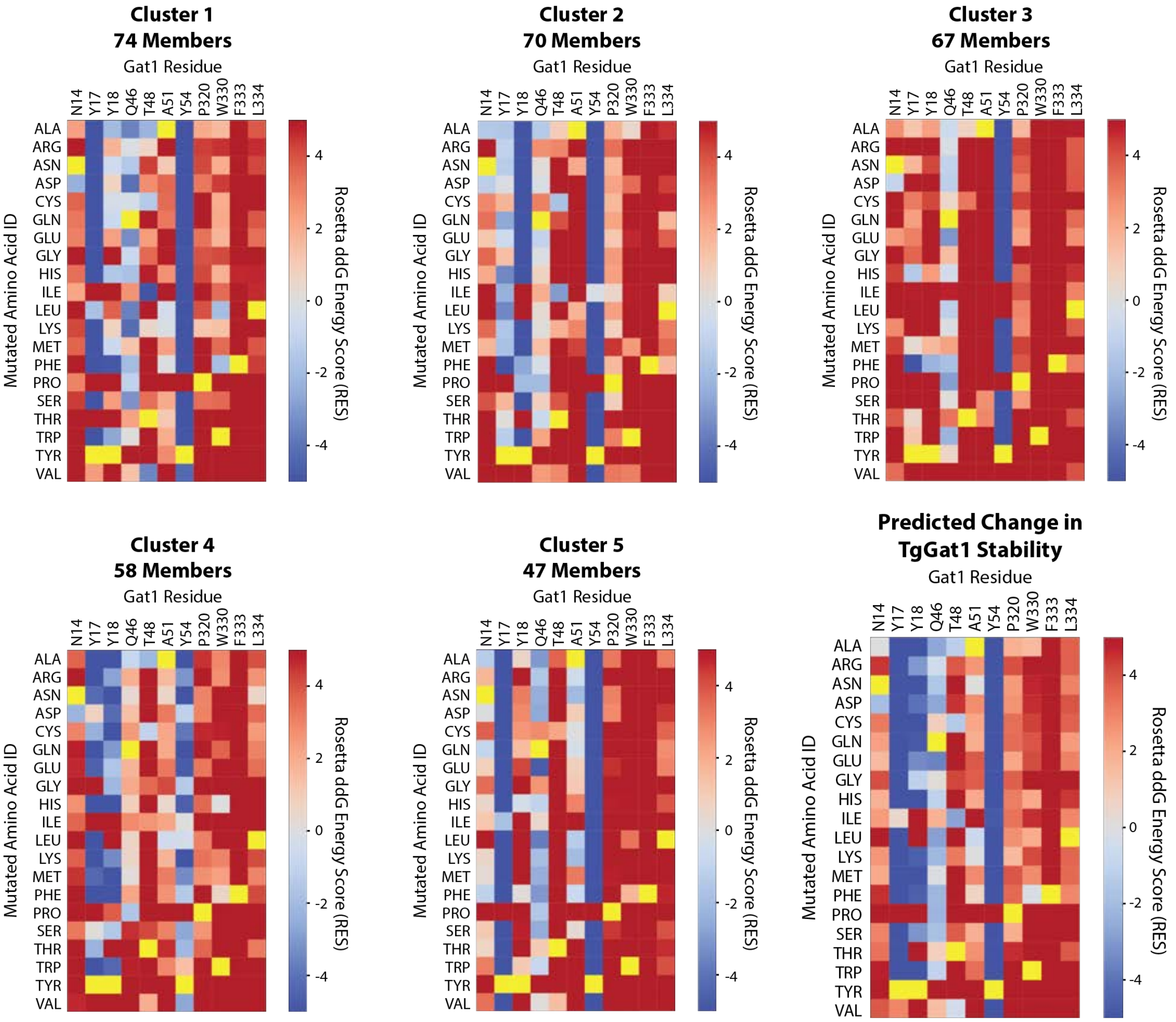

**Figure S15.** pLDDT scores for AlphaFold3 models of mutant GAT1/SKP1 complexes. Average per residue pLDDT scores, as depicted by coloring according to the key at the bottom, were averaged across all 5 structures generated by AlphaFold3 and plotted onto the top scoring model (upper left panels).
(A) Calculated for the 2:1 GAT1(Y18K):SKP1 complex.

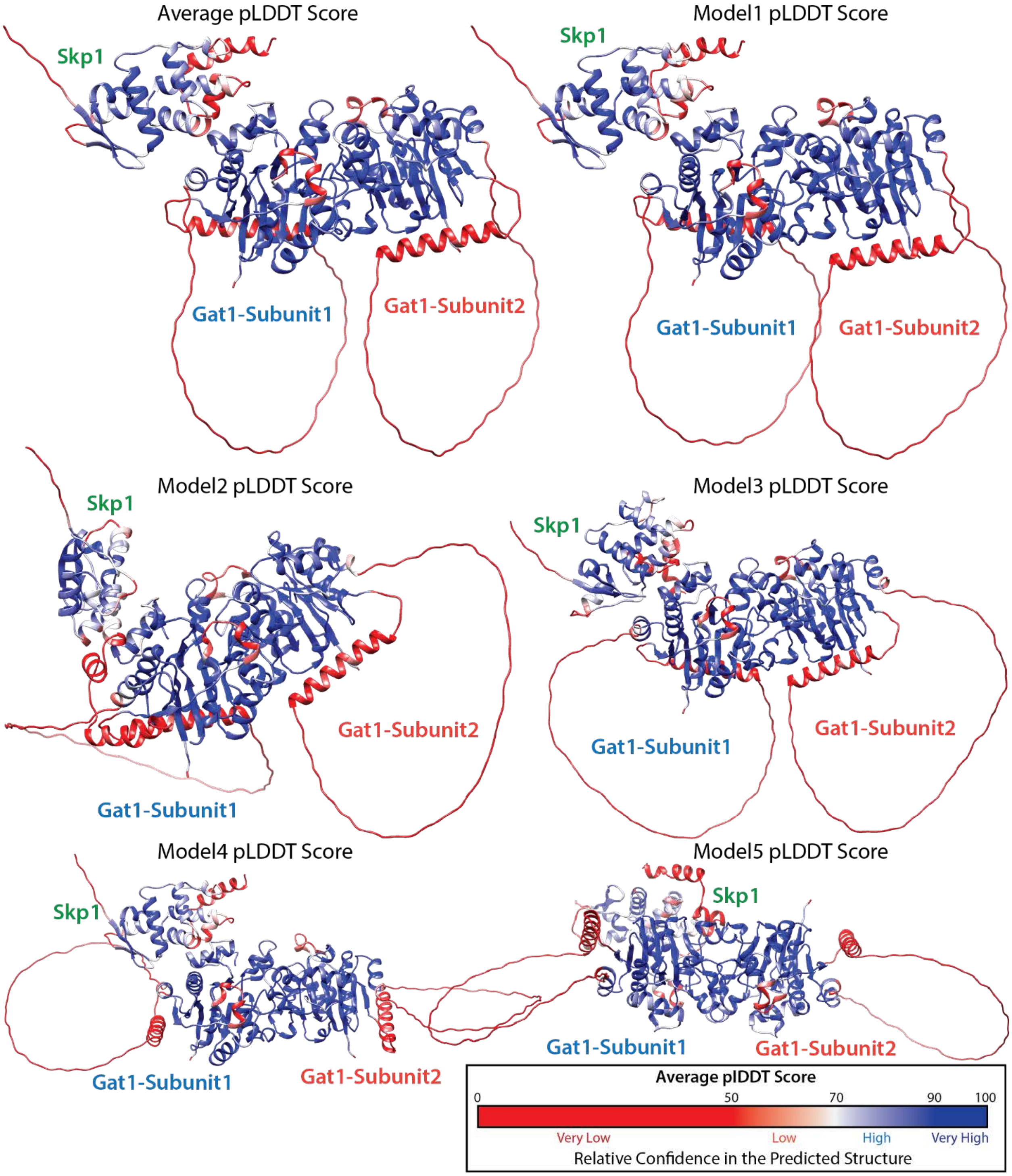

(B) Calculated for the 2:1 GAT1(A51K):SKP1 complex.

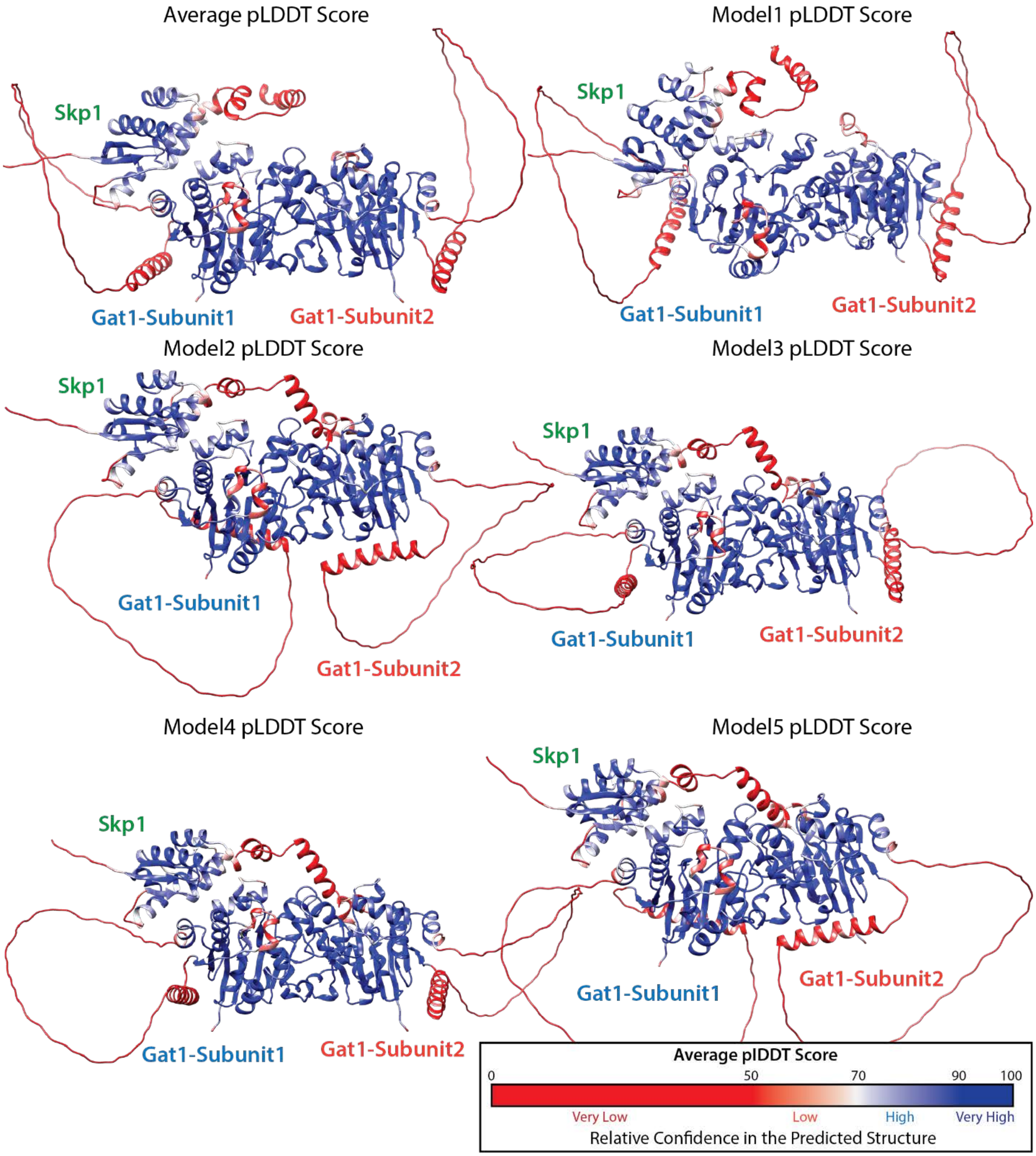

(C) Calculated for the 2:1 GAT1(Y18K/A51K):SKP1 complex.

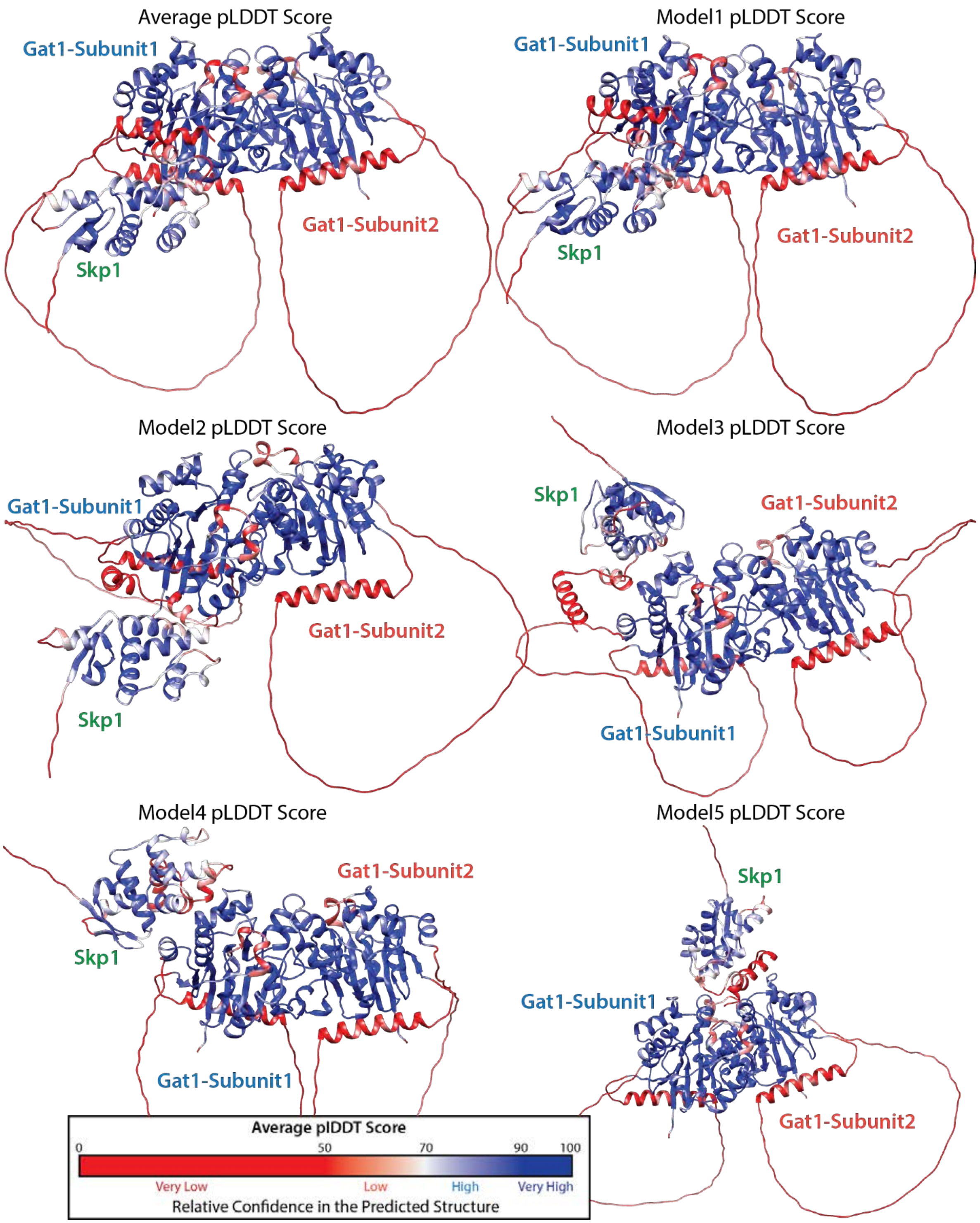

**Figure S16.** A sample of distances between oppositely charged residues between SKP1 and GAT1. Distances between oppositely charged residues with pairwise interaction energies ≤ -1 kcal/mol are shown. A total of 146 such interactions were observed across all 10 simulations. Productively contributing distances between charged functional groups (<8 Å) are highlighted. Trajectories shown are selected mostly from Simulation 1 starting with the folded SKP1-CTR or unfolded SKP1-CTR.

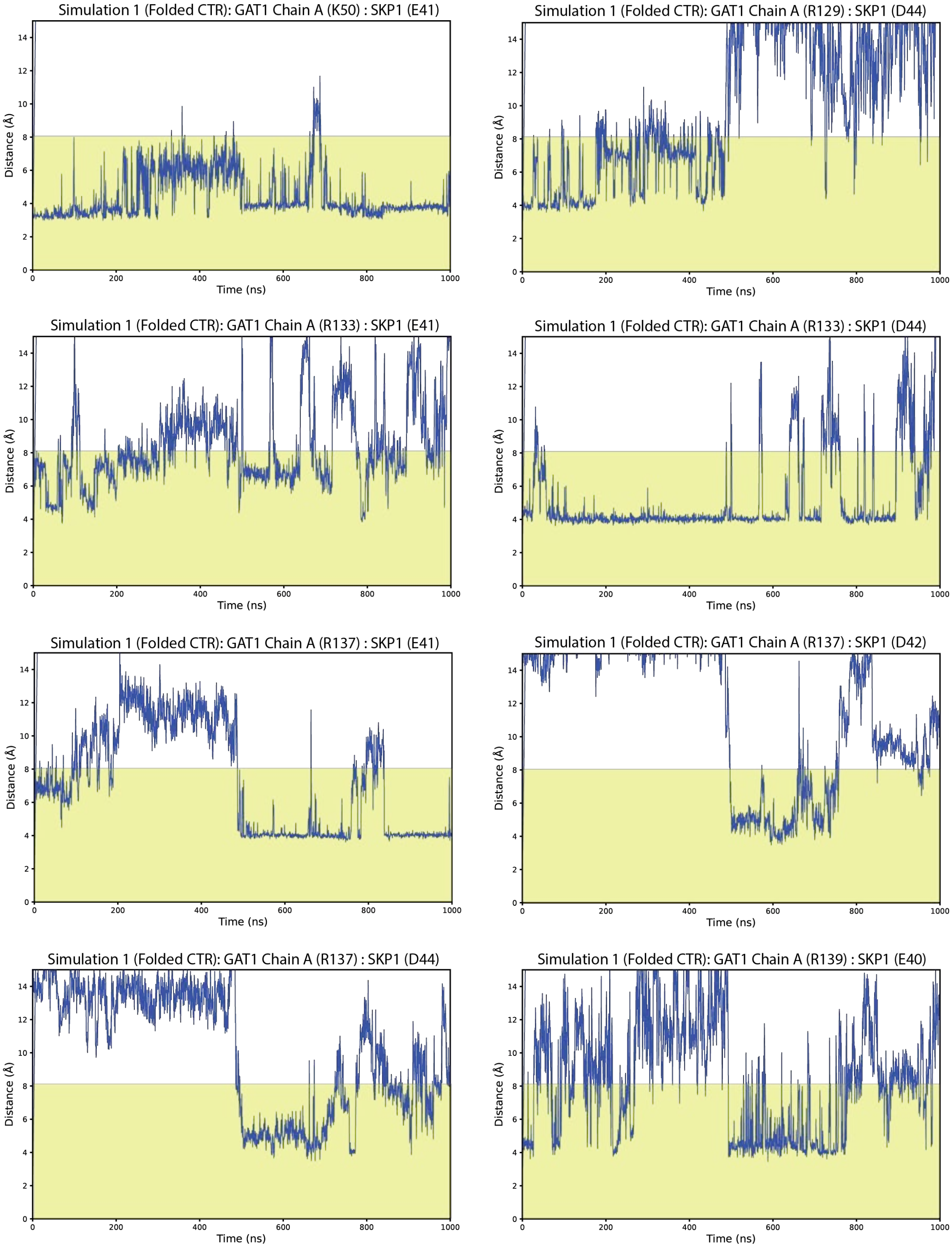

Simulation 1 (Folded CTR): GAT1 Chain A (R139) : SKP1 (D42)

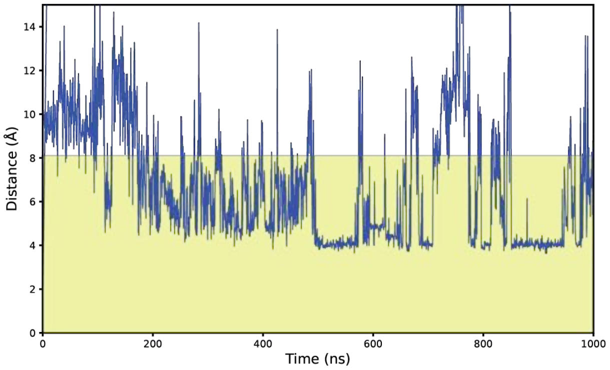


Simulation 1 (Folded CTR): GAT1 Chain A (E318) : SKP1 (K105)

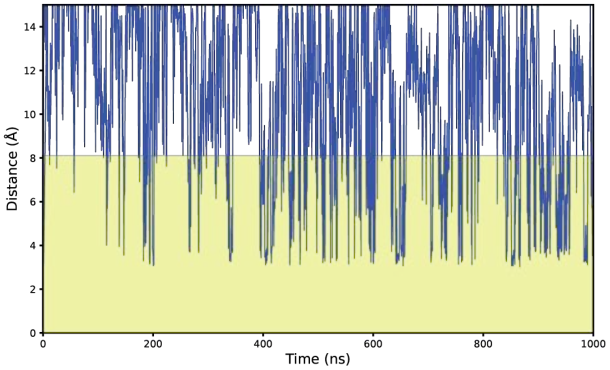


Simulation 1 (Folded CTR): GAT1 Chain A (D324) : SKP1 (K137)

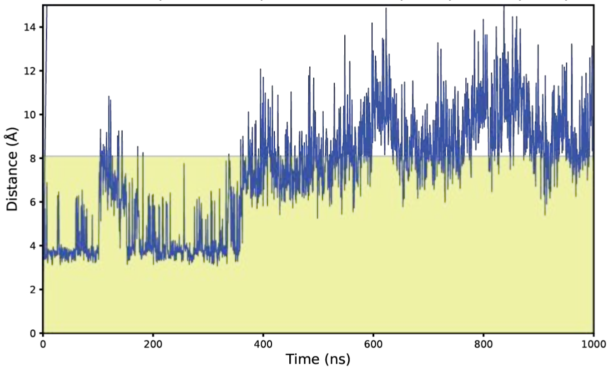


Simulation 1 (Folded CTR): GAT1 Chain A (E327) : SKP1 (K137)

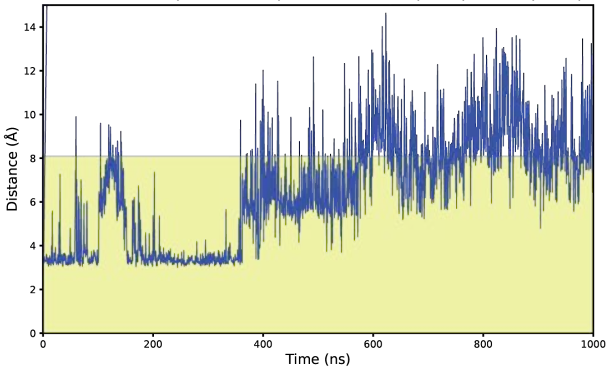


Simulation 1 (Folded CTR): GAT1 Chain A (E327) : SKP1 (R144)

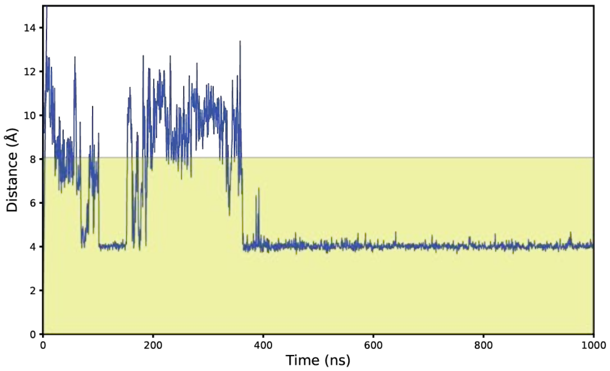


Simulation 1 (Folded CTR): GAT1 Chain A (D341) : SKP1 (R144)

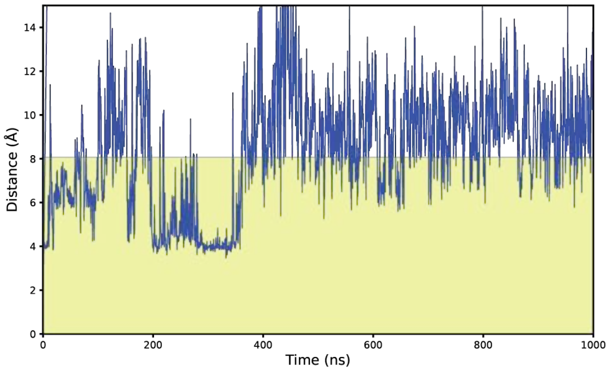


Simulation 2 (Folded CTR): GAT1 Chain A (K50) : SKP1 (E41)

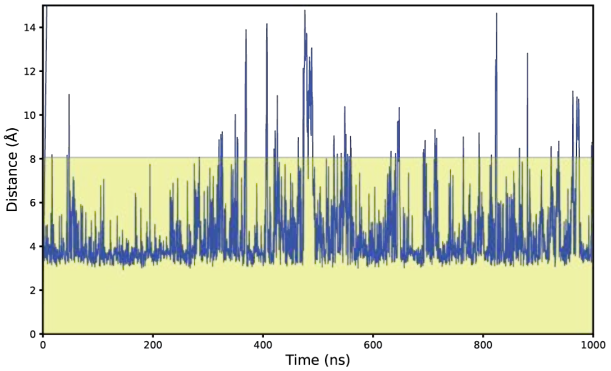


Simulation 3 (Folded CTR): GAT1 Chain A (K50) : SKP1 (E41)

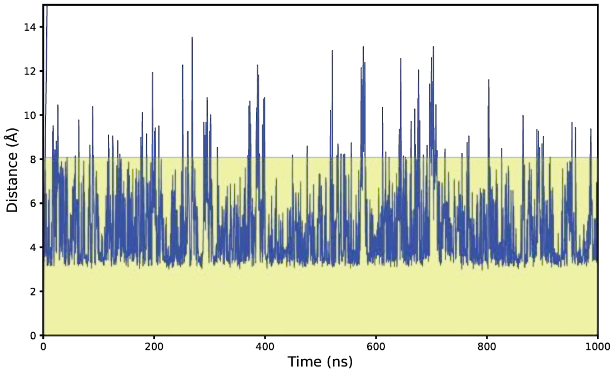

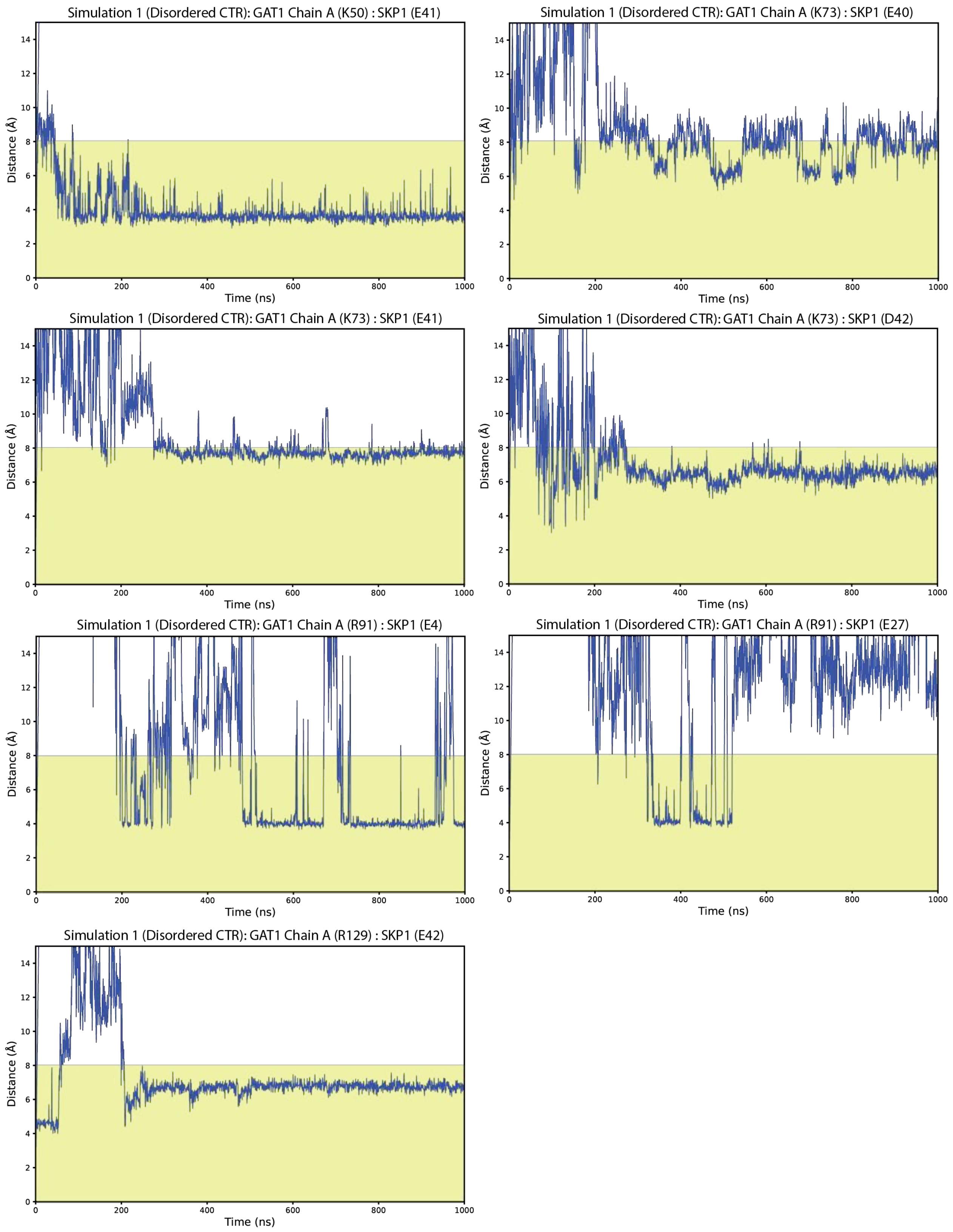
Simulation 1 (Disordered CTR): GAT1 Chain A (K50) : SKP1 (E41)
Simulation 1 (Disordered CTR): GAT1 Chain A (K73) : SKP1 (E40)
Simulation 1 (Disordered CTR): GAT1 Chain A (K73) : SKP1 (E41)
Simulation 1 (Disordered CTR): GAT1 Chain A (K73) : SKP1 (D42)
Simulation 1 (Disordered CTR): GAT1 Chain A (R91) : SKP1 (E4)
Simulation 1 (Disordered CTR): GAT1 Chain A (R91) : SKP1 (E27)
Simulation 1 (Disordered CTR): GAT1 Chain A (R129) : SKP1 (E42)
Distance (Å)
Time (ns)

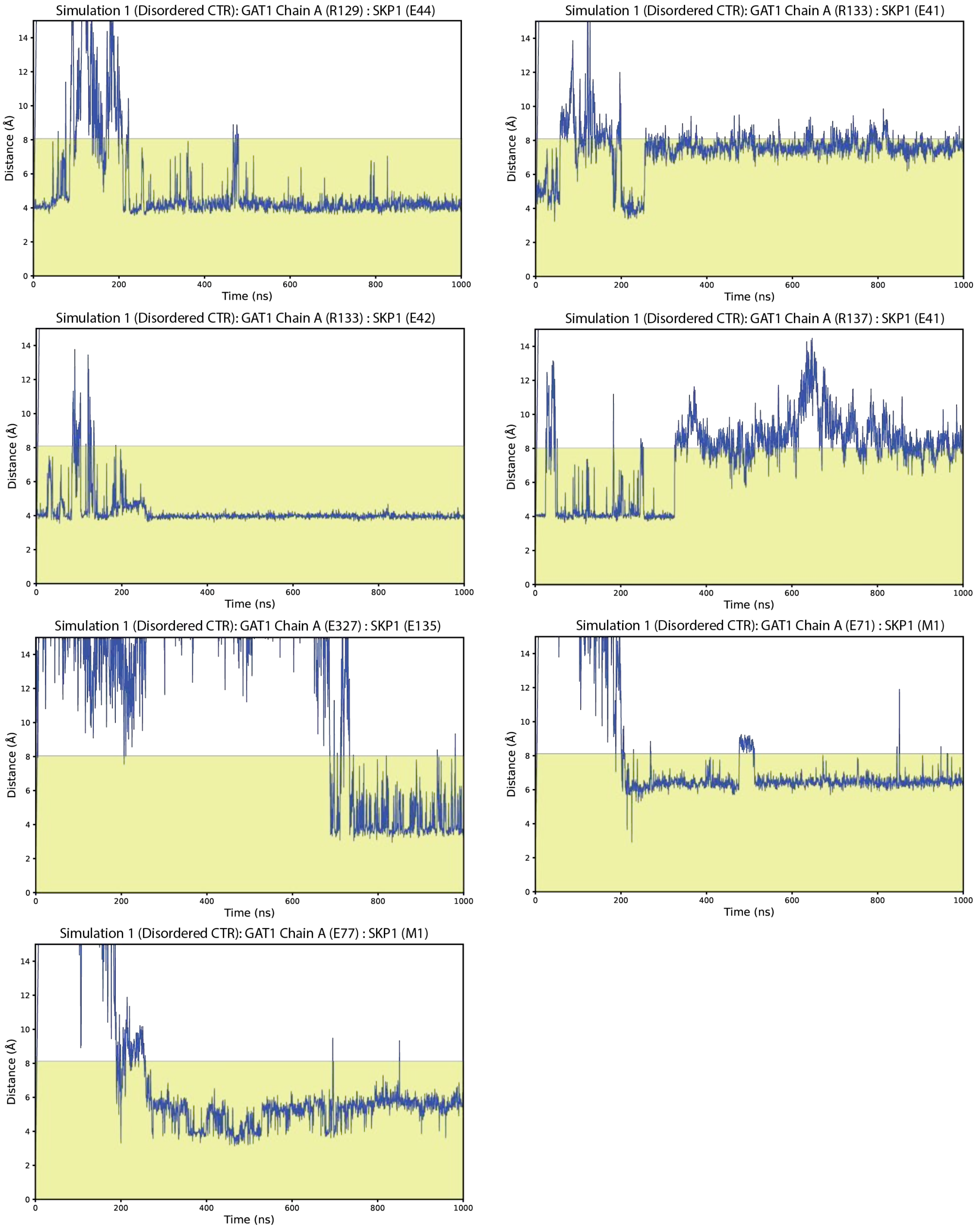

Simulation 1 (Disordered CTR): GAT1 Chain A (R129) : SKP1 (E44)
Simulation 1 (Disordered CTR): GAT1 Chain A (R133) : SKP1 (E41)
Simulation 1 (Disordered CTR): GAT1 Chain A (R133) : SKP1 (E42)
Simulation 1 (Disordered CTR): GAT1 Chain A (R137) : SKP1 (E41)
Simulation 1 (Disordered CTR): GAT1 Chain A (E327) : SKP1 (E135)
Simulation 1 (Disordered CTR): GAT1 Chain A (E71) : SKP1 (M1)
Simulation 1 (Disordered CTR): GAT1 Chain A (E77) : SKP1 (M1)
Distance (Å)
Time (ns)

**Figure S17.** Residuals and c(s) distributions for SKP1ΔCTR homodimerization with GAT1. c(s) distributions and associated sedimentation data used to model the apparent affinity of the SKP1ΔCTR homodimer with a 1:10 ratio of GAT1:SKP1 present. Regions used to generate the c(s) isotherms are indicated. Supports Fig. 8B.

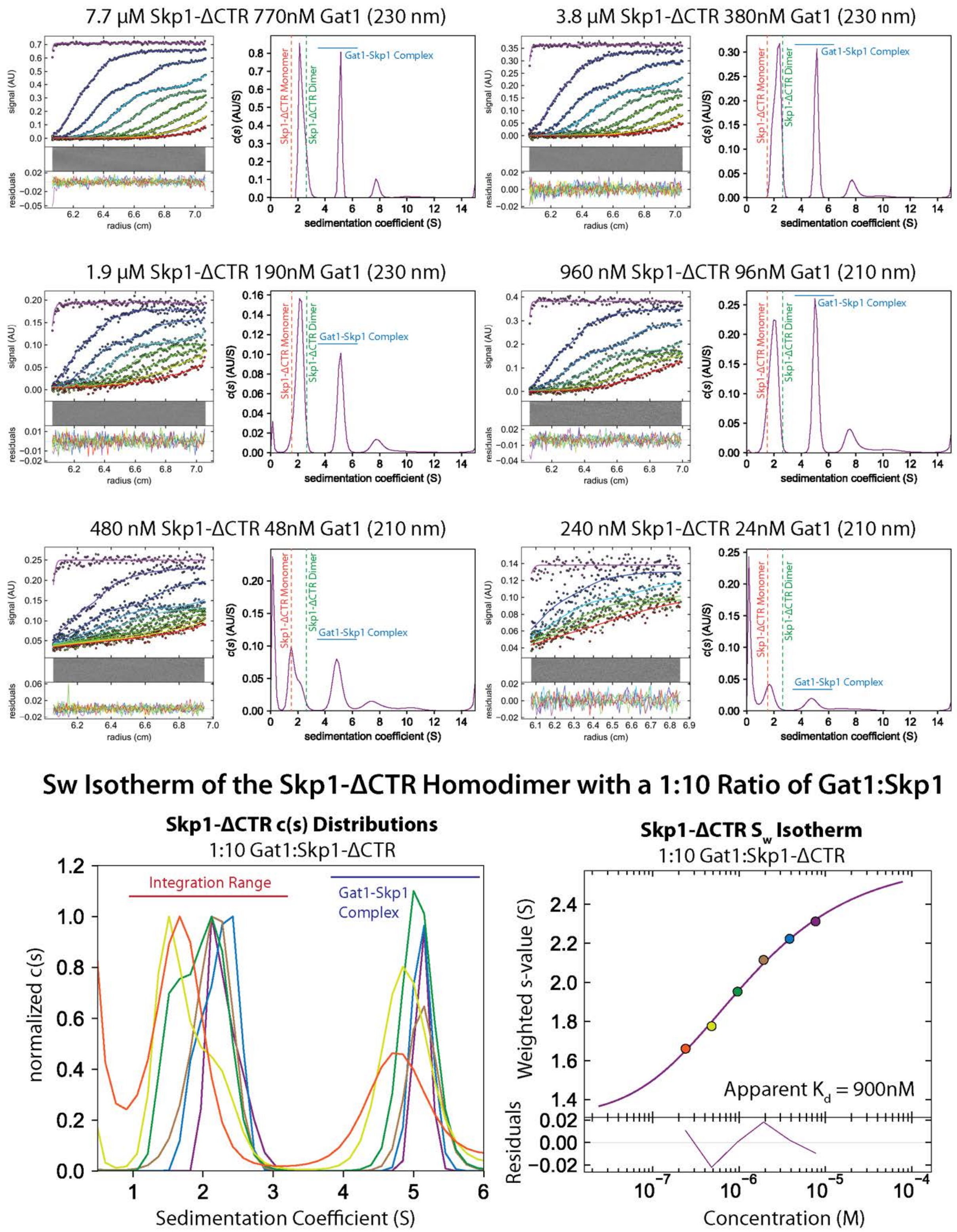

**Figure S18.** Residuals and c(s) distributions for SKP1-Scrambled6 with GAT1. c(s) distributions and associated sedimentation data used in modeling the interaction between SKP1-Scrambled6 and GAT1. Regions of the distributions used to model complex stoichiometry is indicated. Supports Fig. 8C.

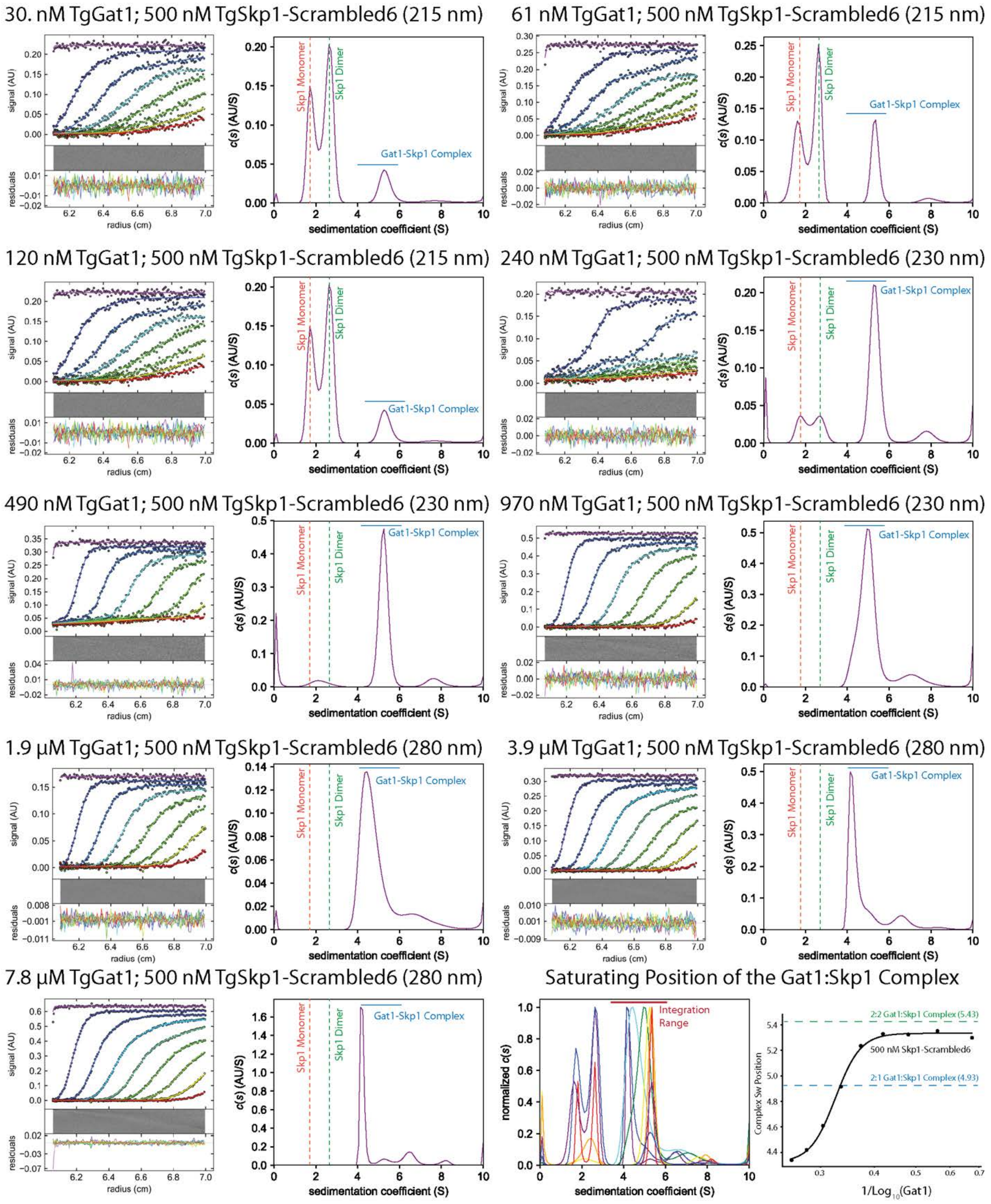

**Figure S19.** Mobility of residues in complex with glycosylated and unmodified SKP1. Simulations were initiated using identical protein structures with and without the glycan. The fold of SKP1's CTR was derived from an AlphaFold3 complex between SKP1 and FBXO13 following previous studies which suggested that glycosylation stabilizes SKP1's CTR in a folded conformation [15-17]. TASD50 mobility score comparisons for glycosylated and unmodified SKP1 simulations in the (A) GAT1 Subunit 1, (B) GAT1 Subunit 2, and (C) the SKP1 subunit are shown. Deviation mobility matrices for unmodified (D) and glycosylated (E) simulations are shown.

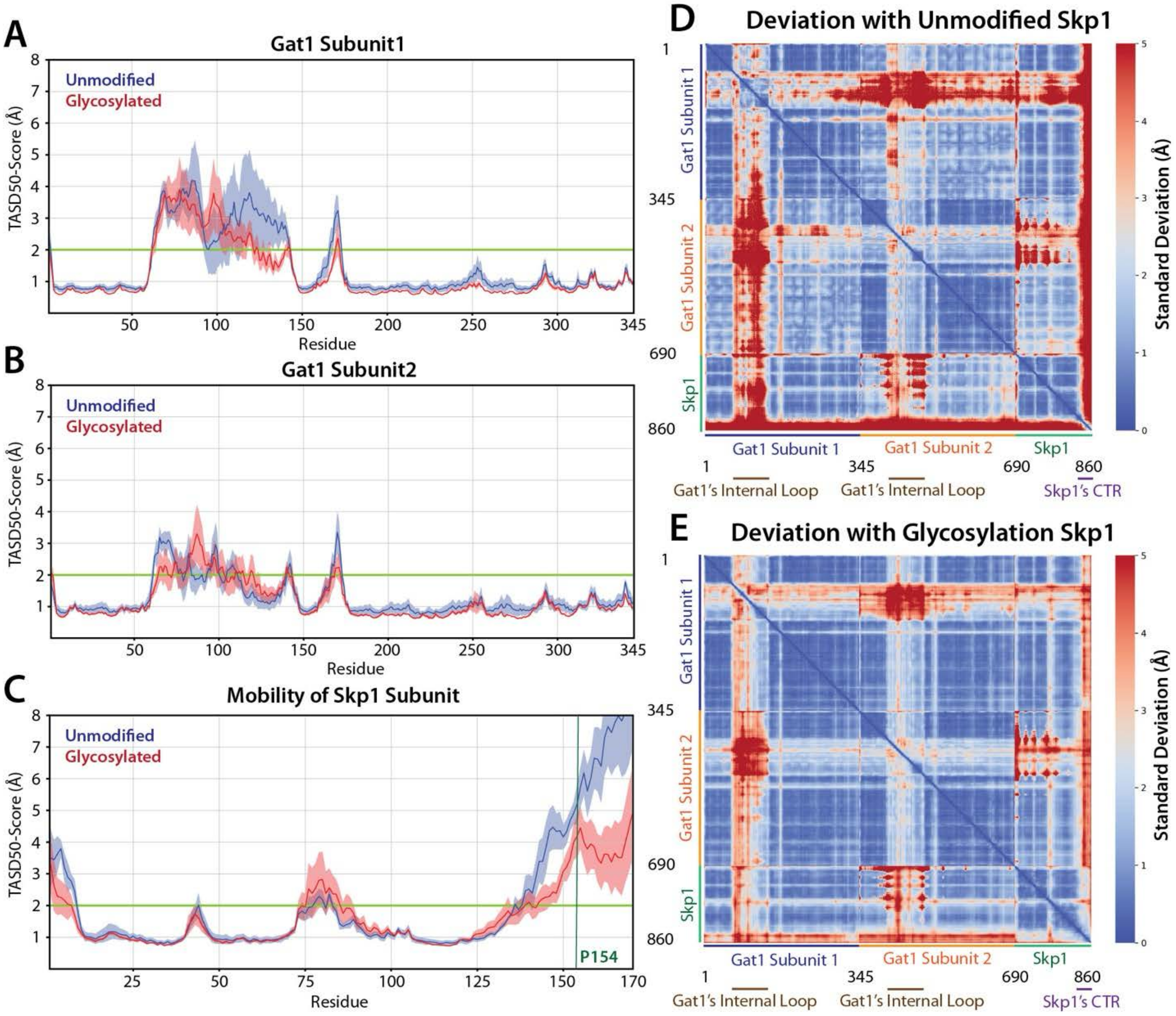
